# Global emergence and dissemination of *Neisseria gonorrhoeae* ST-9363 isolates with reduced susceptibility to azithromycin

**DOI:** 10.1101/2021.08.05.455198

**Authors:** Sandeep J. Joseph, Jesse C. Thomas, Matthew W. Schmerer, Jack Cartee, Sancta St Cyr, Karen Schlanger, Ellen N. Kersh, Brian H. Raphael, Kim M Gernert, Antimicrobial Resistant *Neisseria gonorrhoeae* Working Group.

## Abstract

*Neisseria gonorrhoeae* multi-locus sequence type (ST) 9363 genogroup isolates have been associated with reduced azithromycin susceptibility (AZM^rs^) and show evidence of clonal expansion in the U.S. Here we analyze a global collection of ST-9363 genogroup genomes to shed light on the emergence and dissemination of this strain. The global population structure of ST-9363 genogroup falls into three lineages: Basal, European, and North American; with 32 clades within all lineages. Although, ST-9363 genogroup is inferred to have originated from Asia in the mid-19^th^ century; we estimate the three modern lineages emerged from Europe in the late 1970s to early 1980s. The European lineage appears to have emerged and expanded from around 1986 to 1998, spreading into North America and Oceania in the mid-2000s with multiple introductions, along with multiple secondary reintroductions into Europe. Our results suggest two separate acquisition events of mosaic *mtrR* and *mtrR* promoter alleles: first during 2009-2011 and again during the 2012-2013 time, facilitating the clonal expansion of this genogroup with AZM^rs^ in the U.S. By tracking phylodynamic evolutionary trajectories of clades that share distinct demography as well as population-based genomic statistics, we demonstrate how recombination and selective pressures in the *mtrCDE* efflux operon granted a fitness advantage to establish ST-9363 as a successful gonococcal lineage in the U.S. and elsewhere. Although it is difficult to pinpoint the exact timing and emergence of this young genogroup, it remains critically important to continue monitoring it, as it could acquire additional resistance markers.

## Introduction

*Neisseria gonorrhoeae*, the causative agent of gonorrhea, represents a major global public health threat (Wi et al. 2017). The prevalence and incidence rates of gonorrhea are high both globally and in the U.S. In 2016, the World Health Organization (WHO) estimated there were 87 million new cases of gonorrhea worldwide (Rowley et al. 2019), with 616,392 cases reported in the U.S. in 2019 which is a 56% increase since 2015 (CDC 2021).

Since the 1940’s, *N. gonorrhoeae* has successively developed resistance to each previously recommended drug therapy for treatment and instances of gonorrhea treatment failure over the years caused by a resistant gonococcal strain have been well-documented (Unemo, Steven Seifert, et al. 2019; Unemo, Lahra, et al. 2019a). In 2013 and 2019, the Centers for Disease Control and Prevention (CDC) classified antibiotic resistant *N. gonorrhoeae* as an urgent threat in “Antibiotic Resistance Threats in the U.S.’’ (Centers for Disease Control and Prevention (U.S.) 2019). Between January 2010 and December 2020, CDC and WHO recommended dual therapy with an extended-spectrum cephalosporin ((ESC), most often ceftriaxone (CRO)), and a second drug, the macrolide azithromycin (AZM), exclusively since 2015, for the treatment of uncomplicated gonococcal infections of the cervix, urethra, and rectum as a strategy for preventing ceftriaxone resistance and treating possible coinfection with *Chlamydia trachomatis* (Workowski et al. 2010; Workowski 2015). Due to the increased incidence of AZM reduced susceptibility in gonococcal isolates, the potential impact of AZM use on susceptibility among commensal organisms and concurrent pathogens, along with the continued low incidence of reduced ceftriaxone susceptibility in previous years in the U.S., CDC, as of December 2020, recommends removal of AZM and the use of a single 500 mg intramuscular (IM) dose of CRO for treatment of uncomplicated urogenital, anorectal, and pharyngeal gonorrhea for persons weighing <150 kg (300 lb) (St. Cyr 2020).

In the U.S., Gonococcal Isolate Surveillance Project (GISP) data indicated that the percentage of isolates with reduced susceptibility to AZM (AZM^rs^; MIC ≥ 2 μ*g/ml*) was relatively low for several years, but showed a steep increase over a six-year period (0.6% in 2013 to 4.4% by 2018) (Bowen et al. 2019; Thomas et al. 2019; Reimche et al. 2021). A previous study on GISP isolates from 2000 – 2013 demonstrated that reduced susceptibilities for AZM (MIC ≥ 4 *μg/mL*) was primarily attributed to specific mutations in the 23S rRNA gene (e.g., C2611T and A2059G). More recently several studies have reported cumulative mutations in the *mtr* locus acquired by homologous recombination with commensal *Neisseria* spp. are responsible for elevated AZM MICs between 2 to 4 μg/ml in the U.S. (Grad et al. 2016; Wadsworth et al. 2018; Thomas et al. 2019; Gernert et al. 2020). These mosaic-like sequences were shown to have strong epistatic effects, increase transcription of the *mtrCDE* operon, and enhance efflux pump function (Wadsworth et al. 2018).

During 2014 – 2018, we detected an increase in prevalence of GISP isolates in the U.S. with reduced susceptibility to AZM (MIC ≥ 2μ*g/ml*), belonging to a few closely related sequence types (STs) within the multi-locus sequence typing (MLST) scheme for *Neisseria* spp. (Gernert et al. 2020; Reimche et al. 2021). This group was predominately of MLST ST-9363 among others and was recently designated as “core-genome group cluster 16” or the ST-9363-associated genome group (ST-9363 genogroup) (Harrison et al. 2020). The increasing prevalence of ST-9363 genogroup isolates with AZM^rs^ prompted us to perform an in-depth phylogenomic analysis. Here, we analyzed a global collection of 1,978 *N. gonorrhoeae* genomes belonging to the ST-9363 genogroup. These isolates were collected in the U.S. and other international locations (Canada, Europe, Oceania, and Asia) ranging over a period of 22 years (1996 – 2018). We sought to further understand the origin and expansion of this genogroup in the U.S. and worldwide, with a particular focus on the emergence and spread of antimicrobial resistance (AMR) alleles responsible for AZM^rs^ over time.

## Results

### Time-resolved phylogenetic inferences and antimicrobial resistance variants for ST-9363 genogroup

The clonal timed phylogenetic analysis of ST-9363 genogroup gonococci for a global collection of 1,978 isolates from 1996 to 2018 (Table S1) estimated a substitution rate of 4.752 × 10^−6^ substitutions per site per year, (95% highest posterior density (HPD) ranging from 4.163 × 10^−6^ to 4.933 × 10^−6^), which was similar to previous estimates from other population level studies that included gonococcal isolates from a single or similar STs (Sánchez-Busó et al. 2019; Osnes et al. 2020; Osnes et al. 2021). The time to the most recent common ancestor (tMRCA) of the entire ST-9363 genogroup was estimated to be in the mid-19th century (1853; 95% highest posterior density (HPD): [1833 - 1883]). Asian isolates were found on deeper branches of the phylogeny compared to the isolates from the rest of the world, which suggests an Asian origin for this genogroup similar to other gonococcal lineages (Sánchez-Busó et al. 2019) (Figure 1). Almost all the present day North American, European and Oceania ST-9363 genogroup isolates were estimated to have emerged from a common ancestor inferred to have existed around 1978 (95% HPD: [1974 - 1986]). In the time-resolved phylogeny, the global collection of ST-9363 genogroup isolates were broadly partitioned into three main lineages - ‘basal lineage’ (1983; 95% HPD: [1981 - 1986]), ‘European lineage’ (1990; 95% HPD: [1986 - 1998]) and ‘North American lineage’ (2005; 95% HPD: [2004-2006]) (Figure 1). By performing non-parametric tests using TreeStructure, which accounts for the underlying coalescent process in the time-scaled phylogenetic tree to determine the global population structure, 32 clades were identified (Figures 1 and 2). All major lineages and clades described were supported with > 0.70 posterior probabilities. Timed phylogenies and population trajectories estimated by modelling the effective population size over time for each of these 32 clades are shown in Figure S1.

**Figure 1.**
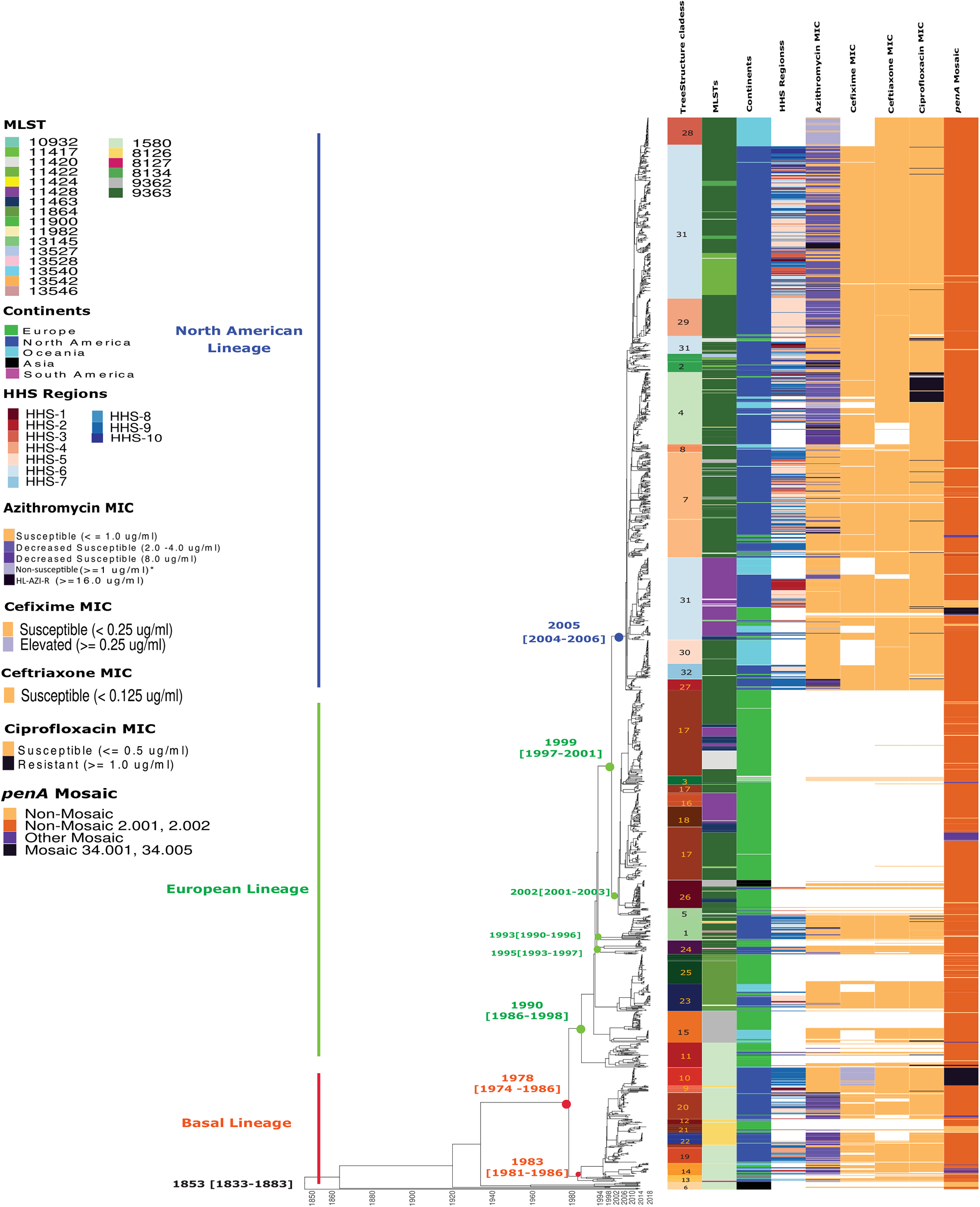
Whole genome sequence dated phylogeny with the maximum clade credibility reconstructed using BEAST annotated with TreeStructure clades. MLSTs, geographical locations and MICs of various antibiotics along with the penA allele profile of all the 1978 ST-9363 genogroup gonococcal isolates included in this study. White/blank regions in the figure indicate that the data is not available for the corresponding isolate on the phylogenetic tree. *Azithromycin MIC non-susceptible >= 1 are shown for the Oceania isolates as the exact MICs were not publicly available and in Williamson et al. (2019) all Oceania isolates with MICs >= 1μg/ml were considered as resistant to AZM, which does not conform to CLSI breakpoint. HL-AZI-R stands for “High-level Azithromycin resistance”.

**Figure 2.**
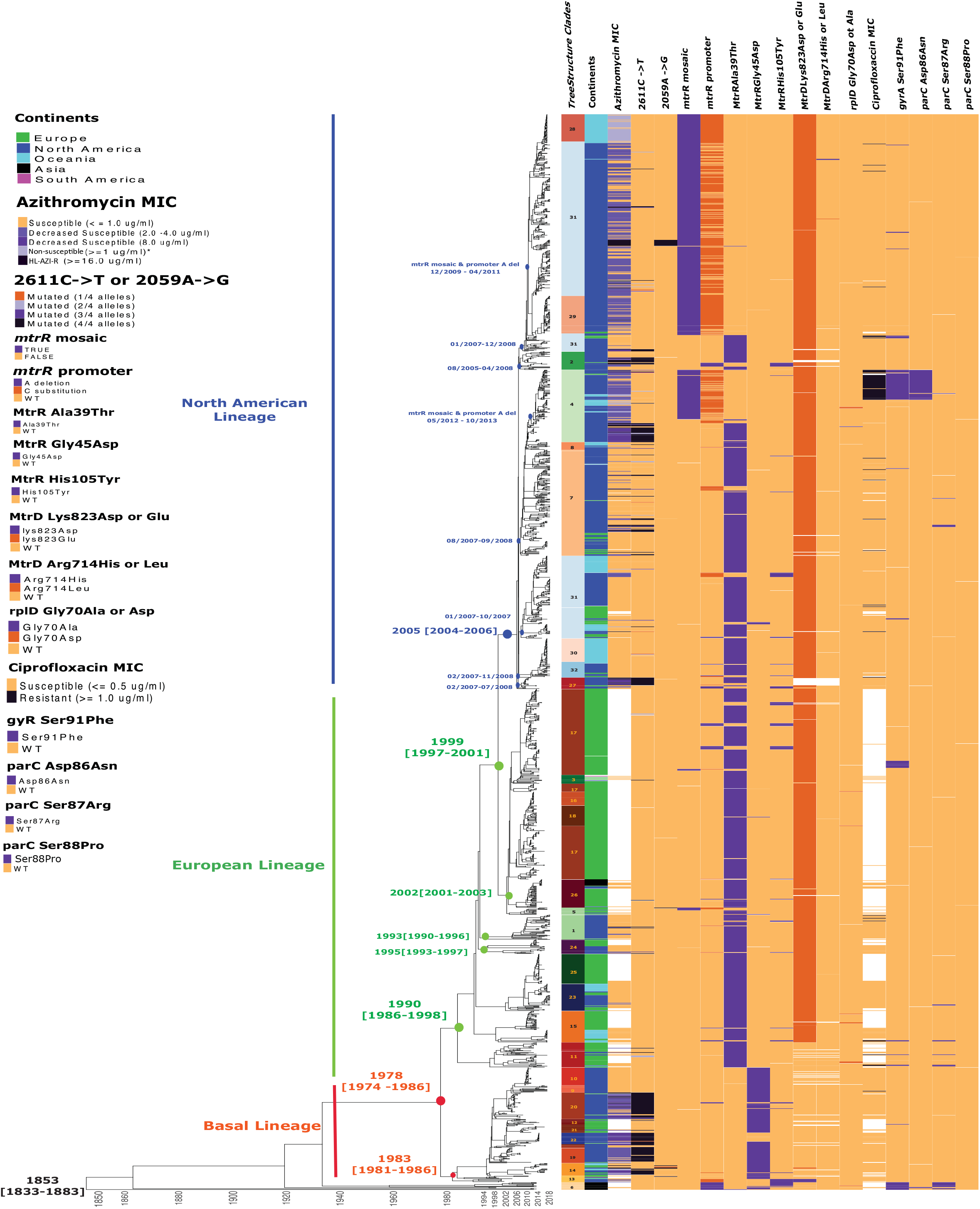
Whole genome sequence dated phylogeny with the maximum clade credibility reconstructed using BEAST annotated with TreeStructure clades, Continents, AZM MICS and the known resistance conferring genetic variants for AZI and CIP of all the 1978 ST-9363 genogroup gonococcal isolates included in this study. White/blank regions in the figure indicate that the data is not available for the corresponding isolate on the phylogenetic tree. *Azithromycin MIC non-susceptible >= 1 are shown for the Oceania isolates as the exact MICs were not publicly available and in Williamson et al. (2019) all Oceania isolates with MICs >= 1μg/ml were considered as resistant to AZM, which does not conform to CLSI breakpoint. HL-AZI-R stands for “High-level Azithromycin resistance”.

The basal lineage consisted of two MLSTs that are single locus variants of ST-9363 (MLST ST-1580 (n=157) and ST-8126 (n=52), and were predominantly from North America (78.2%), Europe (17.5%) and Oceania (3.8%), and formed nine MLST-specific and/or geographically specific clades (Figures 1 and 2). Within the basal lineage, AZM isolates (MIC ≥ 4*μg/ml*) possessing 23S rRNA mutations (primarily C2611T mutations) formed four clades (clade 14 (all 3 continents), clade 19 (Europe), clade 20 (North America) and clade 22 (North America)) (Figure 2). The majority of basal lineage isolates did not carry any of antimicrobial resistance (AMR) markers associated with low-level reduced susceptibility to AZM (*2μg/mL*), particularly those in the *mtr* locus; although 87.7% (185/211) of the isolates did possess the MtrR Gly45Asp mutation, which was associated with 0.25 to 0.5 μg/ml increase in AZI MIC, while none of the isolates carried the MtrR Ala39Thr and MtrD Lys823Glu mutations that were associated with increased MtrCDE expression as well as changes in the MtrD functionality and efflux. 91% (30/33) of the isolates clustered in the basal lineage clade 10 (n=33; North America) also possessed an elevated cefixime MIC (MIC ≥ 0.25 μg/ml) due to possession of mosaic *penA* 34.001 allele. Tracking the tMRCA for clade 10 suggested the acquisition of *penA* 34.001 allele possibly happened in the U.S between November 2008 and June 2009 (95% credibility interval) (Figure 1). We also detected a minority of isolates in the basal lineage harboring the GyrA Ser91Phe mutation responsible for conferring fluoroquinolone resistance. All 14 of the Asian isolates in the basal lineage (clade 6; n=14), carried either ParC Ser87Arg (n=11) or ParC Asp86Asn (n=2). Only one European isolate in the basal lineage (clade 12) carried the GyrA Ser91Phe mutation.

The European ST-9363 genogroup lineage (n=696; years of isolation from 2004 to 2018) consisted of predominantly MLST ST-9363 (42.1%) and its single locus variants, ST-11864 (14.7%), ST-9362 (10.1%), ST-11463 (7.5%) and ST-1580 (6.6%), while the double locus variant, ST-11428, accounted for 11.12% of the isolates. The majority of isolates in this lineage were from Europe (78.6%) followed by North America (14.3%), Oceania (5.1%) and Asia (1.7%), and they formed 12 clades (Figures 1 and 2), although with more geographical variation than the basal lineage. Interestingly, there were only nine isolates in the entire European lineage that carried either the 23S rRNA mutations C2611T (n = 8; clades 1, 3, 11, 24 and 26)) or A2059G (n=1; clade 5) responsible for higher AZM MICs (mutated at least 3 out of 4 alleles), of which five isolates were from North America and four from Europe. ST-9363 genogroup isolates carrying both the MtrR Ala39Thr (present in all European lineage clades) and MtrD Lys823Glu (also present in all European clades except clade 11) mutations, both associated with the slight decrease in AZM susceptibility (MICs still < 2μg/ml), were first observed in the European lineage. Interestingly, tMRCA of clade 11 predates the rest of the European clades indicating the acquisition of MtrR Ala39Thr occurred first in the late 1980’s to early 1990’s, while MtrD Lys823Glu was estimated to be acquired later in the mid 1990’s. The number of isolates harboring other important AZM AMR variants were very low among the ST-9363 genogroup isolates in the European lineage including *mtrR* mosaic (n=8), *mtrR* promoter alleles (A deletion (n=15); A->C substitution in the inverted repeat (IR) (n=7)), MtrR Gly45Asp (n=2), MtrR His105Tyr (n=25) and RplD Gly70Asp/Ala (n=8). A very recent report from Germany (Banhart et al. 2021) detected the presence of ST-9363 genogroup isolates in 2018, similar to the U.S. ST-9363 genogroup isolates in that they carried both *mtrR* mosaic and *mtrR* promoter alleles. However, our preliminary timed phylogenetic analysis with those German isolates showed most of those isolates clustered within the European lineage (Figure S2). Similarly, AMR markers conferring genotypic fluoroquinolone resistance, GyrA Ser91Phe in addition to either Asp95Ala or Asp95Gly (n=18), ParC Ser87Arg (n=5) and both GyrA Ser91Phe and ParC Ser87Arg (n=2) were very low among the ST-9363 genogroup isolates in the European lineage compared to other gonococcal lineages circulating in Europe.

The North American lineage (n=1057; years of isolation from 2009 to 2018) consisted of predominantly MLSTs ST-9363 (72.5%), ST-11428 (13.1%), ST-11422 (7.6%) and ST-8134 (1.6%), mostly from North America (77.3%), Oceania (15.9%) and Europe (6.8%) and was partitioned into 10 clades. There were 123 isolates in the entire North America lineage that carried either of the 23S rRNA mutations: C2611T (n=112) or A2059G (n=11) responsible for high-level AZM MICs (MICs ≥ 4 μg/ml), of which six isolates with C2611T were from Oceania and 117 were from North America (clades 2, 4, 7, 27 and 31). The signature AMR genotype associated with slight increase in AZM MICs but still susceptible to AZM (<2 μg/ml), MtrR Ala39Thr and MtrD Lys823Glu, first observed and widely present among the European isolates, was also present in 511 and 1031 North American lineage isolates, respectively. More than 41% of the North American lineage isolates harbored *mtrR* mosaic (n=495) and *mtrR* promoter alleles (A deletion (n=15); A to C substitution in the IR (n=422)), which are AMR variants identified as associated with reduced susceptibility to AZM in the U.S. (Shafer 2018; Wadsworth et al. 2018; Thomas et al. 2019; Gernert et al. 2020). Interestingly, MtrR Ala39Thr mutations were not present in the North American isolates carrying *mtrR* mosaic and *mtrR* promoter alleles; where most likely the mutation could be reverted due to homologous recombination, while the MtrD Lys823Asp/Glu mutations were intact in all the isolates. This suggests the presence of the MtrD Lys823Asp variant in the novel MtrCDE recombinant sequence, and highlights that the *mtr* locus could be a hot spot of homologous recombination among the North American ST-9363 genogroup isolates. The absence of MtrR Ala39Thr mutations in 2018 German isolates clustered within the European lineage supports the same conclusion (Figure S2). Isolates carrying other AMR markers associated with reduced AZM susceptibility were less represented in this lineage including: MtrR Gly45Asp (n=4), MtrR His105Tyr (n=26), MtrD Arg714His/Ala (n=3) and RplD Gly70Asp/Ala (n=4). AMR markers conferring genotypic fluoroquinolone resistance were carried in isolates of clade 4 [GyrA Ser91Phe and ParC Asp86Asn (n=53)] and of clade 31 [GyrA Ser91Phe) (n=11)]. Overall CIP resistance among the ST-9363 genogroup isolates was low at 6.3% (67/1057) compared to other MLSTs circulating in North American/USA (31.2%) (Anon 2021). Also, approximately 90% of the ST-9363 genogroup isolates from the U.S. GISP 2018 carry PorB Gly120Asp/Lys and PorB Gly121Asp/Asn mutations (data not shown), conferring low-level tetracycline (TET) resistance (with MIC = 2.0 – 4.0 μg/ml).

### Geographic dispersal patterns of ST-9363 genogroup

From the timed phylogeny (Figure 1), it appears that the ancestor of the currently observed ST-9363 genogroup emerged in Asia in the mid-19th century (Figure 2). Ancestral state reconstruction conducted (excluding all the Asian and South American isolates due to very low number of isolates) to understand the spatio-temporal dispersal history and geographical origins of this genogroup suggested a European origin for the 20th century ST-9363 genogroup isolates with an estimated posterior probability of 86.5%, while the support for a North American origin was only 13.4% (Figure 3). The down-sampled phylogeographic analysis also suggested a European (79.1%) rather than a North American (20.9%) origin (Figure S3). The global transmission trends inferred from the medoid tree using TransPhylo also suggested a European origin of the current ST-9363 genogroup isolates circulating in North America (Figure S4).

**Figure 3.**
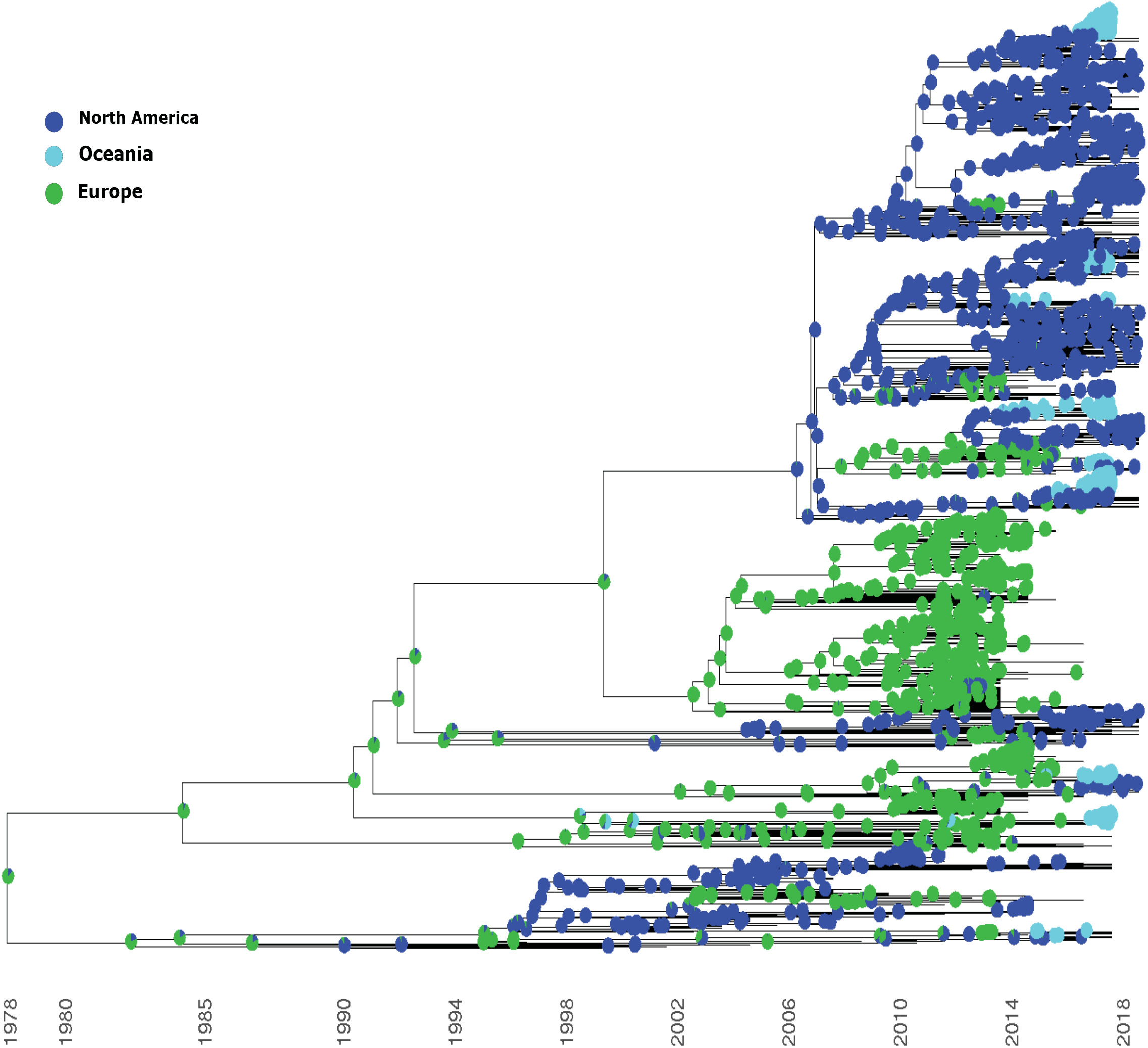
Ancestral state reconstruction based on the maximum clade credibility reconstructed using BEAST with geographical locations (Continents) as discrete traits. Each node indicates the estimated posterior probabilities of each of the discrete traits - Europe, North American and Oceania as pie charts.

The basal lineage, which had a deep split from the common ancestor of all the 20th century ST-9363 genogroup isolates in the phylogeny and also was separated from the European and North American lineages, was inferred to have emerged between December 1981 and April 1986 (95% credibility interval) (Figure 2). The observed phylogeographical patterns suggested that the deeper ancestral nodes in this lineage were mapped to Europe with high confidence (83.2% posterior probability; Figure 3), and the North American isolates were descendants of more than one expansion out of Europe in the early 1990s, which were again probably reintroduced back into Europe and Oceania in the 2000’s and 2010’s, respectively, from North America (Figure 3). There was no evidence of onward dissemination of 3 out of the 4 basal clades (clades 19, 20, 22) that acquired 23S rRNA mutations (C2611T) conferring reduced susceptibility to AZM; the effective population size (EPS) was declining for the last 6 to 10 years for these three clades. Clade 14 with a fraction of strains with C2611T variants and a fraction with A2059G variants, was the exception, with an EPS increasing in 2016-2017 (Figure S1). Even though clade 10 isolates that harbored *penA* mosaic 34.001 alleles with an elevated cefixime MIC ≥ 0.25 might have been circulating in the western states of the U.S. (HHS-9), tracing its phylogenomic trajectory indicated the inferred EPS peaked in 2010 and then remained constant with no evidence of ongoing dissemination after 2011 (Figure S1).

The present-day ST-9363 genogroup isolates that belonged to European and North American lineages appeared to have originated and expanded from a common ancestor inferred to have existed approximately around 1990 (95% HPD: 1986 to 1998) (Figure 2). During the 1990s there were multiple strains, many of which were single locus variants of ST-9363, circulating in Europe. These were eventually introduced into North America and Oceania; examples include: clades 11 (ST-1580), 15 (ST-9362), 23 (ST-11864) and 25 (ST-11864) (Figure 1). Initially there were two independent introductions of ST-9363 isolates into Europe. The first introduction appeared to be around 1993 (CI: 1990 -1996; clade 1), which contained mostly North American isolates (all from the U.S.; years of isolation from 2013 to 2018), and not only included ST-9363 isolates but also its single locus variants, ST-11982, ST-11422, ST-13546, ST-13542 and ST-13527. The second introduction of exclusively ST-9363 isolates possibly happened around 1995 (CI: 1993 to 1997; clade 24) with isolates from North America (U.S. (n=12); Canada (n=2)) and Europe (n=12) with years of isolation ranging from 2009 to 2016. Tracing the population trajectories for these 2 clades indicated a decline in the inferred EPS for the last 5 years. In Europe, a third independent expansion of ST-9363 isolates started in 2002 (CI: 2001 to 2003), from this point on, this lineage rapidly spread across Europe forming six clades; either by multiple introductions or acquiring mutations and evolving into other closely related variants like ST-11428 and ST-11463.

The North American lineage and the third expanding European ST-9363 clades were inferred to have diverged sometime around 1997 to 2001 (95% CI) and based on the phylogeographic analysis this transition from Europe to North America was supported with 99.8% posterior probability towards a European origin (Figure 3). The tMRCA from where the recent North American lineage emerged was estimated to be around June 2004 and September 2006 (95% CI). There were at least six independent introductions of the ST-9363 genogroup strains into North America between August 2005 and December 2008, all clustered as monophyletic branches and were either disseminated from Europe or descendants of unsampled strains circulating within North America during the same time period. Out of six introductions, two (clades 7, 8 and 4 inferred to be emerged between August 2007 and September 2008; and clades 28, 29 and 31 emerged around January 2007 and December 2008) appeared to be successful introductions based on their clonal expansions in North America and Oceania with a few reintroductions into Europe. Even though initially these two sub-lineages contained similar AMR profiles as the European ST-9363 genogroup isolates, which carried both the MtrR Ala39Thr and MtrD Lys823Glu mutations; later on, both sub-lineages separately acquired the *mtrR* mosaic allele and promoter A -> C genotype. The first acquisition of this new AMR genotype was inferred to happen between December 2009 and April 2011 (Clade 28, 29, 31); while the second acquisition happened around May 2012 and October 2013 (Clade 4). Tracing the population trajectories for the 10 clades indicated a decline in the inferred EPS for only two clades (clades 8 and 27) in the last 2 to 8 years since 2018; while the EPS for the rest of the clades were inferred to be expanding.

To ensure the above evolutionary estimates were not overly shaped by biased sampling, we repeated the analyses on a down-sampled dataset that contained only isolates from North America, Europe and Oceania (see Materials and Methods). The inferences from this dataset largely restate our estimates on the overall population structure and the origins of the European and North American lineages; but due to the exclusion of all the Asian isolates estimates for tMRCA for all the 20^th^ century isolates and the basal lineage appeared to be a little earlier than the estimates with the full dataset (Figure S3; Table S2 (a) and S2(b)).

### Recombination and its role in the clonal expansion of ST-9363 genogroup

The mean ρ/θ and r/m values estimated for the entire ST-9363 genogroup were 0.127402 and 3.1342, respectively. The ρ/θ estimate is slightly lower than previous estimates for the entire *N. gonorrhoeae* species (ρ/θ = 0.41) (Vigué and Eyre-Walker 2019) while the r/m estimate was higher than previously reported species level ratios (2.0 (Vigué and Eyre-Walker 2019) and 2.2 (Ezewudo et al.)). Higher r/m estimates occur when very closely related strains were assessed, similar to the isolates analyzed in this study, as natural selection might not have a chance yet to remove the deleterious mutations introduced by recombination as in distantly related strains within a species (Vigué and Eyre-Walker 2019). r/m estimates are more appropriate for species-level recombination summaries as it indicates how important the effect of recombination was in the diversification of the bacterial species relative to mutation, whereas ρ/θ measures how often recombination events happen relative to mutation. We decided to further analyze the ρ/θ estimates for each clade to understand whether the rates of recombination were different across the three lineages as well as among the 32 clades (Table S3).

The distribution of ρ/θ estimates of isolates in each clade is shown in Figure S5 as well as the per clade average ρ/θ estimates are shown in Table S3. The Kruskal-Wallis rank sum test indicated that ρ/θ was significantly different at least among any two lineages (p-value = 6.081e^−06^) as well as among any two clades (p-value = 2.494e^−13^); suggesting that across lineages as well as clades within lineages must be evolving differently in terms of their recombination rates. The *post hoc* pairwise statistical testing using Dunn’s test revealed that the rates of recombination were significantly different between basal versus European lineages comparison (adj p-value = 2.1564e^−02^) and European versus North American lineages (adj p-value = 3.3638e^−06^); while the comparison between the basal and North American lineages were not statistically significant different (adj p-value = 0.054). Pairwise comparison of ρ/θ estimates among clades indicated that the recombination rates among 18 clade pairs were significantly different (adj p-value < 0.05; Table S4; Figure S5). Interestingly the estimated recombination rates for the North American clades carrying the *mtrR* mosaic and promoter alleles were significantly different from those estimated for the European clades with no mosaic *mtrR* and *mtr* promoter alleles. For example, ρ/θ estimates for North American clades with *mtrR* mosaic and *mtr* promoter alleles (isolates mostly with AZI MICs > 2 *μg/mL*) such as clade_31_mosaic_mtrR were significantly different compared to clade 17 (adj p-value = 0.0116: European), clade 23 (adj p-value =0.0223; European) and clade 30 (adj p-value = 0.01505; North American) that did not carry those AMR markers (Table S4).

Within clade recombination hotspots across the genome for three clades are summarized in Figure S6 (clade_31_mosaic_mtrR; North American), Figure S7 (clade 4; North American) and Figure S8 (clade 17; European). The majority of those events occurred in the internal branches (i.e., ancestral recombination events), which were present in multiple isolates and were shared through clonal descent rather than independent acquisitions. The number of recombination events in the terminal branches that are isolate-specific and represent independent recent acquisitions were less frequent. There were a number of genes frequently present within these hotspots across these clades, which include genes that were also found to be under recombination in previous *N. gonorrhoeae* species-level analyses, including glycoside hydrolase family protein, phage associated proteins, the *mtr* efflux pump genes, 50S ribosomal protein L3 (rplC) and ABC transporter permease (feuB) (Wadsworth et al. 2018; Ezewudo et al.); while the frequency of the recombination events in these genes varied across clades. Although a couple of North American ST-9363 genogroup clades recently disseminated successfully in the population, we were not able to locate a particular recombination event/hotspot in its chromosomes that appears to be exceptional on a macroscopic scale compared to other clades with the exception of enriched presence of mosaic *mtrR* and *mtrR* promoter alleles.

Recent studies have reported an association between mosaic *mtr* efflux pump alleles and increased resistance to AZM (Trembizki et al. 2014; Demczuk et al. 2016; Grad et al. 2016; Rouquette-Loughlin et al. 2018; Whiley et al. 2018; Thomas et al. 2019; Gernert et al. 2020). Wadsworth et al (2018), using genome sequence-derived population genetic statistics along with genetic transformation analyses of the *mtr* efflux pump genes, revealed that the presence of mosaic-like alleles in the *mtrR* and *mtr* promoter and *mtrCDE* regions increased transcription of MtrCDE and MtrD function, which facilitates increased expulsion of AZM resulting in high AZM MICs. Similarly, we also attempted to understand the impact of higher recombination rates at the *mtr* efflux pump genes in some important ST-9363 North American clades (clade 31 (Figure S9(a), clade 4 (Figure S9(b), clade_31_mosaic_mtrR (Figure S9(c) and clade 29 (Figure S9(d)) and European clades (clade 17 (Figure S9(e), clade 23 (Figure S9(f), clade 24 (Figure S9(g) and clade 1 (Figure S9(h)) by assessing patterns of allelic diversity for the whole genome, the entire *mtr* locus genes, *mtrD* gene and the *mtrR* plus *mtr* promoter region in each clade. For the European clades, Tajima’s D statistics were negative across the *mtrRCDE* locus, which indicate an excess of low frequency alleles in this region introduced by recombination and were maintained by positive selection, while the linkage disequilibrium was strong at the *mtrD*, the *mtrR* plus *mtr* promoter regions, along with the entire *mtrRCDE* region suggesting that the high linkage within the *mtr* locus recombination hotspots did not have sufficient evolutionary time to break down local associations and that the local elevation of rare mutations introduced by recombination has not fixed in the population, yet.

The North American clade 31 (with no mosaic *mtrR*) has similar trends of Tajima’s D and linkage disequilibrium (LD) as the European clades. For isolates in clades 31 (with mosaic *mtrR* and *mtrR* promoter mutation) and 29, with an estimated acquisition of *mtrR* mosaic alleles sometime between December 2009 and April 2011 (Figure 2), Tajima’s D estimates for *mtrD* were negative while for the *mtrR* plus *mtr* promoter region Tajima’s D estimates were zero indicative of a neutral evolution of this allele. LD was also zero at the *mtrR* plus *mtr* promoter regions in these two north American clades suggesting the mosaic *mtrR* and *mtr* promoter alleles were possibly fixed in the population, and new rare alleles are only being introduced in the *mtrD* regions largely by recombination and/or by point mutations. For the North American clade 4, with an estimated acquisition of *mtrR* mosaic alleles after 2012, Tajima’s D for the entire *mtrRCDE* and *mtrD* were positive, while for the *mtrR* plus *mtr* promoter region Tajima’s D estimate was negative but trending positive suggesting that clade 4 might have hit a population bottleneck. LD in clade 4 still exists in the *mtrRCDE* region suggesting that the mosaic *mtrR* and *mtr* promoter alleles are still not fixed in the population (Figure 4).

**Figure 4.**
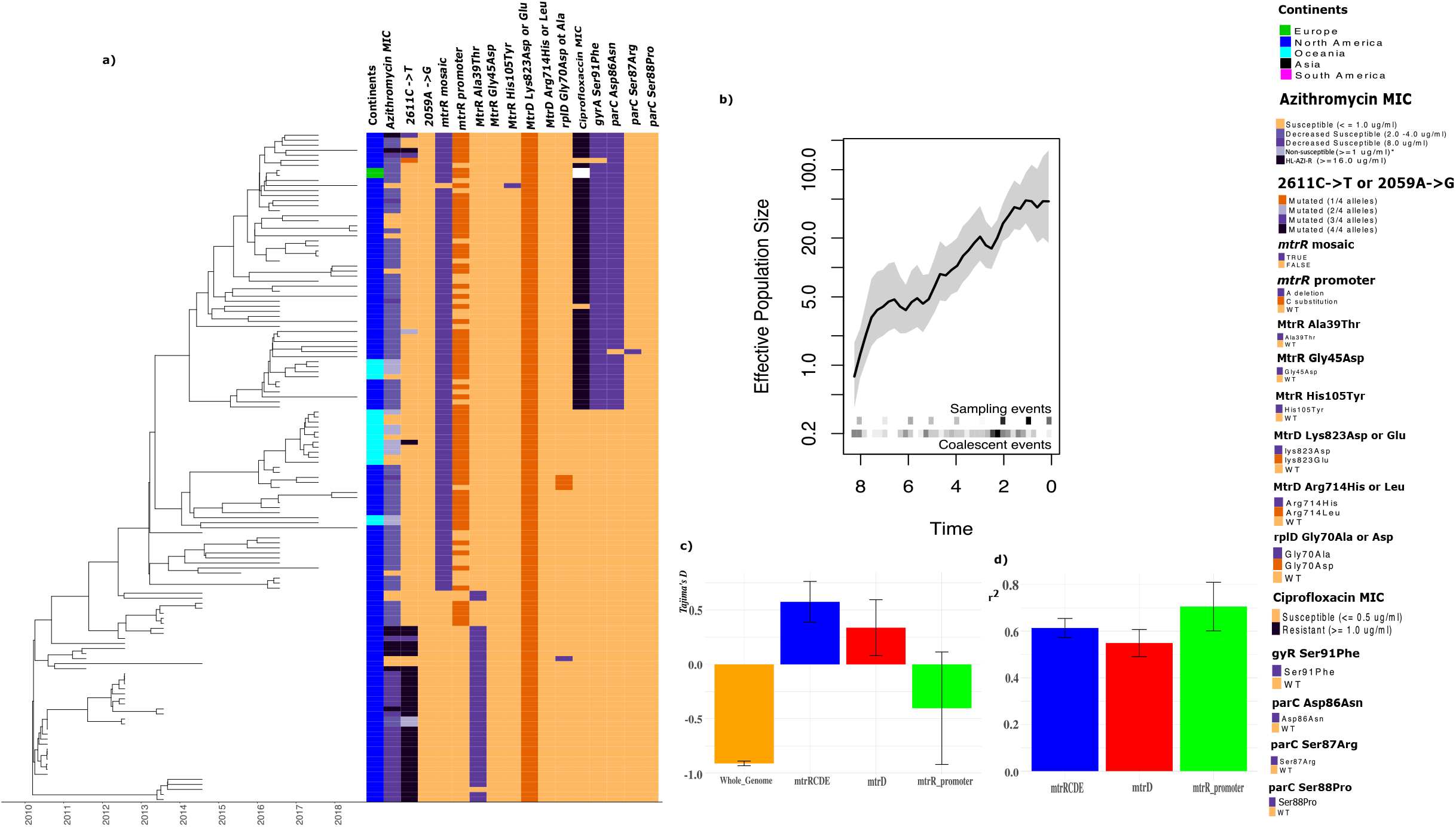
a) Whole genome sequence dated phylogeny of Clade 4 isolates (North American lineage) along with the known resistance conferring genetic variants for AZI and CIP of all the clade 4 ST-9363 genogroup gonococcal isolates. b) Demographic history of clade 4. Time is measured in years and 0 indicates the year 2018 (x-axis) and the effective population size is scaled to the number of generations per year. c) Bar diagram showing the estimated Tajima’s D statistics estimated for the whole genome, the entire *mtr* locus genes, *mtrD* gene and the *mtrR* and promoter region for the clade 4 isolates. d) Estimated linkage disequilibrium (LD) measured by r^2^ for the entire *mtr* locus, *mtrD* gene and the *mtrR* and promoter region for the clade 4 isolates. Both Tajima’s D and r^2^ were estimated over a 100 bp sliding window on the entire whole genome alignment containing only the clade 4 isolates using the R package called PopGenome. *Azithromycin MIC non-susceptible >= 1 are shown for the Oceania isolates as the exact MICs were not publicly available and in Williamson et al. (2019) all Oceania isolates with MICs >= 1μg/ml were considered as resistant to AZM, which does not conform to CLSI breakpoint. HL-AZI-R stands for “High-level Azithromycin resistance”.

## Discussion

An in-depth analysis of *N. gonorrhoeae* ST-9363 genogroup isolates from the U.S and international locations (Canada, Europe, Oceania, South America and Asia) over a period of 22 years (1996-2018) has provided some insights on the emergence and evolution of this lineage with concerning levels of decreased AZM susceptibility, and evidently having a recent and continued clonal expansion in the U.S. since the early 2010s. Even though our timed phylogenetic tree indicated that this genogroup was inferred to have originated in Asia in the mid-19th century, solely based on the phylogenetic placement and inferred tMRCA of the few modern Asian isolates used in this study; phylogeographic stochastic mapping and transmission modelling suggested the current global ST-9363 genogroup isolates have originated from Europe in the late 1970’s and spread from there to North America (U.S and Canada) and Oceania with multiple introductions, along with multiple secondary reintroductions into Europe. The North American lineage appeared to have emerged sometime between 2004 and 2006 and, interestingly, the genotypic profiles of earlier North American isolates were very similar to that of the European isolates. Our analysis indicated two separate acquisitions of mosaic *mtrR* and promoter alleles in the *mtr* locus in the late 2000’s and early 2010’s that might have steered decreased AZM susceptibility, which probably led to the recent clonal expansion of ST-9363 genogroup in the U.S. Our phylogenetic comparison identified highly similar ST-9363 isolates with mosaic *mtr* locus from Australia in 2017 (Williamson et al. 2019) and Canada (Demczuk et al. 2016). Most recently a report from Germany (February 2021) (Banhart et al. 2021) identified highly similar isolates with the same strain profile as in the U.S in 2018, supporting an international clonal expansion of this lineage. But with a relative lack of recently sampled genomes from other geographical areas other than the U.S. and lack of genomes from Africa and South America, it is difficult to gauge the spread of this genogroup worldwide. For example, our additional timed phylogenetic analysis comparing very recently available 2018 German ST-9363 genogroup isolates against the global collection indicated that the majority of those isolates with mosaic *mtrR* and *mtrR* promoter alleles clustered within the European lineage suggesting that these German strains probably do not have a U.S origin; rather they might have acquired these AMR genotypes independently. In fact, this suggests that the low number of European ST-9363 isolates with mosaic *mtrR* locus could be biased due to the lack of genome sequences for recent isolates from Europe.

Obtaining representative samples for the global ST-9363 gonococcal population is often a challenge. Convenient sampling by incorporating all available unique genomes from a genogroup along with the use of robust evolutionary models similar to this study and other genome-based gonococcal evolutionary analysis (Osnes et al. 2020; Osnes et al. 2021) are appropriate to estimate global evolutionary trends, although it might be inadequate for estimating incidence or prevalence of these strains. Our analyses performed on a down-sampled geographically balanced dataset largely identified the same evolutionary trends similar to the findings inferred using the full dataset. The credibility intervals for the tMRCA for both European and North American lineages overlapped with our original estimates, while tMRCA for all the 20^th^ century isolates and the basal lineage appeared to be earlier. This happened mainly due to the exclusion of all Asian isolates in the down-sampled dataset, which led to the loss of informative sites as well as reduced temporal signal (Root-to-tip R^2^ = 0.35). It has been previously shown that most of the modern gonococcal lineages circulating in the world emerged from Asia (Sánchez-Busó et al. 2019), and with just 26 Asian ST-9363 genogroup genomes our full dataset correctly estimated the Asian origin. Therefore, we believe the inferences from the full dataset are more robust compared to the down-sampled dataset in this study, but a globally representative dataset would have made much more reliable evolutionary inferences.

The fully resolved, timed phylogenetic analysis (using evolutionary “molecular clock” models that estimated the coalescent events occurred over time) was ideal for understanding the evolution of ST-9363 genogroup as these isolates were closely related strains due to the recent clonal expansions. This enabled us to effectively translate the branch lengths to time points, which essentially provided a better phylogenetic resolution for evolutionary inferences. Moreover, timed phylogenetic analysis allowed us to find clades with a similar history of effective population size changes using the TreeStructure algorithm, which are essentially partitions likely to share distinct demographic or epidemiological histories and even clades with beneficial AMR mutations that expanded in the recent past (Volz et al. 2020). Also, clades detected by this method enabled us to further characterize them using phylodynamic methods to track their population trajectories (Retchless et al. 2021). For example, clade 31 in the North American lineage (Figures 1 & 2) appeared to be polyphyletic, suggesting possible multiple importations of a particular strain in a specific geographic area or population such as men having sex with men (MSM) and men having sex with men and women (MSMW). The initial two introductions of clade 31 strain were estimated to be during the 2007-2008 period with only 16.85% (30/178) isolates reported sexual orientation as MSM/MSMW. The third introduction happened later in the early to mid-2010s and identified concurrently with the acquisition of mosaic *mtrR* allele and promoter A > C allele; and this partition contained isolates predominantly from the MSM population (73.26%, 211/288) including MSM/MSMW (Table S1). Effective population size of the entire clade 31 isolates showed an increasing trend until 2016 suggesting a higher rate of transmission events among the MSM population facilitating acquisition of beneficial AMR variants by sustained recombination events especially at the *mtr* locus. This occurrence could have led to the selection and fixation of mosaic *mtrR* operon that increases the expression and function of efflux pump to effectively remove AZM and other antibacterials. Another polyphyletic European clade detected by TreeStructure was clade 17, which appeared to have emerged from an ancestor that existed sometime between 2001-2003. It eventually surfaced as a clonally expanding clade in Europe. With the most recent isolate sampled in this clade being from the year 2016, the estimated effective population size of this clade was expanding until 2014, and after that it started to decline. Without much of the demographic data available for clade 17 we cannot make any further inferences for this clade. Similarly, the modeling parameters inferred from the global transmission analysis had a better fit using the parameters derived from a modelling study in a community of MSM (prior 1 was the best fit model parameter for the different assumptions tested on sampling densities (Figures S9(a) and S9(b)); this suggests MSM populations might be drivers for shaping the current population structure of this genogroup. Also, the genome-based basic reproductive number (R_0_ = 1.09 (credibility interval: 1.07 - 1.11)) for ST-9363 genogroup estimated by the global transmission analysis suggests that the number of secondary cases of disease caused by a single infected individual are increasing, which could occur when STIs are introduced in specific sexual activity populations such as MSM populations (Jolly and Wylie 2002; Ito et al. 2019).

Even though 35.4% of the U.S. (2019) (CDC 2021) and > 60% of the global gonococcal isolates (2018) were resistant to CIP (Unemo, Lahra, et al. 2019b), ST-9363 genogroup included only an estimated 5% of isolates with either higher CIP MICs (>=1.0 μg/ml) or carrying a GyrA Ser91Phe in addition to the ParC Asp86Asn, ParC Ser87Arg and/or ParC Ser88Pro mutations. In other gonococcal lineages like MLST-1901 fluoroquinolone resistance was found to be universal (Osnes et al. 2021). The low incidence of CIP resistance among ST-9363 genogroup isolates is likely due to the recent emergence of this lineage. Interestingly, during the 2016-2018 period, 53 North American, clade 4, ST-9363 isolates picked up both GyrA Ser91Phe and ParC Asp86Asn mutations conferring high levels of CIP resistance. This resistance has emerged already on a genomic background with mosaic *mtr* operon associated with elevated AZI MICs. Kunz et al 2012 (Kunz et al. 2012) reported that GyrA Ser91Phe and GyrA Asp95Asn conferred a fitness benefit in gonococci by conferring intermediate fluoroquinolone resistance *in vivo;* while an additional ParC Asp86Asn mutation could give high levels of fluoroquinolone resistance but the fitness benefit will be curbed. However, they also observed the emergence of compensatory mutations in the *mtrR* gene and *mtr* promoter region to restore the fitness benefit (Dillon and Parti 2012). Similarly, ST-9363 clade 4 isolates might have hit a bottleneck after the acquisition of both the GyrA and ParC mutations, which can be inferred from (1) the “leveling off” of the EPS estimates after 2017 and (2) also by the positive trend of Tajima’s D estimates at the *mtr* locus, (an indicator of the onset of balancing selection that typically happens during a population bottleneck (Biswas and Akey 2006)), and (3) along with the persistence of LD at the *mtr* locus maintained by recombination suggesting that the mosaic *mtrR* and promoter alleles are not yet fixed in the population (Figure 5). Unfortunately, based on the isolate collection ending in 2018 for this study, we cannot infer whether the compensatory mutations helped to recover clade 4 isolates from the population bottleneck unless we continue to track this clade beyond the year 2018.

In summary, after assembling and analyzing a global genome collection of ST-9363 genogroup isolates, we described the possible emergence and evolutionary forces that led to AZM^rs^ over time. Based on its genomic background and evolutionary trends, with the progressive acquisition of multiple genetic markers responsible for reduced susceptibility of many antibiotics such as mosaic *mtr* locus (AZM), mosaic *penA* (CFM/CRO), variations in GyrA and ParC (CIP), and PorB (TET), this genogroup has the potential to acquire additional antibiotic resistance markers. All these factors stress the importance of continued surveillance of this gonococcal genogroup both in the U.S. and worldwide, even when combination therapy with oral AZM is no longer recommended for gonococcal treatment but used for treating *Chlamydia* and other bacterial infectious diseases.

## Materials and Methods

### Isolate selection

In total, 1978 genome sequences from *N. gonorrhoeae* isolates belonging to the ST-9363 genogroup that were available as of February 2020 were included. This genogroup included MLST ST-9363 isolates and their closely related single/double locus MLST variants (total of 22 different MLSTs) that clustered within the previously described Clade A in the U.S. (Thomas et al. 2019; Gernert et al. 2020), which was also recently classified as “core-genome group 16”, identified using a locus threshold of 400 or fewer locus differences using a global collection of gonococcal genomes (Harrison et al. 2020). Genomes that matched these 22 MLSTs from the following sources were included: 1) isolates collected mostly through the Gonococcal Isolate Surveillance Project (GISP) and few isolates through Strengthening the US Response to Resistant Gonorrhea (SURRG) in the United States from 2000 to until 2018; 2) PubMLST (as of February, 2020); 3) PathogenWatch (as of February, 2020) and isolates from a number of published studies (Demczuk et al. 2016; Fifer et al. 2018; Harris et al. 2018; Yahara et al. 2018; Peng et al. 2019; Thomas et al. 2019; Williamson et al. 2019; Gernert et al. 2020; Golparian et al. 2020; Town et al. 2020). Only isolates with known years of isolation and country of origin were included. 60-70% of the isolates have recorded MIC or resistance classifications for CFM, CRO, CIP, AZM. The MLST profiles were rechecked and validated by StringMLST (Gupta et al. 2017) using the pubMLST database (Jolley et al. 2018). In total, 1080 genomes from North America, 659 genomes from Europe, 211 genomes from Oceania, 26 genomes from Asia and 2 genomes from South America spanning over 22 years (1996 -2018) were included in this study. To avoid biases in inferences, isolates from less represented geographical areas or continents were dropped appropriately in some analyses as described below. Genome data for an additional 35 ST-9363 genogroup isolates from Germany during 2018, which were released in March, 2021, were also added into the dataset and a separate timed phylogenetic analysis was only performed to understand its evolution compared to the global dataset.

### Whole genome sequencing and data analysis

All new *N. gonorrhoeae* isolates from GISP were sequenced on an Illumina sequencer at the CDC core sequencing facility (Atlanta, GA) or at either the Maryland Department of Health, Tennessee Department of Health, Texas Department of State Health Services or Washington State Department of Health, which are all part of the Antibiotic Resistance Laboratory Network regional laboratories as previously described (Grad et al. 2016; Thomas et al. 2019; Gernert et al. 2020). All raw Illumina sequencing reads were initially screened using Kraken v0.10.5 (Wood and Salzberg 2014), to identify reads assigned to bacterial strains other than *N. gonorrhoeae.* Samples with more than 10% of raw reads assigned to *Neisseria meningitidi*s or other bacterial species were considered contaminated. Fastq files were trimmed using CutAdapt v1.8.3 (Martin 2011) to remove adapter sequences and bases using a minimum Phred quality score cutoff of <Q30 and with a minimum length cutoff of 75 bps. Quality controlled Illumina reads were *de novo* assembled using SPAdes v3.9.0 (Bankevich et al. 2012) using the ‘careful’ option. All assemblies downloaded from PubMLST and other published sources were screened with Mash v2.2.2 for the presence of contigs from other bacterial taxa other than *N. gonorrhoeae*, and their quality were assessed by generating basic assembly statistics such as number of contigs, N50, L50 and GC content using bbmap (https://sourceforge.net/projects/bbmap/). A full-length whole genome alignment was generated using snippy v4.3.8 (https://github.com/tseemann/snippy) with WHO-P (GenBank accession number GCF_900087735.2; MLST ST-8127) set as the reference, which is included within the ST-9363 genogroup and core-genome group 16, and phylogenetically clustered as an outgroup. This full-length whole genome alignment was used as an input to Gubbins v2.3.1 (Croucher et al. 2015) for identifying and filtering regions of homologous recombination. The resulting alignment, containing only the polymorphisms present in the non-recombinant regions, was used as input for RAxML v8.2.9 (Stamatakis 2014) under the GTR+GAMMA model of nucleotide substitution with a majority-rule consensus (MRE) convergence criterion, to reconstruct an ascertainment bias corrected (Stamatakis method) maximum-likelihood (ML) phylogeny, which was used to determine the temporal signal for molecular clock analysis (see below).

*In silico* AMR genotypes (alleles or SNPs) were determined using an in-house customized script called AMR profiler (CDC) (Thomas et al. 2019) that uses raw reads and genome assemblies. This software also performs MLST and *N. gonorrhoeae* multi-antigen sequence typing (NG-MAST) typing schemes using StringMLST (Gupta et al. 2017) and NG-MASTER (Kwong et al. 2016), respectively.

### Timed Phylogenetic analysis

A time-scaled phylogenetic analysis using BEAST (Bayesian Evolutionary Analysis Sampling Trees) v1.8.4 (Suchard et al. 2018) was inferred using all 1978 genomes, collected over multiple years ranging from 1996 to 2018. TempEst v1.5.3 (Rambaut et al. 2016) estimated sufficient temporal signal (Root-to-tip R^2^ of 0.45) for molecular clock analysis based on the ML phylogenetic tree (inferred as described above) as input. The Gubbins alignment with SNP sites determined to be outside of the predicted recombinant segments was used as the input for BEAST 1.8.4 (Suchard et al. 2018). An ascertainment bias correction step was included by providing the count of the monomorphic sites for each nucleotide in the alignment within the BEAST input XML file generated using BEAUti. An HKY (Hasegawa, Kishino, and Yan) substitution model (one of the top 5 models out of 88 evolutionary models based on modelTest -NG (Darriba et al. 2020)), a coalescent model with constant population size, and an uncorrelated strict clock model with an exponential distribution as the prior for the clock rate were used for the BEAST analysis. Seven independent chains of 60 million steps were run, each was sampled every 1000 steps to ensure good mixing, and the initial 6 million steps of each run were discarded as burn-ins. The results were merged (total of 378 million steps) and compared using Tracer (v1.7.1) (Rambaut et al. 2018) to confirm convergence (Effective Sample Size (ESS) > 200), and the maximum clade credibility (MCC) tree was generated using Tree Annotator; which are both provided within the BEAST package (Suchard et al. 2018). The final timed phylogenetic tree was visualized and annotated using the R package ggtree (Yu et al. 2017). Analysis of population structure within the MCC timed tree was then conducted using the nonparametric TreeSructure (Volz et al. 2020) approach, which partitions the tips and internal nodes of a tree into discrete sets characterized by comparable coalescent patterns (with 100,000 tree simulations and a significance threshold of 0.05), to identify major clades. The effective population size (EPS) trajectory over time, which was scaled to the number of generations per year, for each of the clades was modelled using the skygrid model implemented using the R package phylodyn v0.9 (Karcher et al. 2017) with the default parameters. A separate timed phylogenetic analysis (n=2013) using an additional 35 isolates very recently released (March 2021) from Germany (year of isolation 2018) was also performed using BEAST as described above.

### Phylogeographic and global transmission trend analyses

Phylogeographical inferences over time were made for the MCC time-calibrated phylogeny using ML ancestral state reconstruction in order to understand the likely geographical origin of the ST-9363 genogroup by treating isolate location as a discrete trait as implemented in the R package ape (Paradis and Schliep 2019). Only isolates from North America, Europe and Oceania were used in this analysis in order to avoid biases due to the very low number of isolates from Asia and South America. The ancestral states were reconstructed using the three standard models - equal rates model (ER), symmetric model (SYM) and all rates different (ARD) model. Out of the three models tested, the ARD model estimated the highest log likelihood, and the log likelihood tests (known as the G statistic) performed to compare ARD model against ER (p-value=2.629616e-08) and SYM (p-value = 0.02372808) models yielded a significant p-value; confirming that the ARD model is the best model in this analysis. The estimated posterior probabilities of each of the discrete traits - Europe, North American and Oceania - in each of the internal nodes of the timed phylogeny are shown as pie charts using the R package ggtree (Yu et al. 2017).

To preclude any biased evolutionary inferences made from the timed phylogenetic and phylogeographical analysis due to unbalanced sampling across different geographical regions, we randomly generated a down-sampled dataset and re-ran both the analyses. The full dataset contained 211 isolates from Oceania, and we decided to keep all those Oceania genomes and then we randomly selected 211 isolates each from North America and Europe. All the Asian and South American isolates were excluded. Snippy, Gubbins and BEAST analysis were performed to estimate the timed phylogeny on which phylogeographic analysis was repeated as described above, and the ARD model (log likelihood ratio tests against ER (p-value = 1.3507e-07) and SYM (p-value = 0.4857)) was also determined as the best model for the down sampled dataset.

TransPhylo (Didelot et al. 2017) was used to estimate the global transmission trends and reproductive number (R0). Only isolates that belonged to the European and North American lineages were included in this analysis. The implementation and parameters chosen were similar as described previously (Osnes et al. 2020). Briefly, for the generation time distribution (time from being infected to infecting others), two gamma distributions where considered, one which was estimated in an individual-based modelling study from a community of men who have sex with men (MSM) (Whittles et al. 2019), which is called “prior 1” with shape 0.57 and scale 0.30, and a second distribution similar to prior 1 but penalizing transmissions in the incubation phase - “prior 2” with shape 1.20 and scale 0.14. The sampling distribution (time between infection and sampling) was set equal to the generation time distribution. The sampling densities were fixed to the fractions of 0.2, 0.3, 0.4 and 0.5 of the number of cases and Transphylo was run for each of the two parameter priors and sampling fractions. For each configuration, one million MCMC iterations were performed, and the number of iterations were tinned down to 50000 with the first half of the iterations discarded as burn-ins. Sensitivity testing of the outputs from all the eight possible combinations of sampling and generation time priors indicated that the modelling output was sensitive to the choice of generation time and sampling density priors for the ST-9363 genogroup used in this study (Figures S9(a) and S9(b)). For transmission modelling, we used the results generated with a sampling density of 0.4 and prior 1 for the generation time.

### Clade by clade characterization of the recombination events

Recombination events were first detected using Gubbins on the whole genome alignment of 1978 isolates as described above. Estimates ρ/θ and r/m for each of the clades were derived from the Gubbins output. For each shared recombination event that occurred once and spread via clonal descent, the recombination statistics were only counted once. To test if there were any differences in recombination rates among the lineages and the clades, we used the Kruskal-Wallis nonparametric analysis of variance (Kruskal and Wallis 1952) on all the estimated ρ/θ values per lineages and clades, followed by Dunn’s test for post hoc statistical testing (Dunn 1964) for differences between lineages and clades along with Hochberg’s correction (Hochberg 1988) for multiple testing using the R package called ‘dunn.test’ (https://cran.r-project.org/web/packages/dunn.test/index.html). Recombination hotspot regions for selected clades were found by creating separate Manhattan-plots of the number of recombination events happened per 1000 bp discrete window across the genome at the internal nodes (ancestral recombination events) and those events in the terminal branches (current recombination events) that represent independent recent acquisitions, and manually looking for peaks. For some selected clades genome wide estimates of Tajima’s D statistics and linkage disequilibrium (LD) derived due to recombination within the *mtr* locus were assessed from the clade-specific whole genome alignment over 100-bp sliding windows using the R package called PopGenome (Pfeifer et al. 2014).

## Supporting information

All Supplementary Figures

All Supplementary Tables

## Acknowledgements

We thank Alesia Harvey for assistance for verifying isolate epidemiologic and MIC data. We also thank GISP and all sentinel sites that contributed isolates through GISP and SURRG during 2011 – 2018. This work was supported in part by the Centers for Disease Control and Prevention (CDC)’s Combating Antibiotic Resistant Bacteria initiative and partially by Advanced Molecular Detection.

## Disclaimer and conflicts of Interest

The authors declare no competing interests. The findings and conclusions in this report are those of the authors and do not necessarily represent the official position of the CDC.

## Supplementary Data

**Figure S1.** Whole genome sequence dated phylogeny for each of the 32 clades detected by the TreeStructure algorithm based on maximum clade credibility reconstructed using BEAST. Demographic history of each of the 32 clades are also shown where the time is measured in years and 0 indicated the most recent year of isolate sampling per clade (x-axis) and the effective population size is scaled to the number of generations per year for each clade (y-axis). Most recent years of isolates sampled, geographical location and the major MLSTs in each clade are also mentioned.

**Figure S2.** Whole genome sequence dated phylogeny with the maximum clade credibility reconstructed using BEAST along with the known resistance conferring genetic variants for AZI of all the 1978 ST-9363 genogroup + 35 isolates recently released in March 2021 from Germany (n=2013) gonococcal isolates included in this study. The clade with the additional 2018 German isolates clustered within the European lineage is highlighted. White/blank regions in the figure indicate that the data is not available for the corresponding isolate on the phylogenetic tree. *Azithromycin MIC non-susceptible >= 1 are shown for the Oceania isolates as the exact MICs were not publicly available and in Williamson et al. (2019) all Oceania isolates with MICs >= 1μg/ml were considered as resistant to AZM, which does not conform to CLSI breakpoint. HL-AZI-R stands for “High-level Azithromycin resistance”.

**Figure S3**. Phylogeographic ancestral state reconstruction based on the maximum clade credibility reconstructed using BEAST with geographical locations (Continents) as discrete traits on the down sampled dataset. Each node indicates the estimated posterior probabilities of each of the discrete traits - Europe, North American and Oceania as pie charts

**Figure S4.** The estimated medoid transmission tree inferred using TransPhylo. White circles are inferred unsampled individuals. Tip points are colored after the geographical location. Only the European and North American lineage isolates (as denoted in figures 1 and 2) were included in this analysis

**Figure S5**. Bar diagram showing the distribution of the ρ/θ (recombination rates) estimates inferred using Gubbins for each of the 32 clades. The bars are ordered and colored based on the lineages of each of the clades – red color for the basal lineage clades, green color for the European lineage clades and blue color for the North American lineage clades.

**Figure S6**. Manhattan plots of the number of recombinations per 1000 base-pair window for the North American lineage, Clade 31 with isolates that contained the mosaic *mtr* locus alleles. The number of recombination events, both ancestral and current events per 1000 bps window are shown in the y-axis and the genomic location along the reference WHOP genome is shown in the x-axis. Regions that are hotspots for recombination are shown as red circles and the table shows the genes present in those recombination hotspots.

**Figure S7**. Manhattan plots of the number of recombinations per 1000 base-pair windows for the North American lineage, Clade 4. The number of recombination events, both ancestral and current events per 1000 bps window are shown in the y-axis and the genomic location along the reference WHOP genome is shown in the x-axis. Regions that are hotspots for recombination are shown as red circles and the table shows the genes present in those recombination hotspots.

**Figure S7**. Manhattan plots of the number of recombinations per 1000 base-pair windows for the European lineage, Clade 17. The number of recombination events, both ancestral and current events per 1000 bps window are shown in the y-axis and the genomic location along the reference WHOP genome is shown in the x-axis. Regions that are hotspots for recombination are shown as red circles and the table shows the genes present in those recombination hotspots.

**Figure S8**. Bar diagrams showing the estimated Tajima’s D statistics estimated for the whole genome, the entire *mtr* locus genes, *mtrD* gene and the *mtrR* and promoter region for the clade 4 isolates, and the estimated linkage disequilibrium (LD) measure by r^2^ for the entire *mtr* locus, *mtrD* gene and the *mtrR* and promoter region for few clades form North American and European lineages. Both Tajima’s D and r^2^ were estimated over a 100 bp sliding window on the entire whole genome alignment containing only the clade 4 isolates using the R package called PopGenome. **a)** Clade 31 (no mosaic *mtrR* and *mtrR* promoter mutation/substitution) **b)** Clade 4 **c)** Clade 31 (with mosaic *mtrR* and *mtrR* promoter mutation/substitution) **d)** Clade 29 **e)** Clade 17 **f)** Clade 23 **g)** Clade 24 and **h)** Clade 1

**Figure S9**. **a)** Convergence diagnostics of the Transphylo MCMC chains for prior 1. Each of the columns shows the traces of the MCMC iterations for each sampling density at 0.2, 0.3. 0.4 and 0.5. The plots in the first row show the chains for within-host effective population size neg, the second row the reproductive number R, and the third row shows the chain for the transmission tree. In all cases, the chains seemed to have achieved sufficient convergence with effective sampling sizes (ESS) > 200. **b)** Convergence diagnostics of the Transphylo MCMC chains for prior 2. Each of the columns shows the traces of the MCMC iterations for each sampling density at 0.2, 0.3. 0.4 and 0.5. The plots in the first row show the chains for within-host effective population size neg, the second row the reproductive number R, and the third row shows the chain for the transmission tree. The chains for within-host effective population size seemed to have not achieved sufficient convergence with effective sampling sizes (ESS) < 20.

**Table S1.** Isolate information with all the associated metadata and antimicrobial resistance (AMR) genetic variants associated with all the isolates used in this study.

**Table S2**. **a)** Comparison of the BEAST parameter estimates for the full dataset and the down sampled dataset **b)** Comparison of the tMRCA estimated for the major historic events using the full dataset and the down sampled dataset.

**Table S3.** Average recombination statistics per clade

**Table S4.** Results of the per clade pairwise Dunn’s test for comparing recombination rates across clades

