## Supplementary material for "Global emergence and dissemination of *Neisseria gonorrhoeae* ST-9363 isolates with reduced susceptibility to azithromycin": All Supplementary Figures

### Figure S1

#### TreeStructure Clades

##### Clade 1

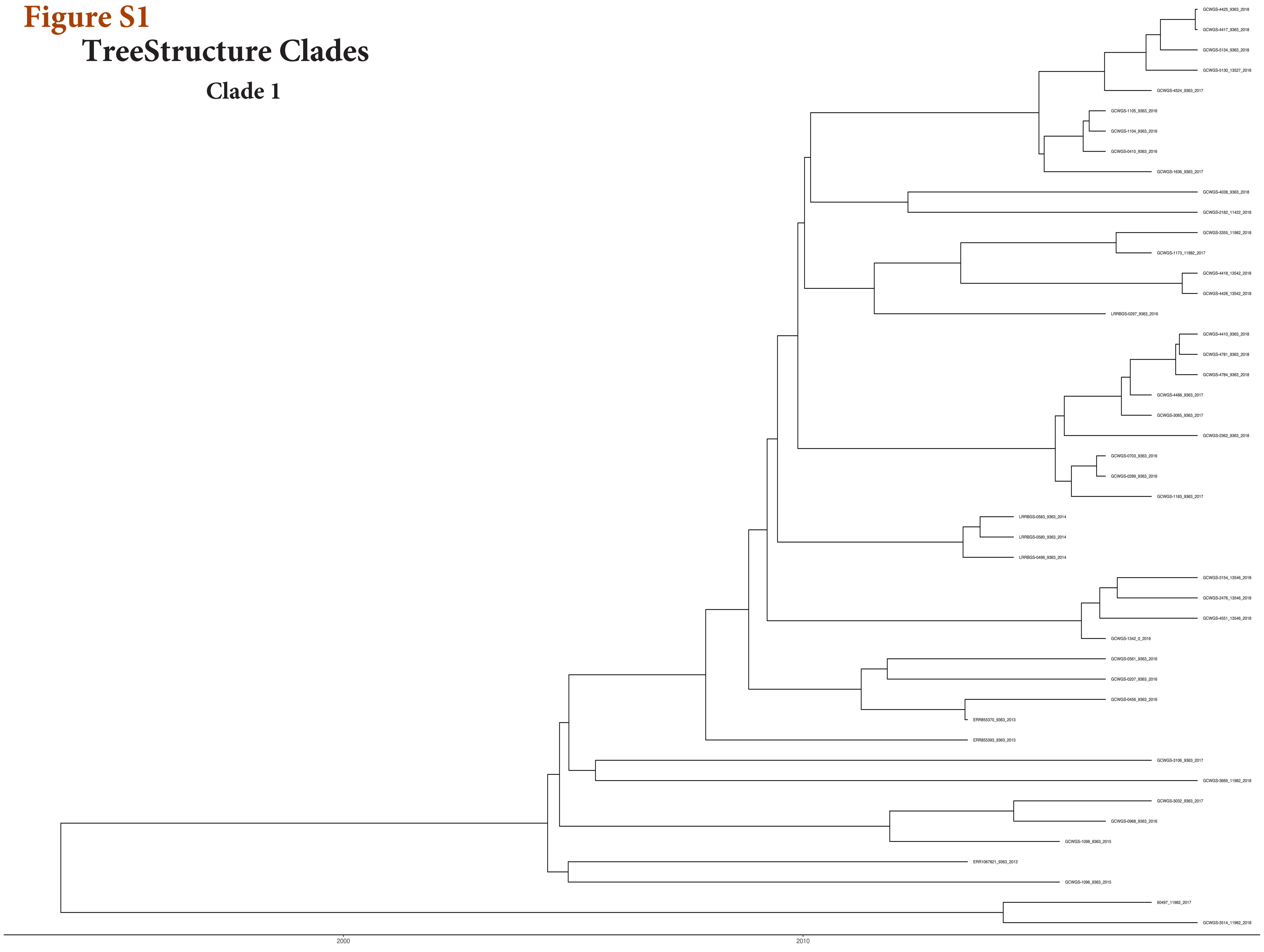


**Major MLST - 9363**
**Majority of Isolates are from North America**
**Most Recent year of isolate - 2018**

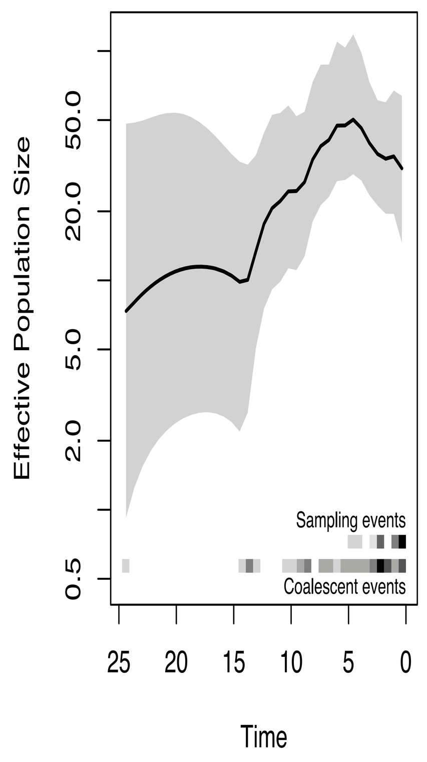


##### Clade 2

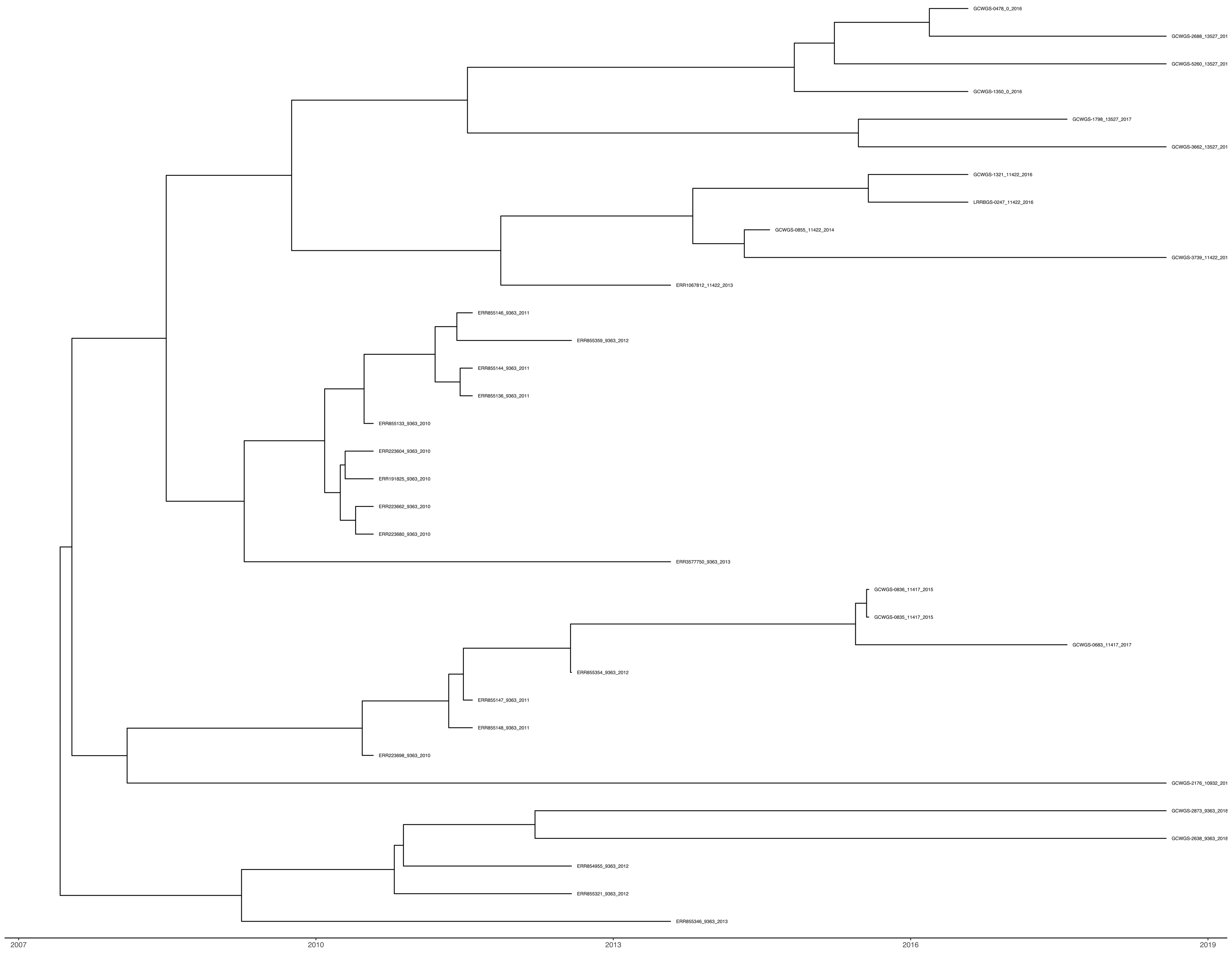


**Major MLST - 9363**
**Majority of Isolates are from North America**
**Most Recent year of isolate - 2018**

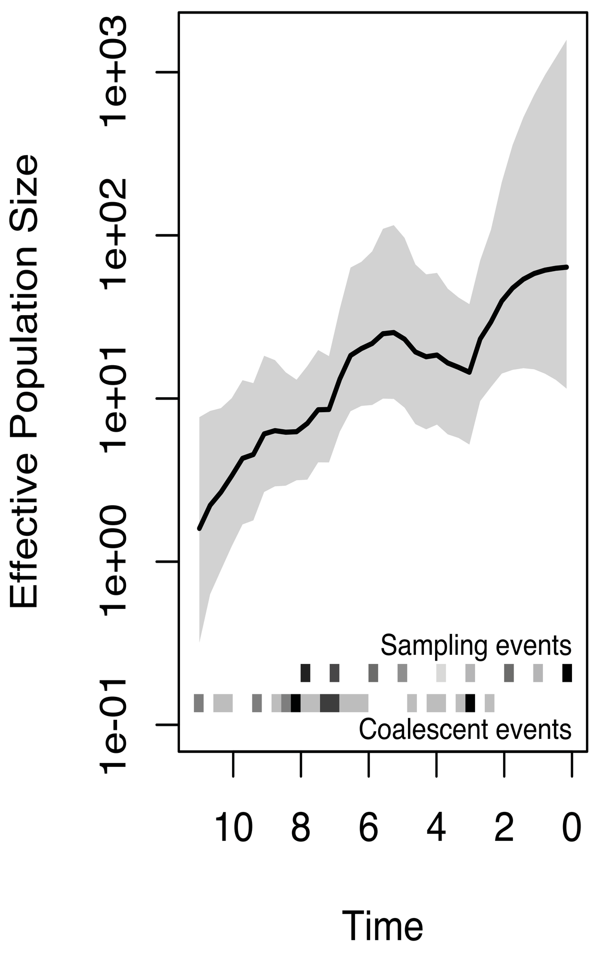


##### Clade 3

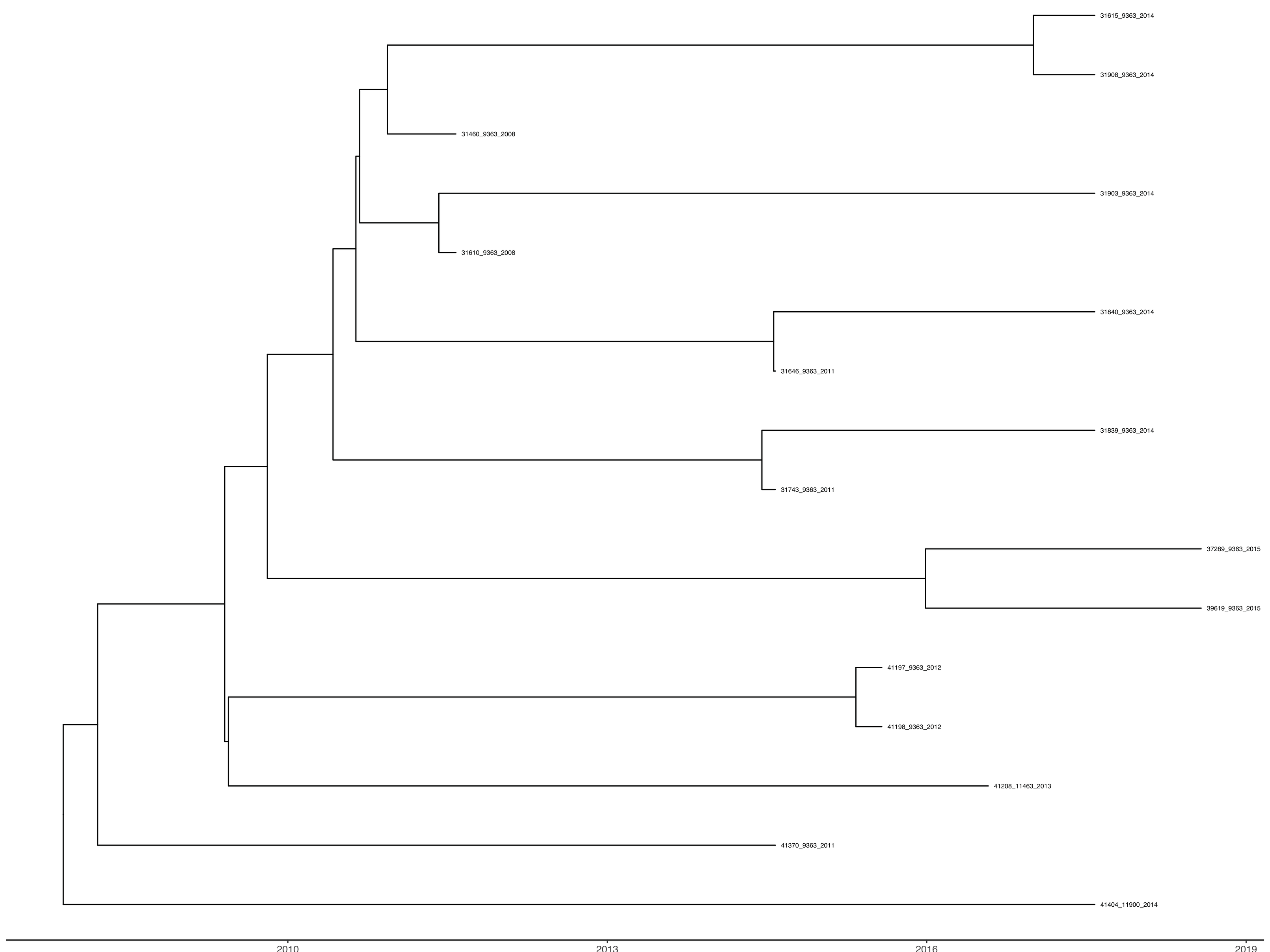


**Major MLST - 9363**
**Majority of Isolates are from Europe**
**Most Recent year of isolate - 2015**

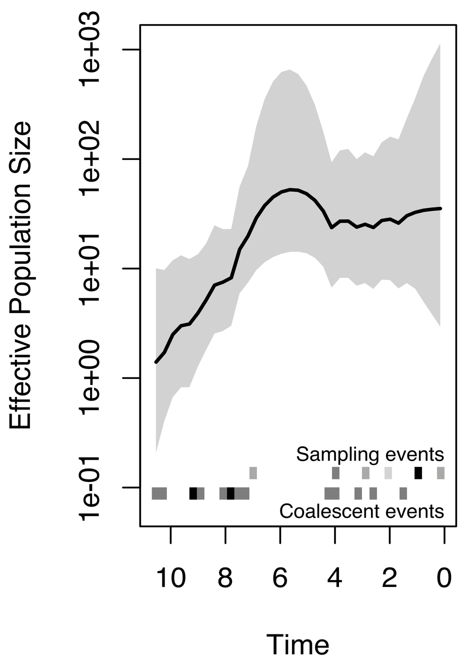

**Clade 4**

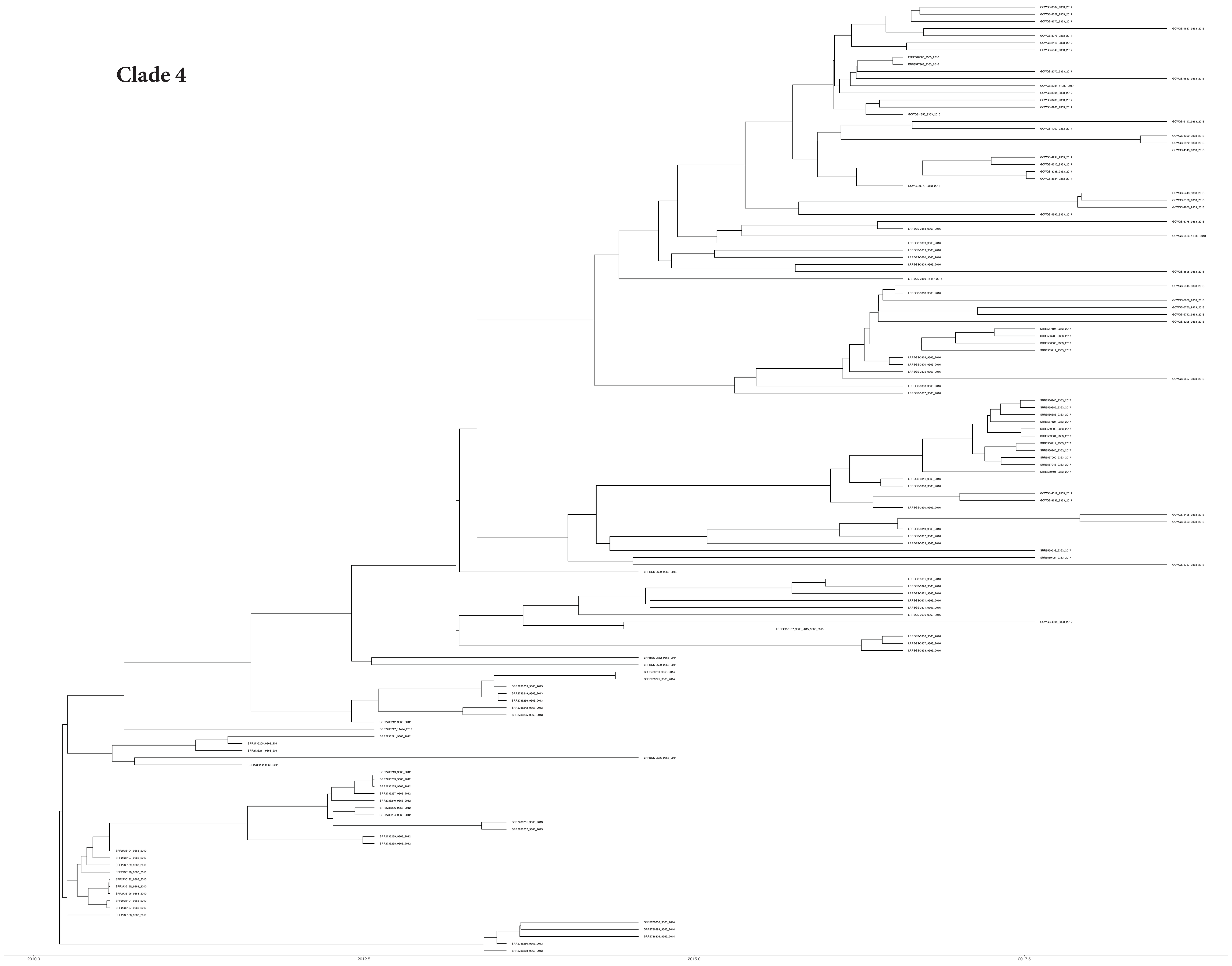

**Major MLST - 9363**
**Majority of Isolates are from North America**
**Most Recent year of isolate - 2018**

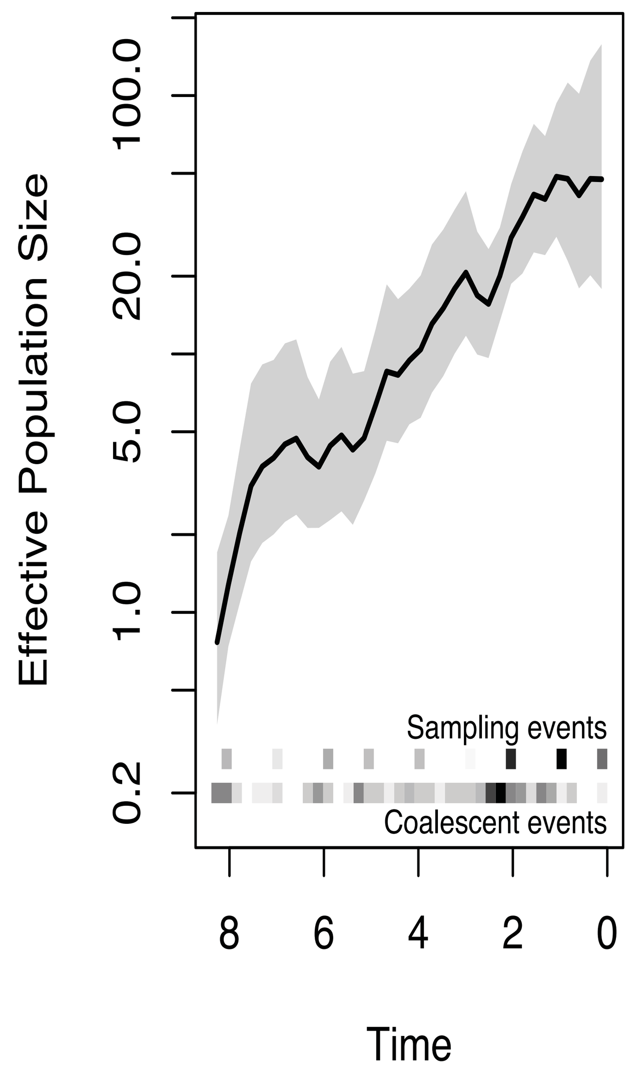


**Clade 5**

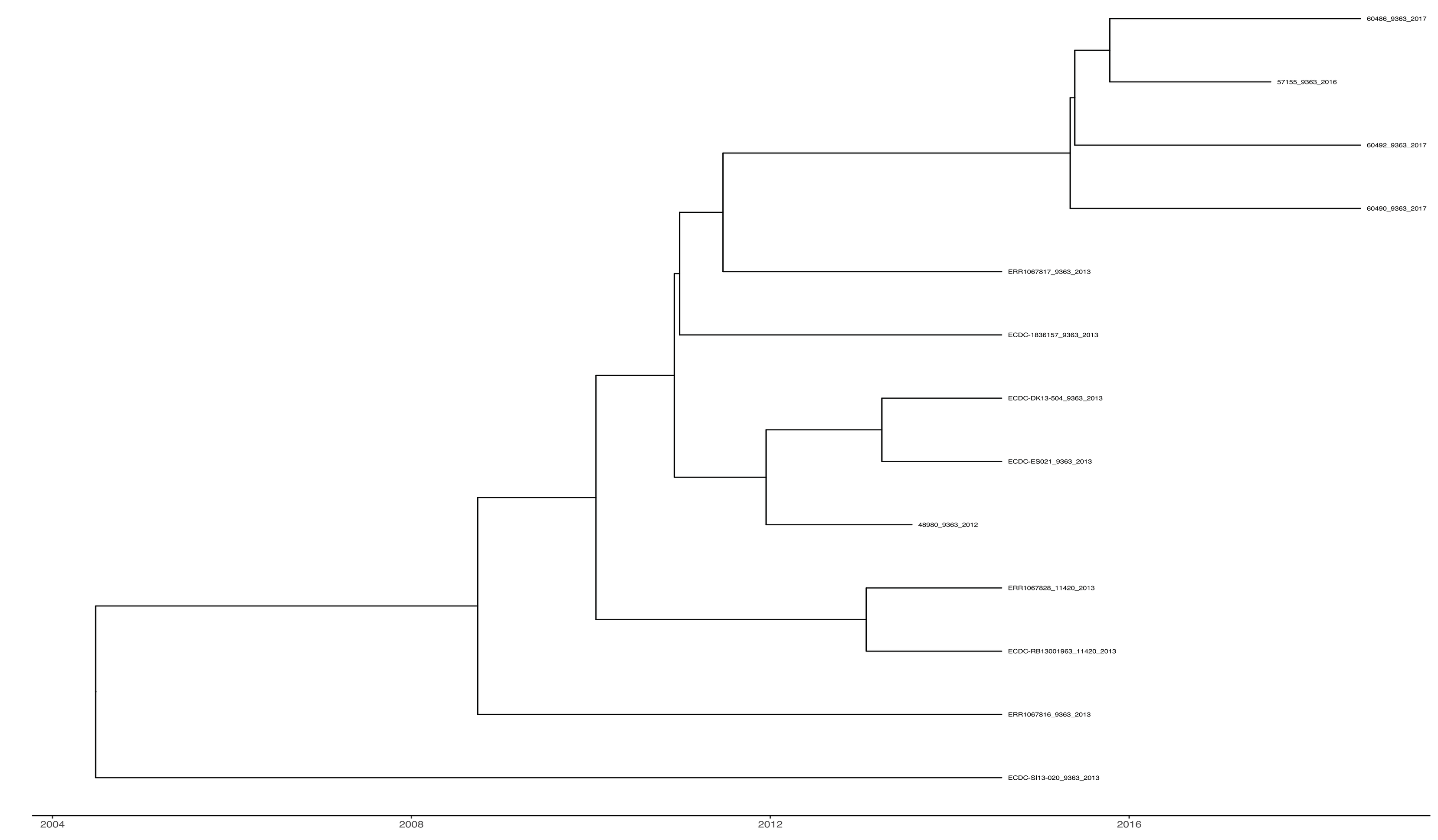


**Major MLST - 9363**
**Majority of Isolates are from Europe**
**Most Recent year of isolate - 2017**

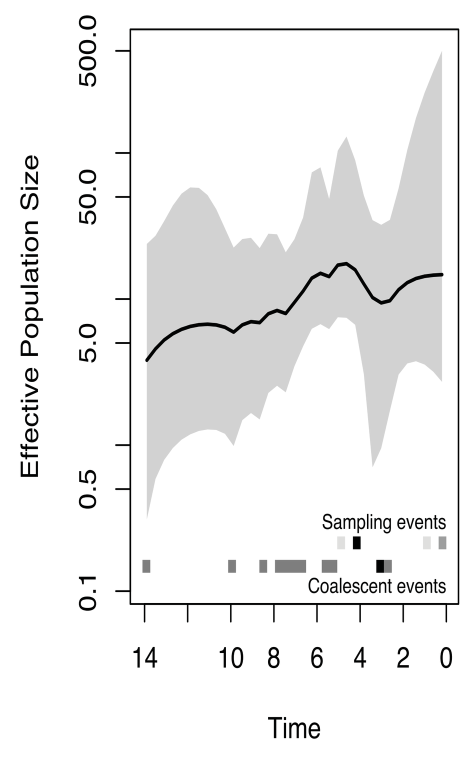


**Clade 6**

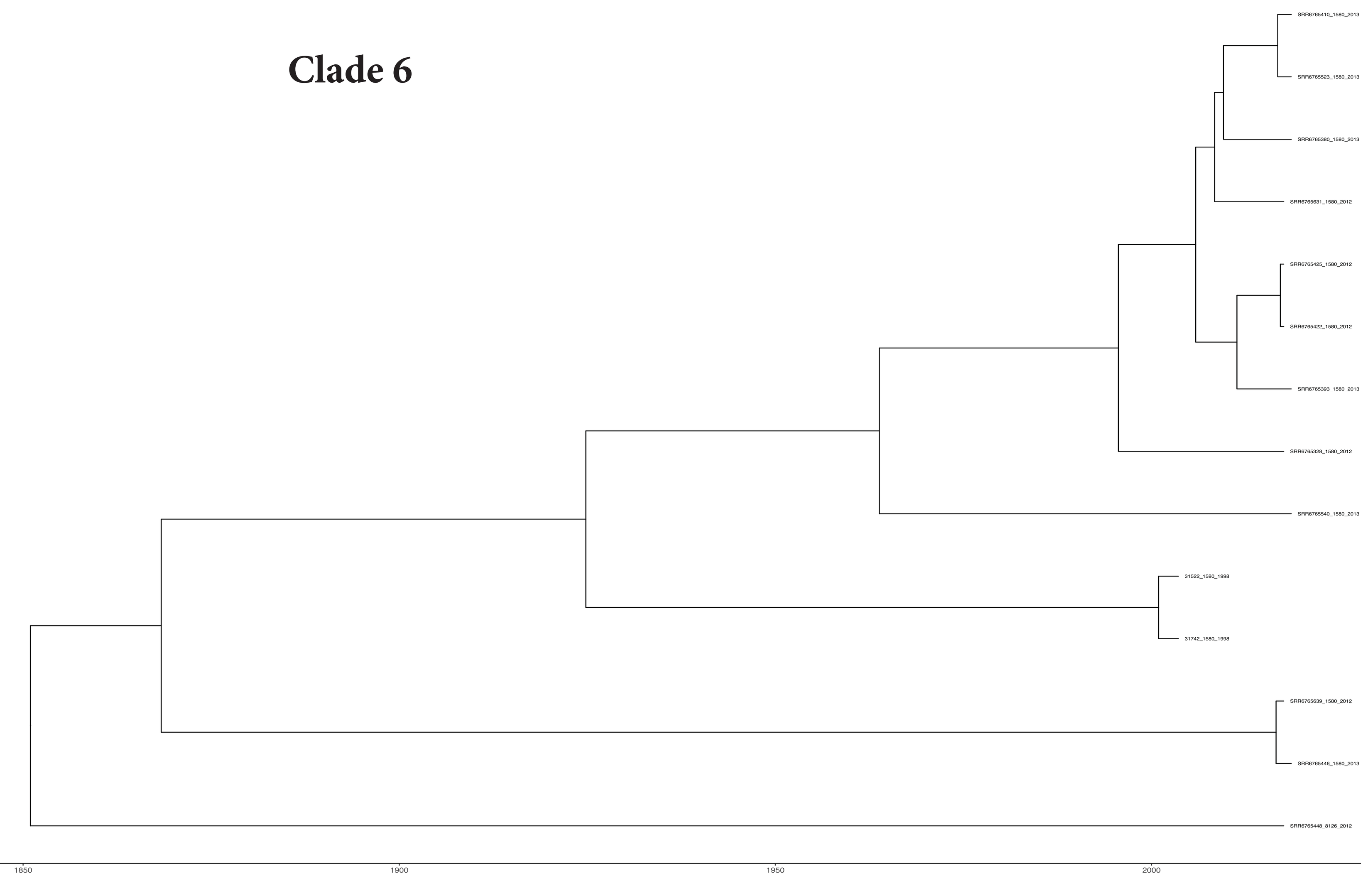

**Major MLST - 1580**
**Majority of Isolates are from Asia**
**Most Recent year of isolate - 2013**

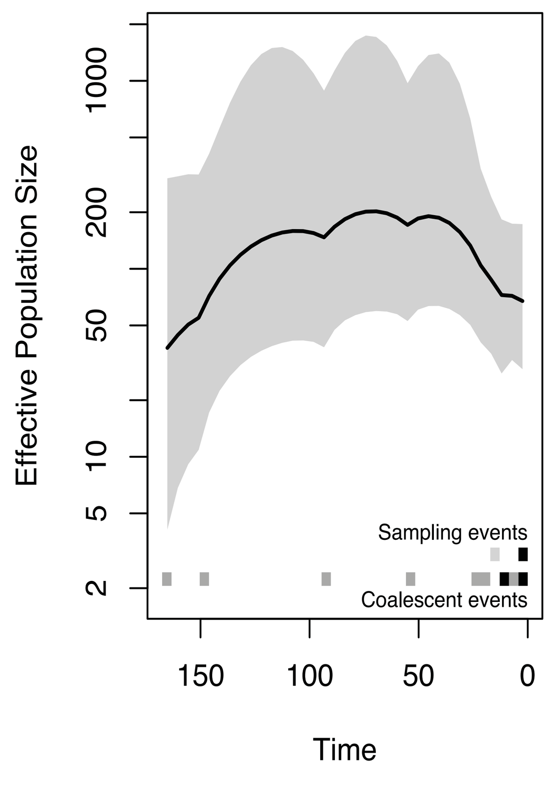

#### Clade 7

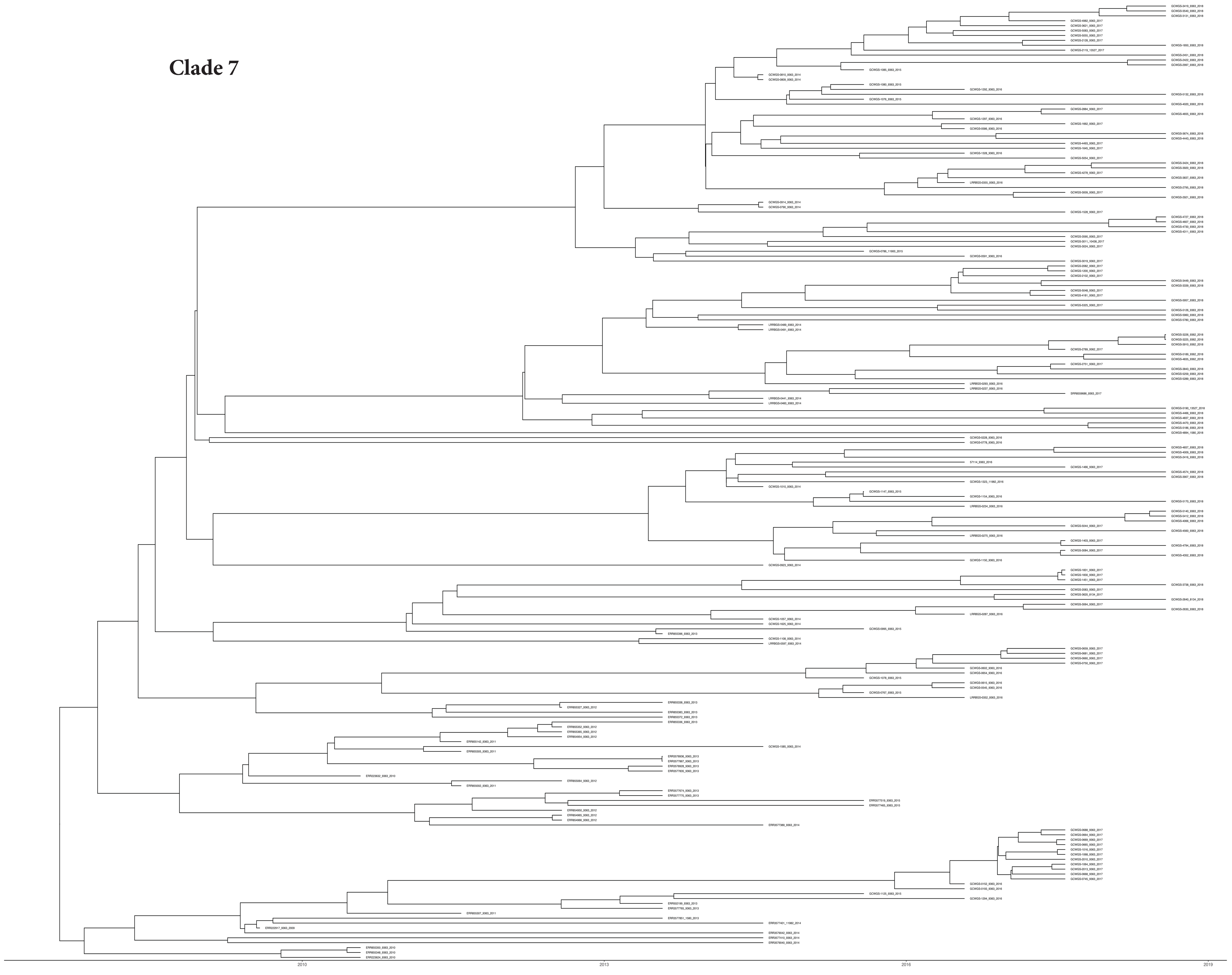

**Major MLST - 9363**
**Majority of Isolates are from North America**
**Most Recent year of isolate - 2018**

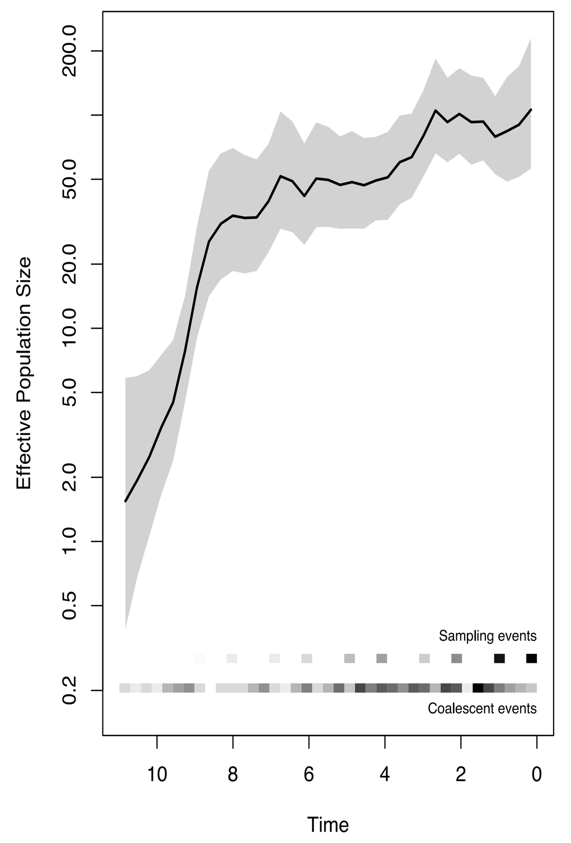


#### Clade 8

**Major MLST - 9363**
**Majority of Isolates are from North America and Oceania**
**Most Recent year of isolate sampling - 2018**

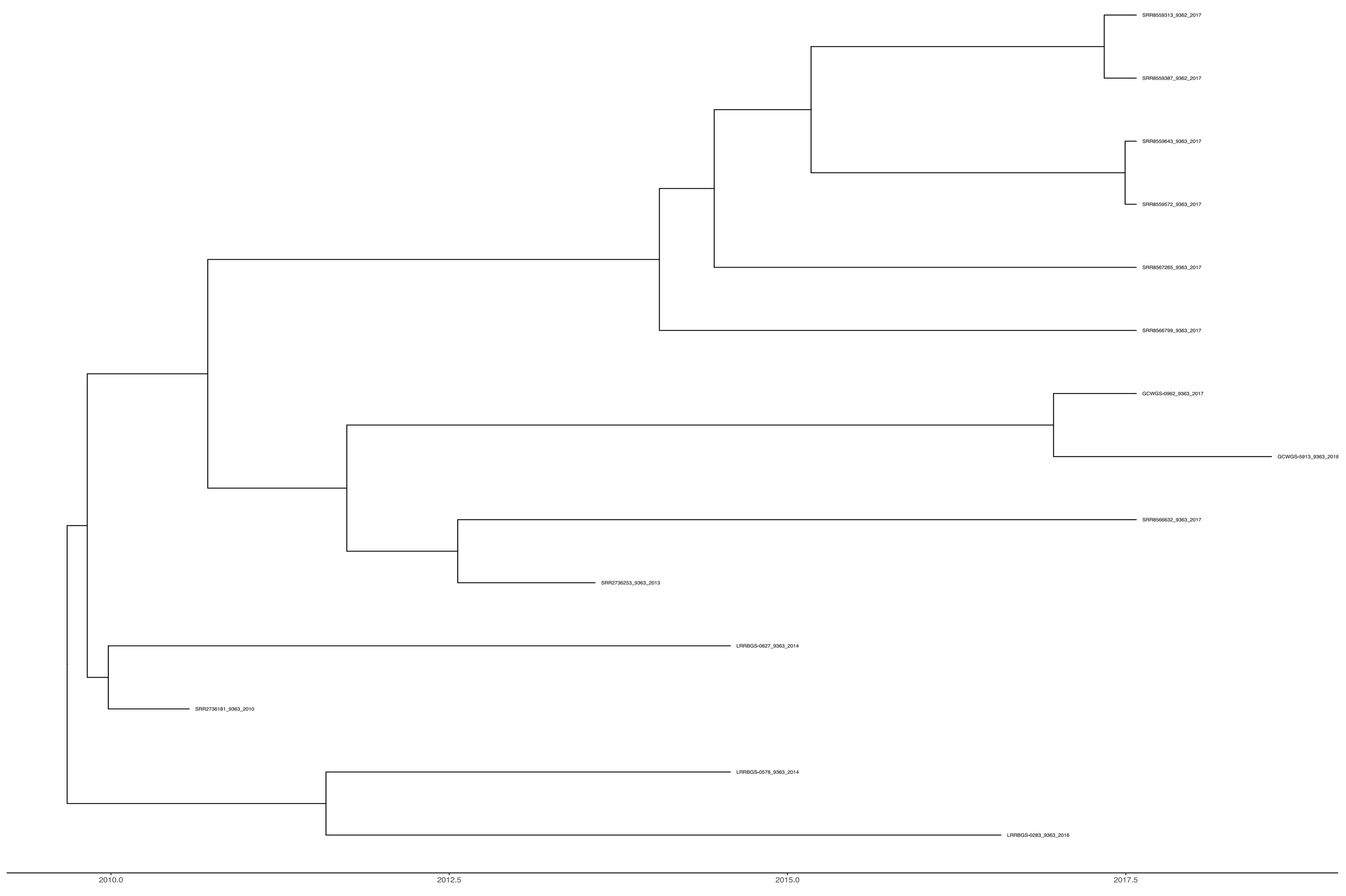

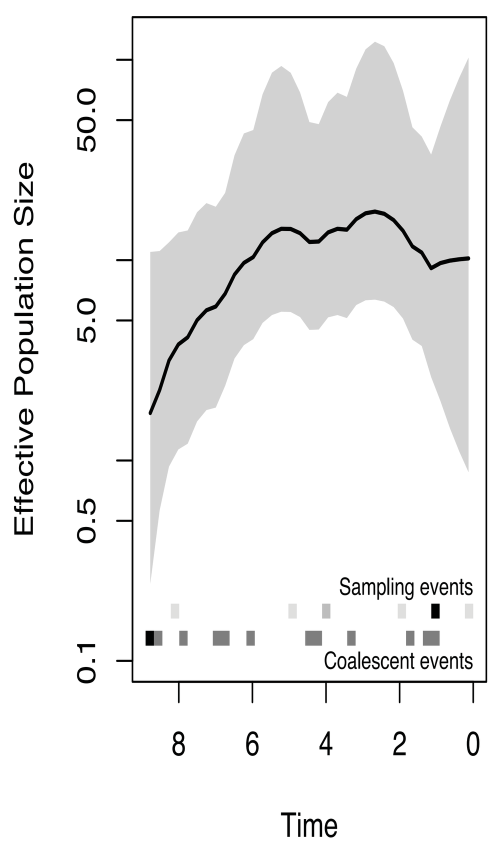


#### Clade 9

**Major MLST - 1580 and 8126**
**Majority of Isolates are from North America**
**Most Recent year of isolate sampling- 2017**

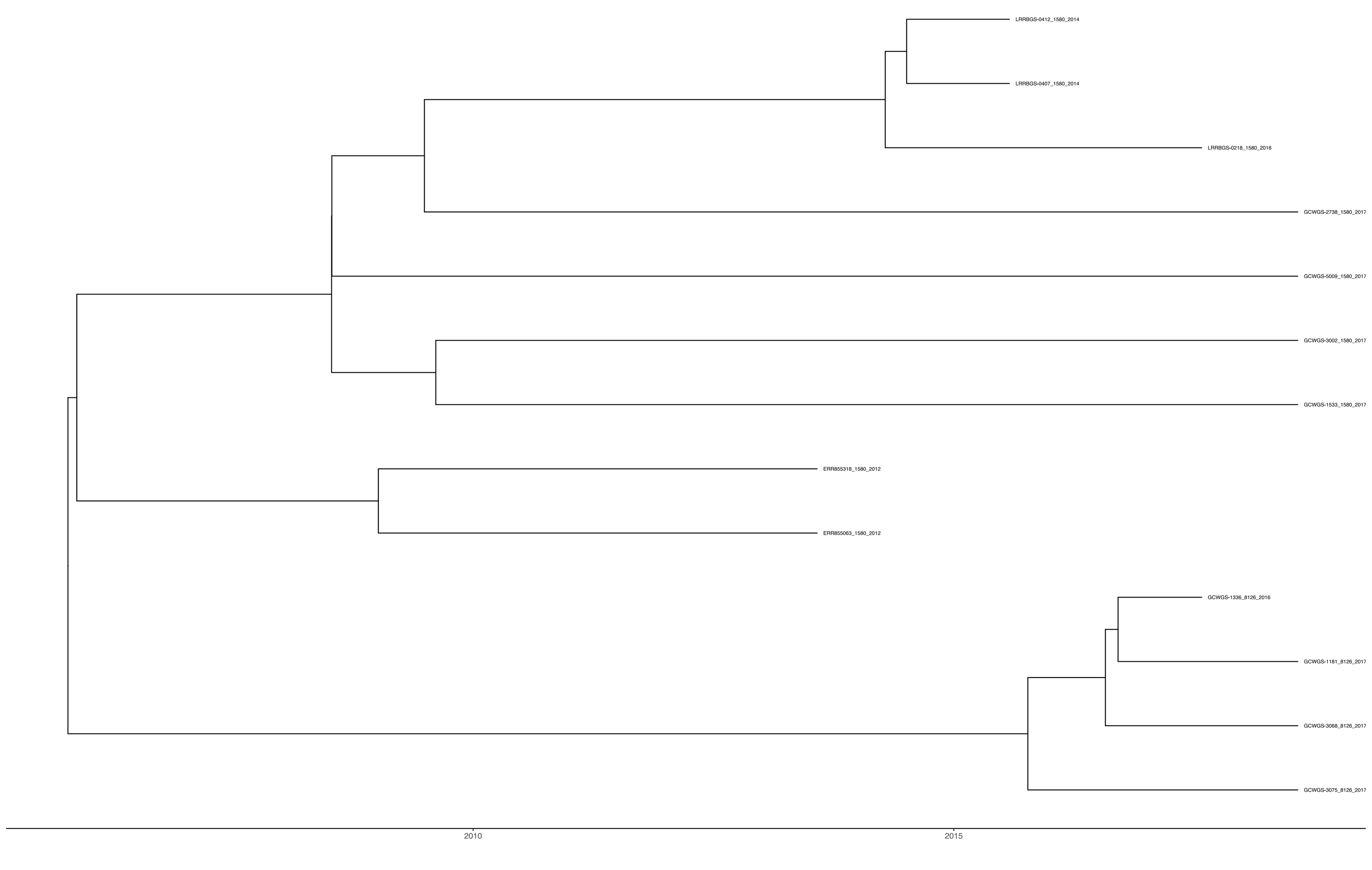

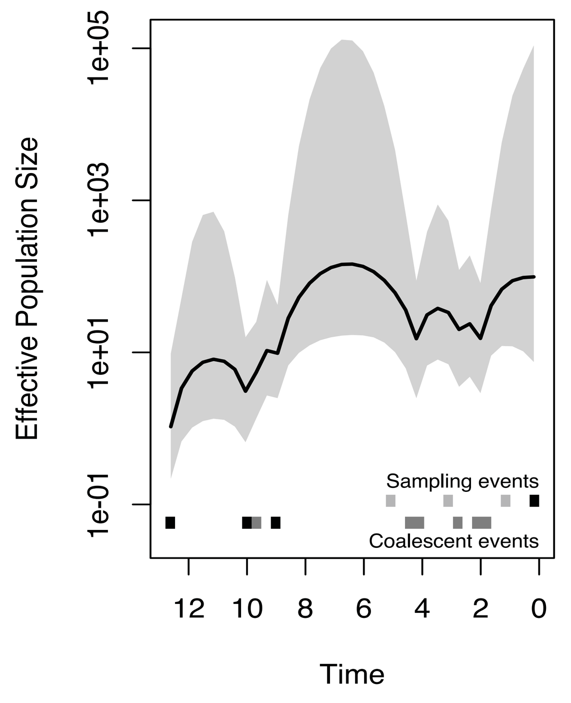

**Clade 10**

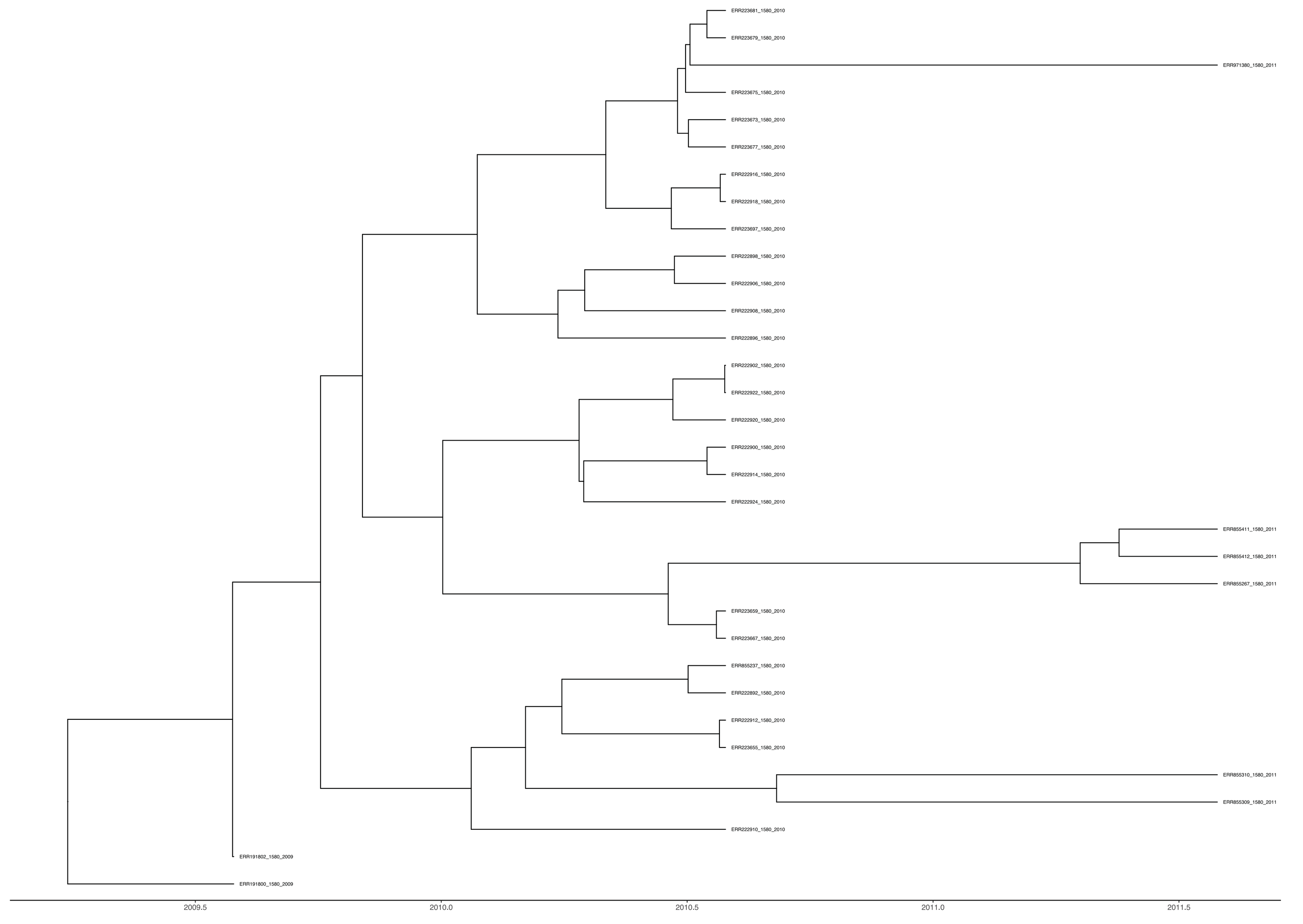

**Major MLST - 1580**
**Majority of the isolates are from North America**
**Most recent year of isolate - 2011**

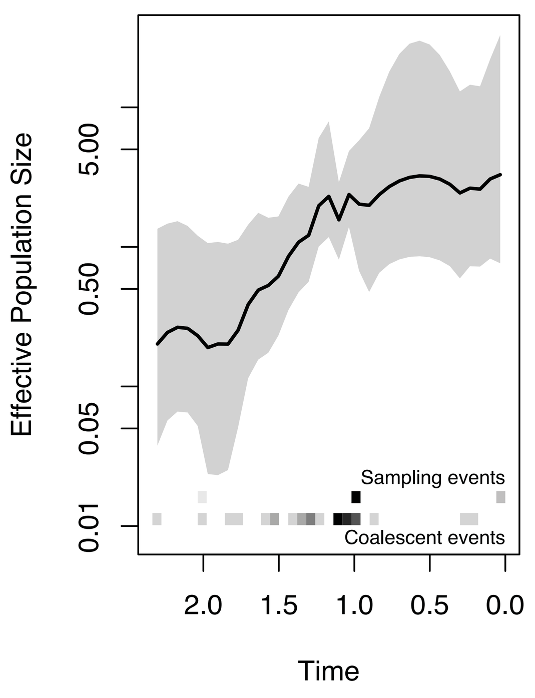


**Clade 11**

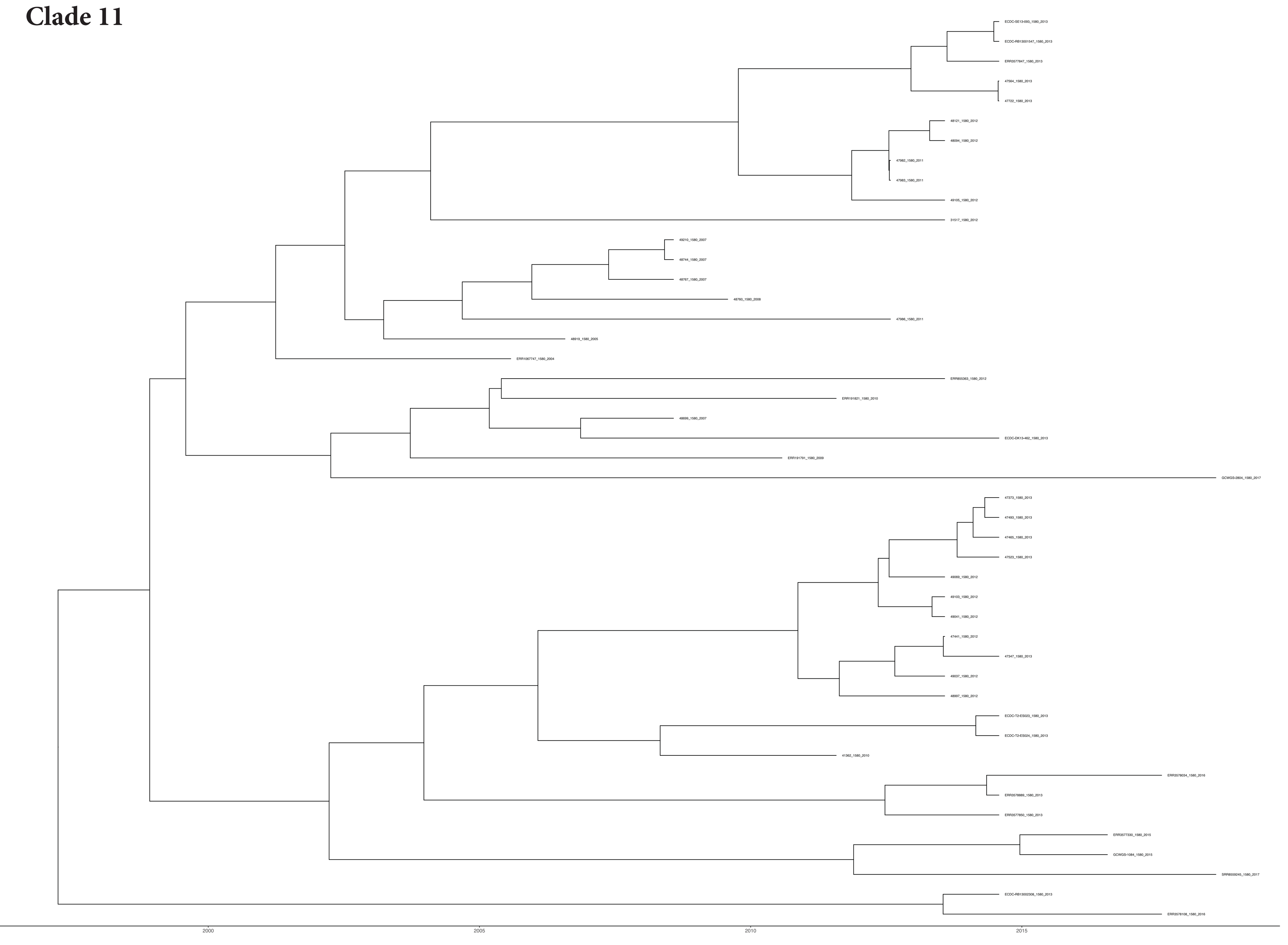

**Major MLST - 1580**
**Majority of the isolates are from Europe**
**Most recent year of isolate - 2017**

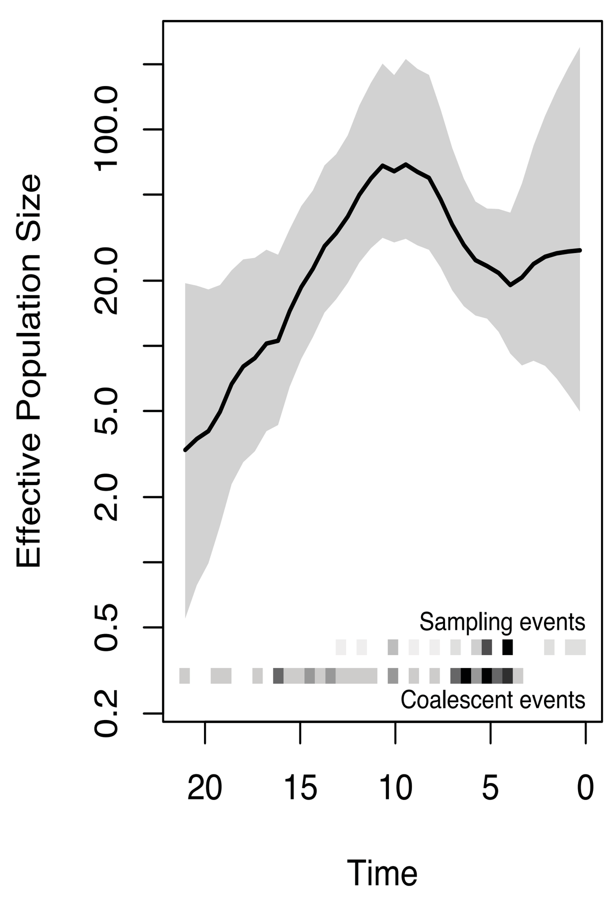


**Clade 12**

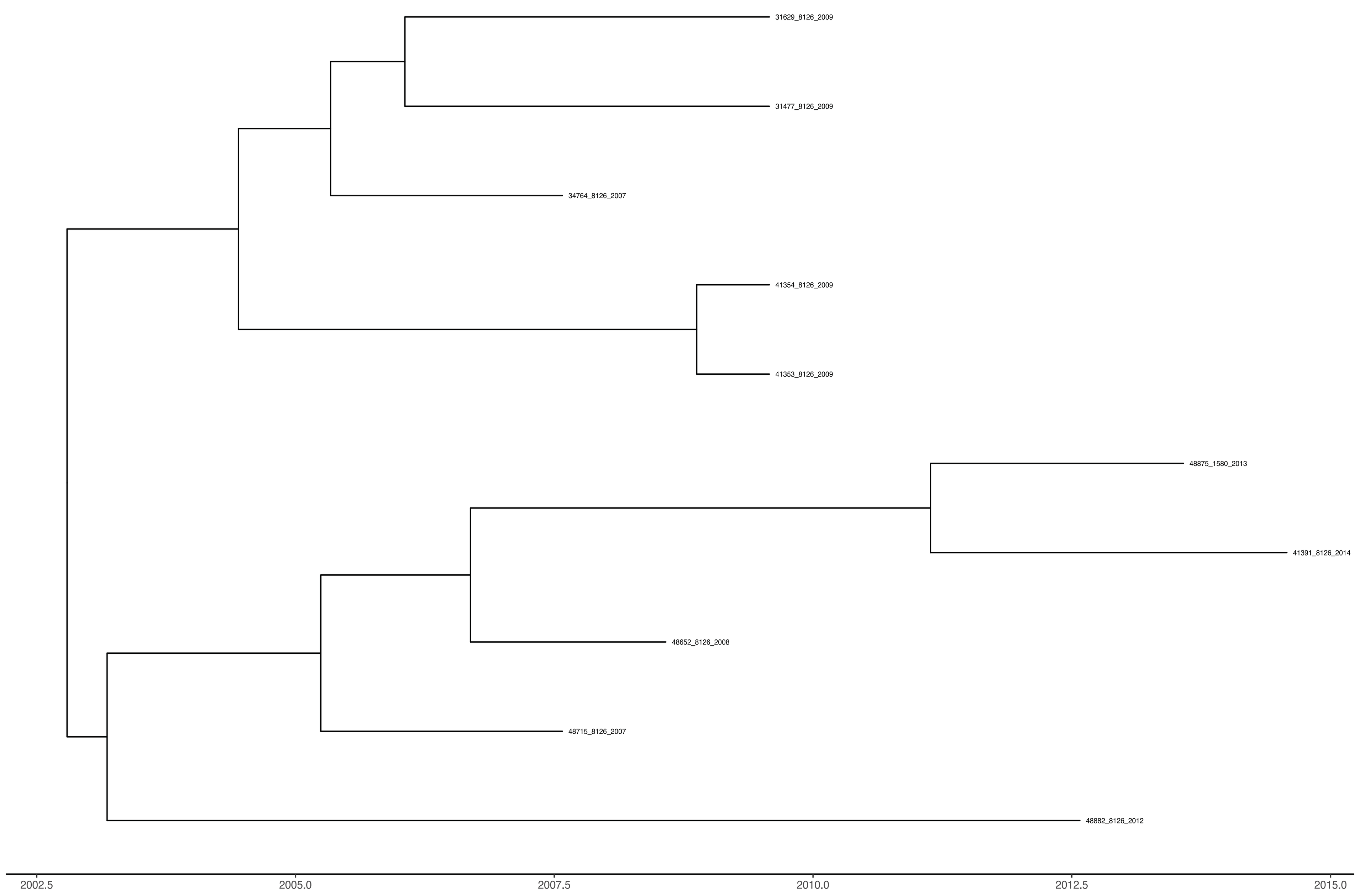


**Major MLST - 8126**
**Majority of the isolates are from Europe**
**Most recent year of isolate - 2012**

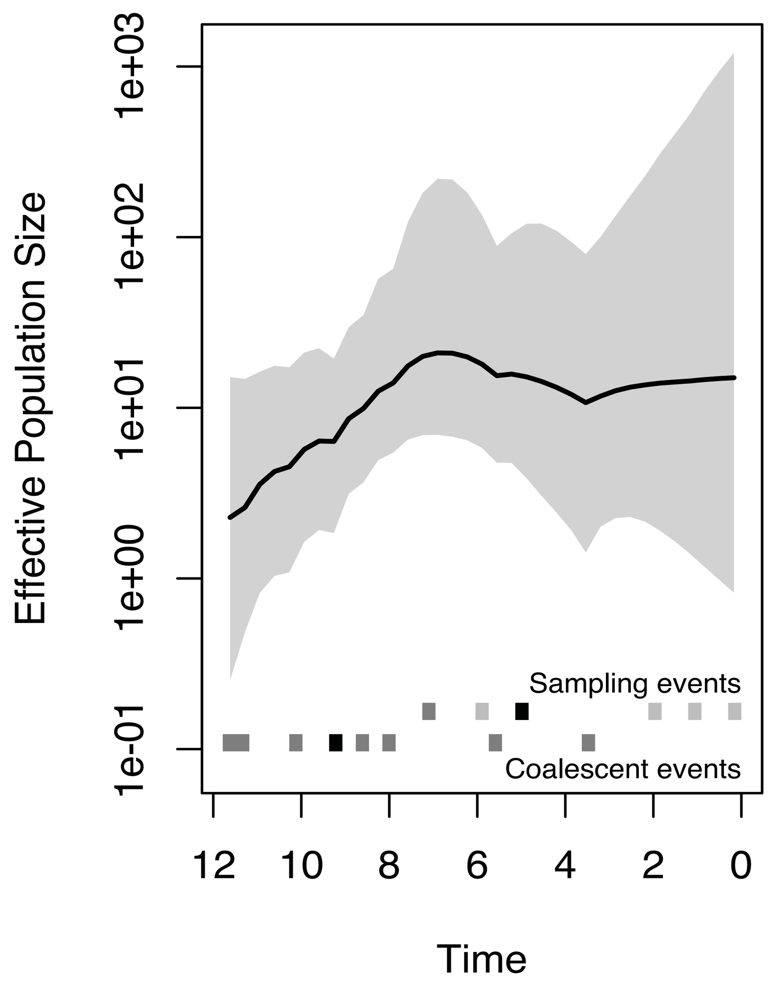

#### Clade 13

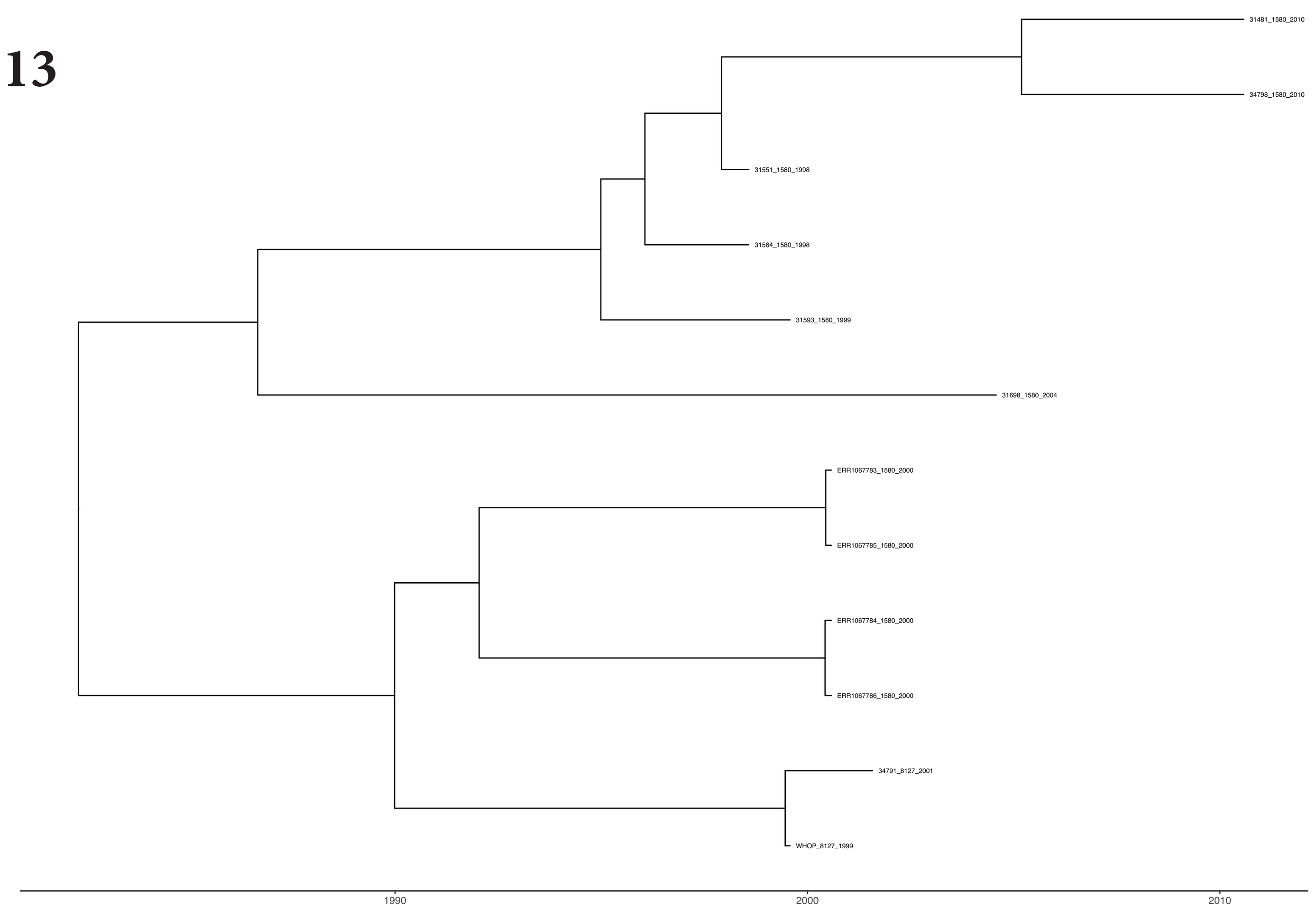


**Major MLST - 1580**
**Majority of Isolates are from Europe**
**Most recent year of isolate sampling- 2004**

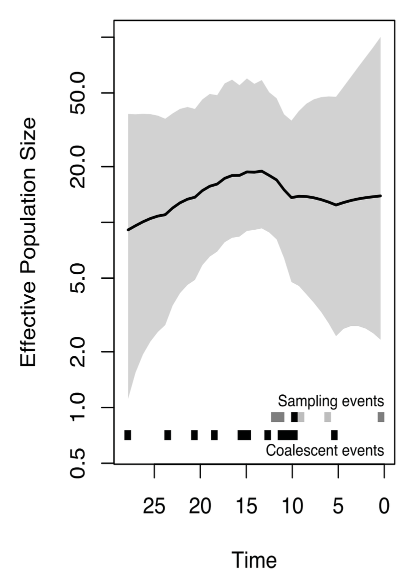


#### Clade 14

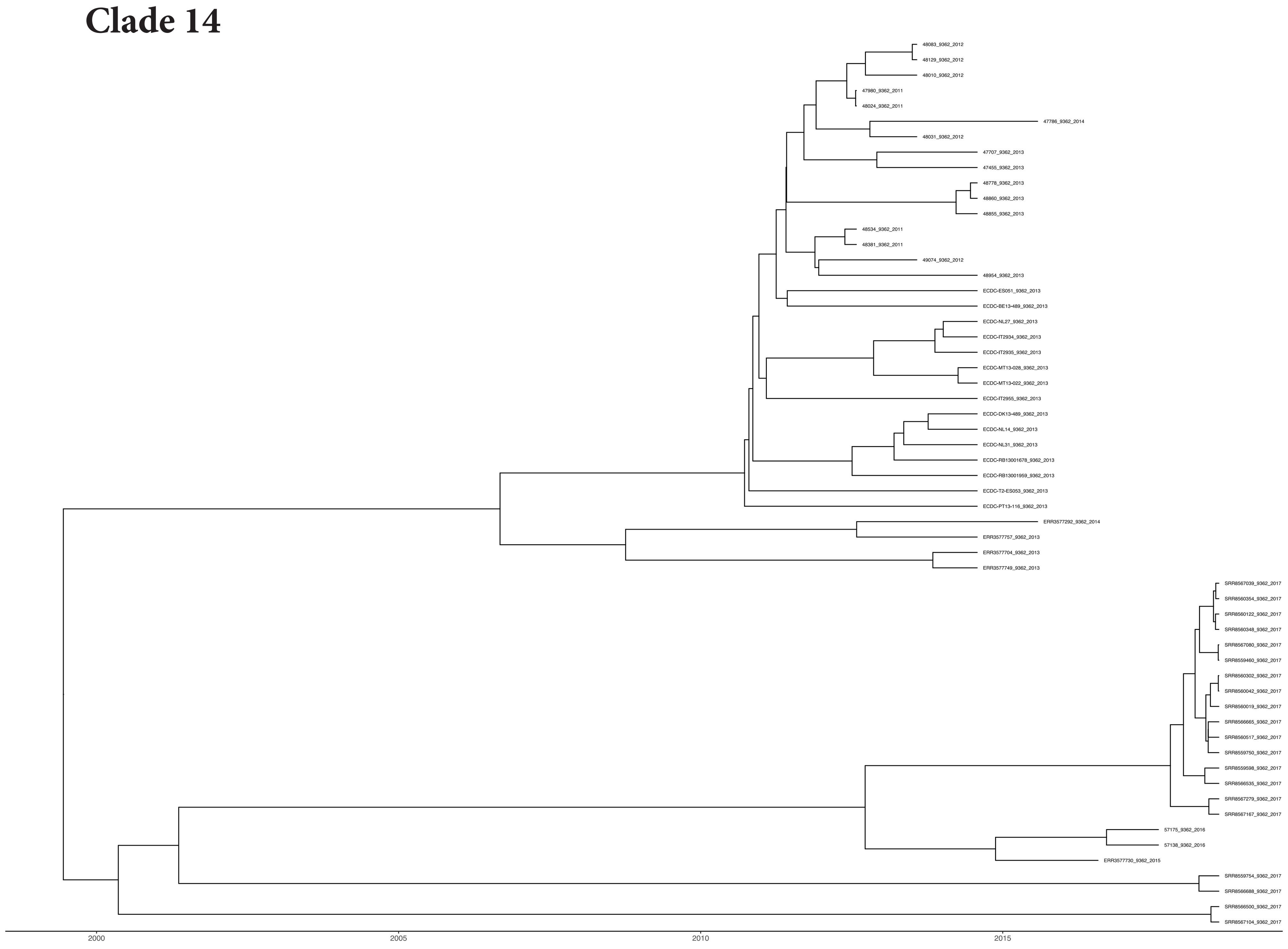


**Major MLST - 1580**
**Majority of Isolates are from Europe, North America and Oceania**
**Most recent year of isolate sampling - 2017**

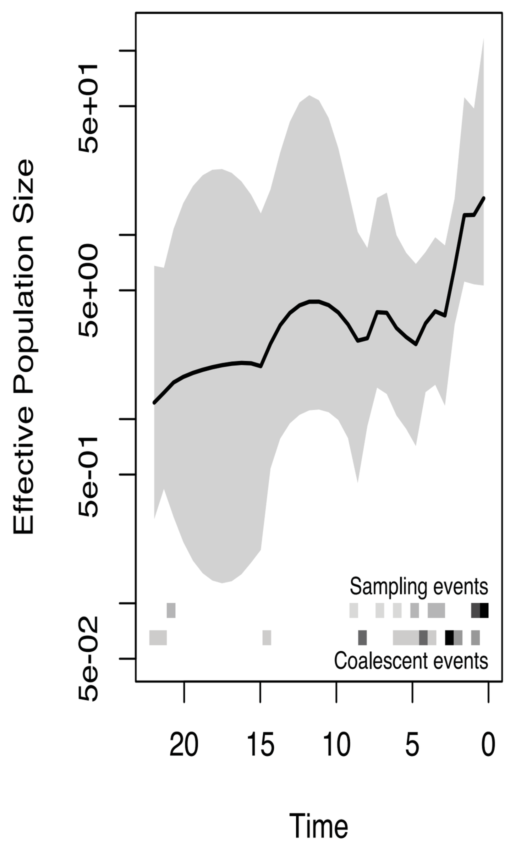


#### Clade 15

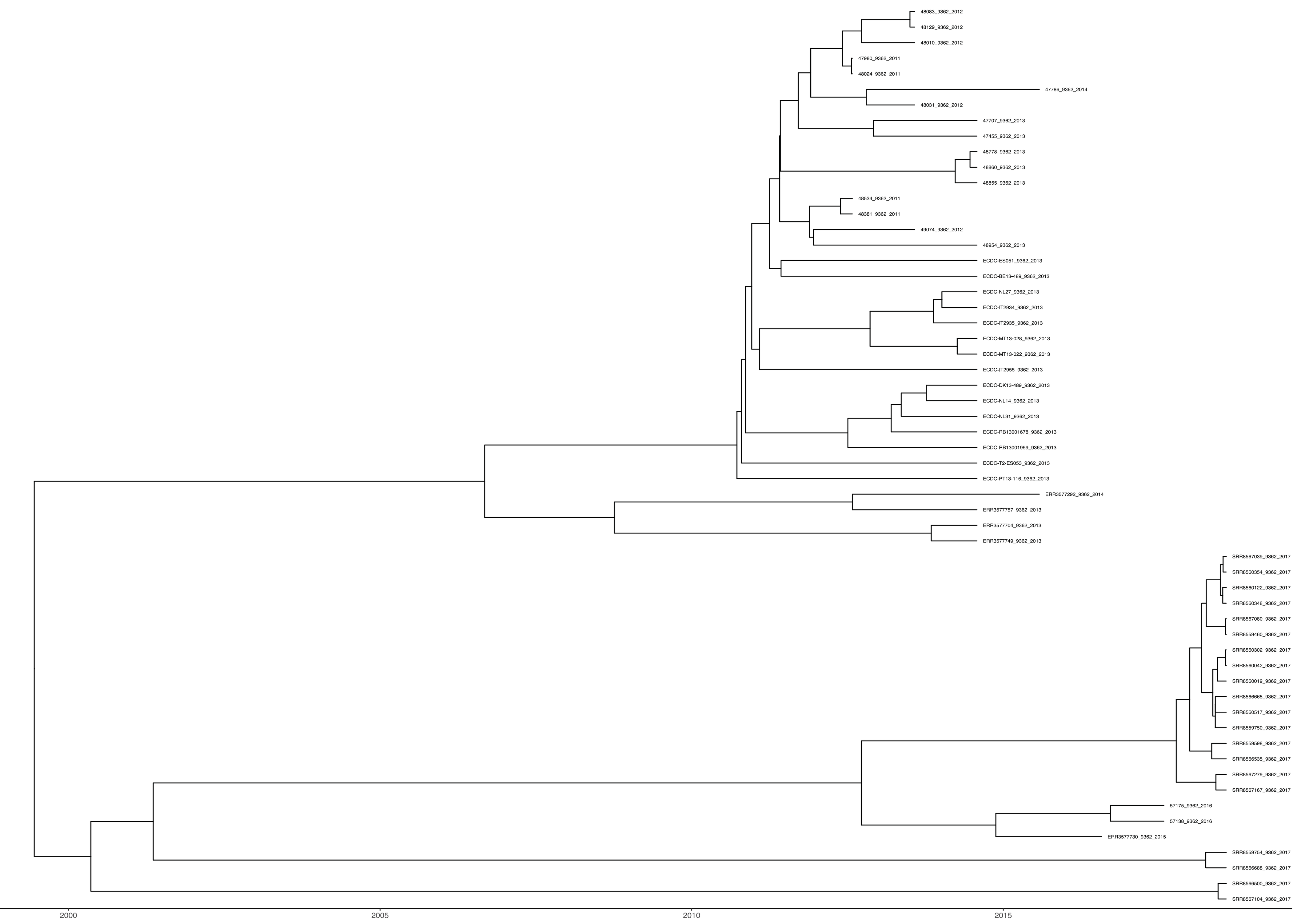


**Major MLST - 9362**
**Majority of Isolates are from Europe and Oceania**
**Most recent year of isolate sampling - 2017**

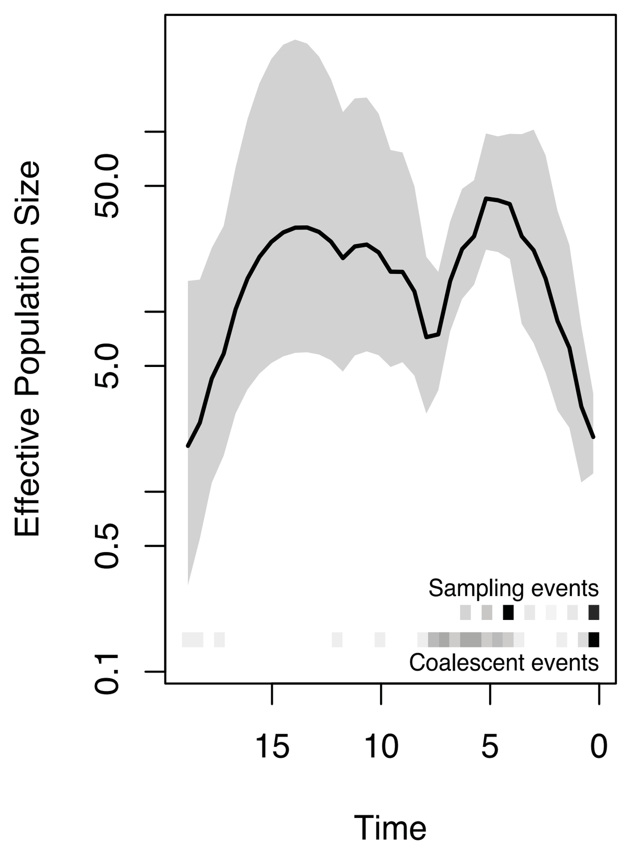

**Clade 16**

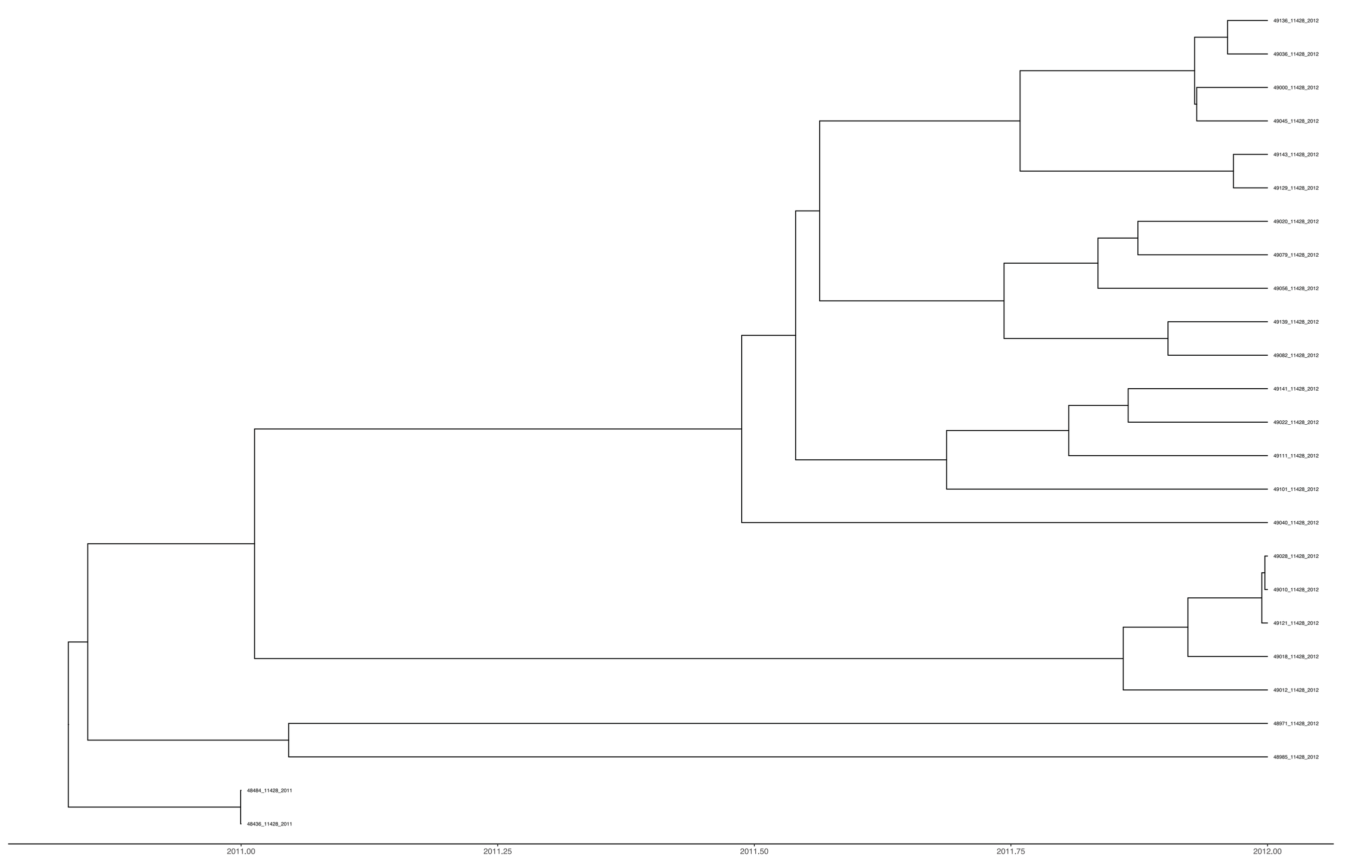

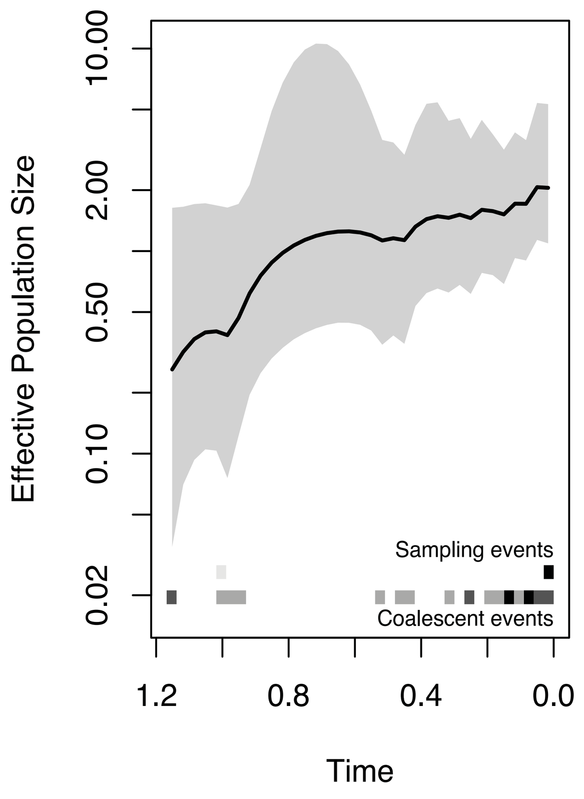


**Major MLST is 11428**
**Majority of the isolates are from Europe**
**Most recent year of isolate sampling - 2012**

**Clade 17**

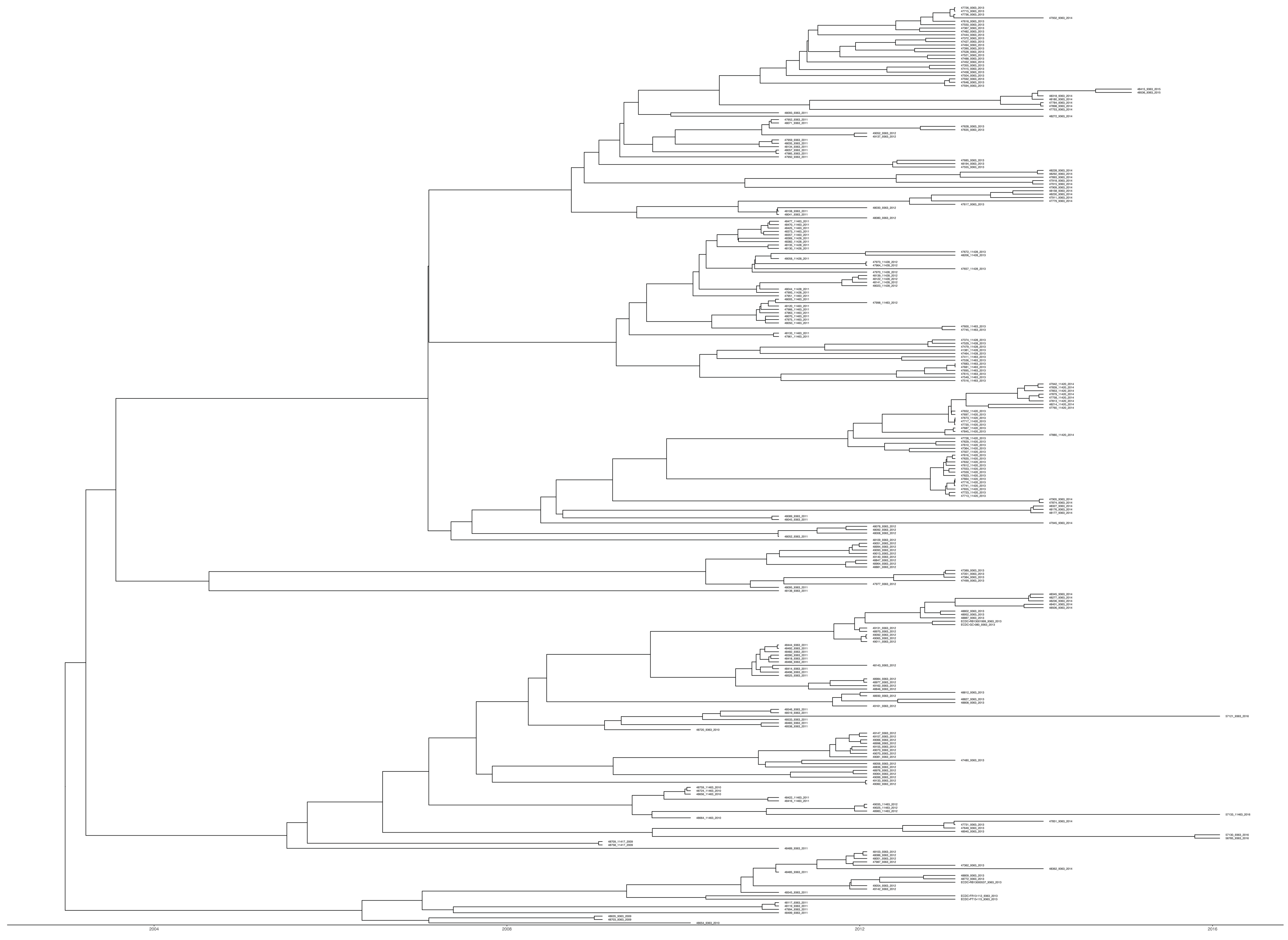

**Major MLSTs are 11428, 9363 and 11420**
**Majority of the isolates are from Europe**
**Most recent year of isolate sampling - 2016**

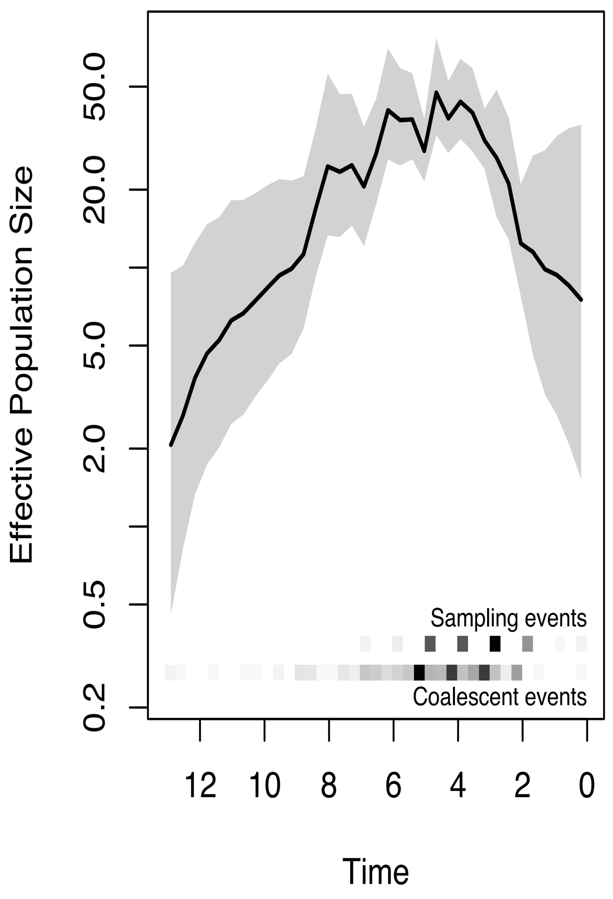


Clade 18

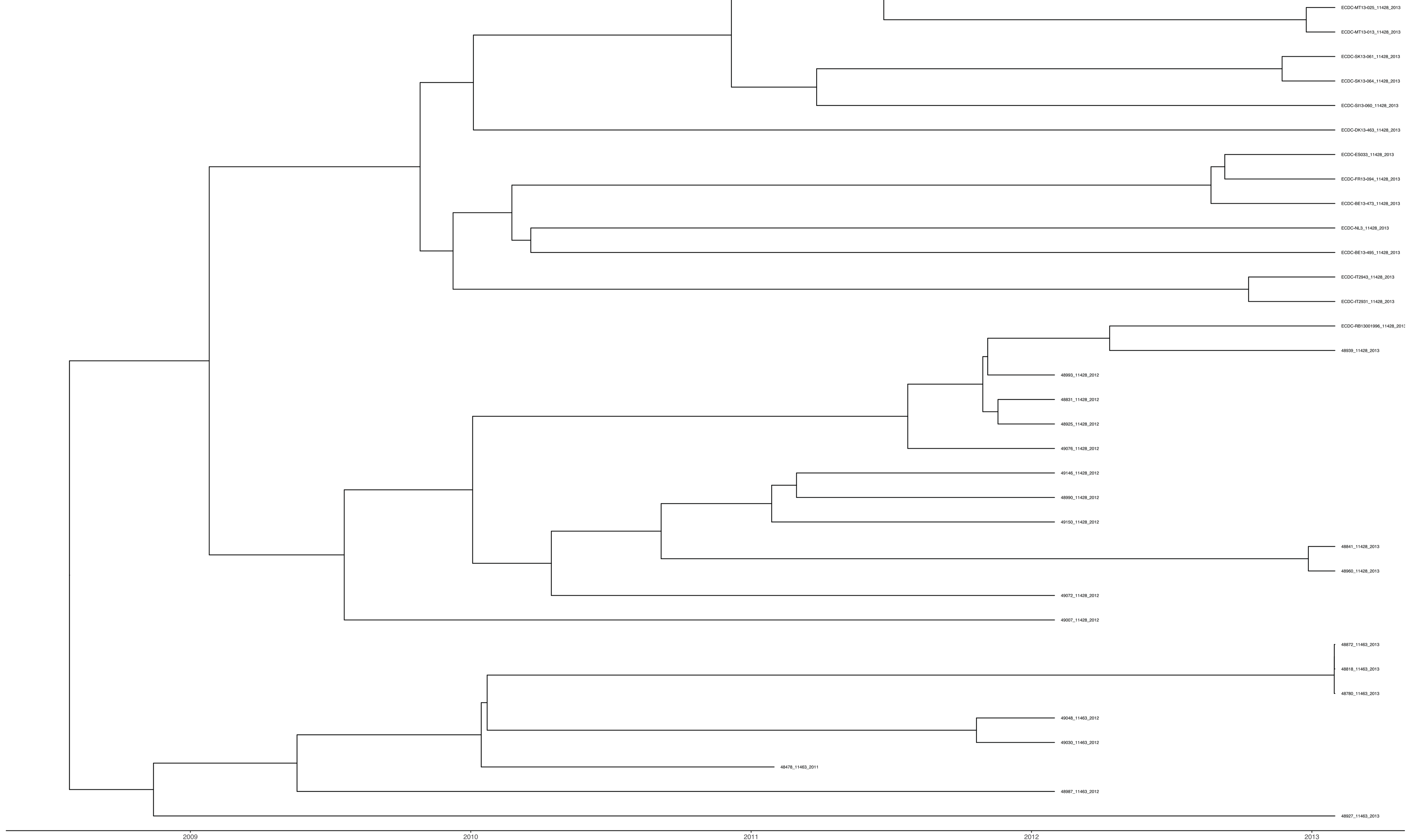

**Major MLST is 11428**
**Majority of the isolates are from Europe**
**Most recent year of isolate sampling - 2013**

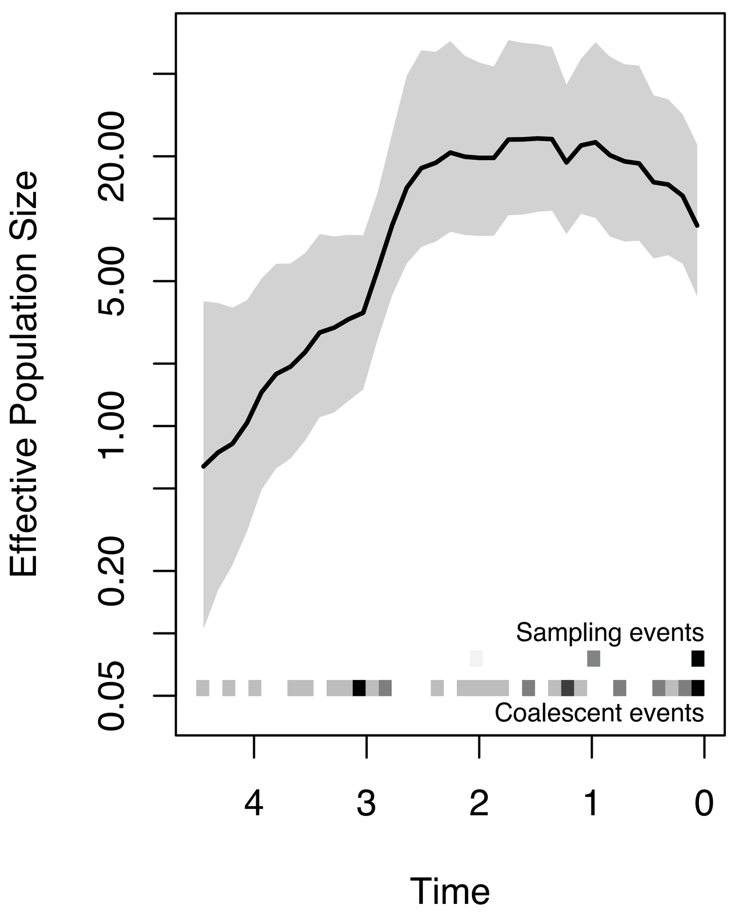

#### Clade 19

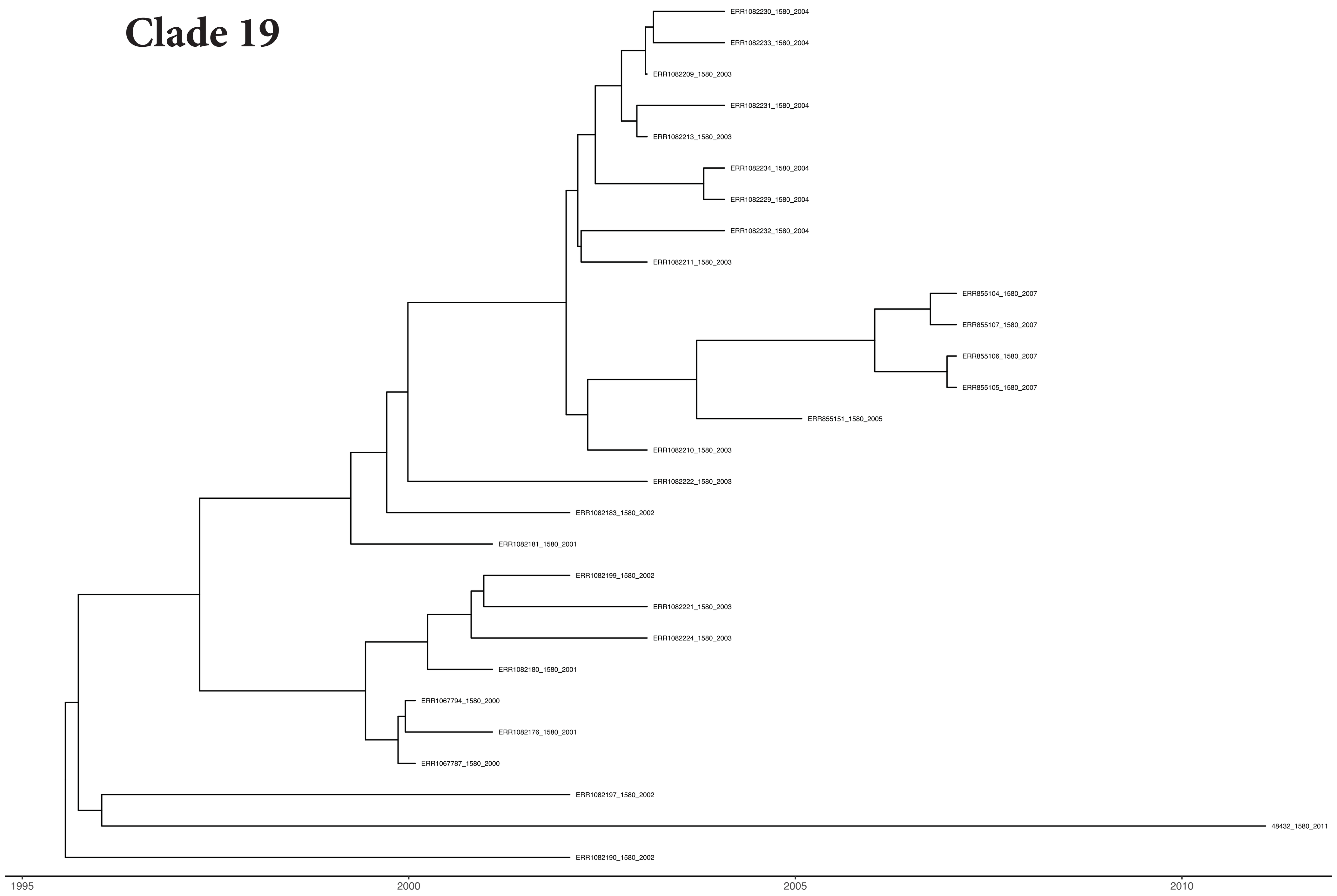


**Major MLST is 1580**
**Majority of the isolates are from Europe**
**Most recent year of isolate sampling - 2011**

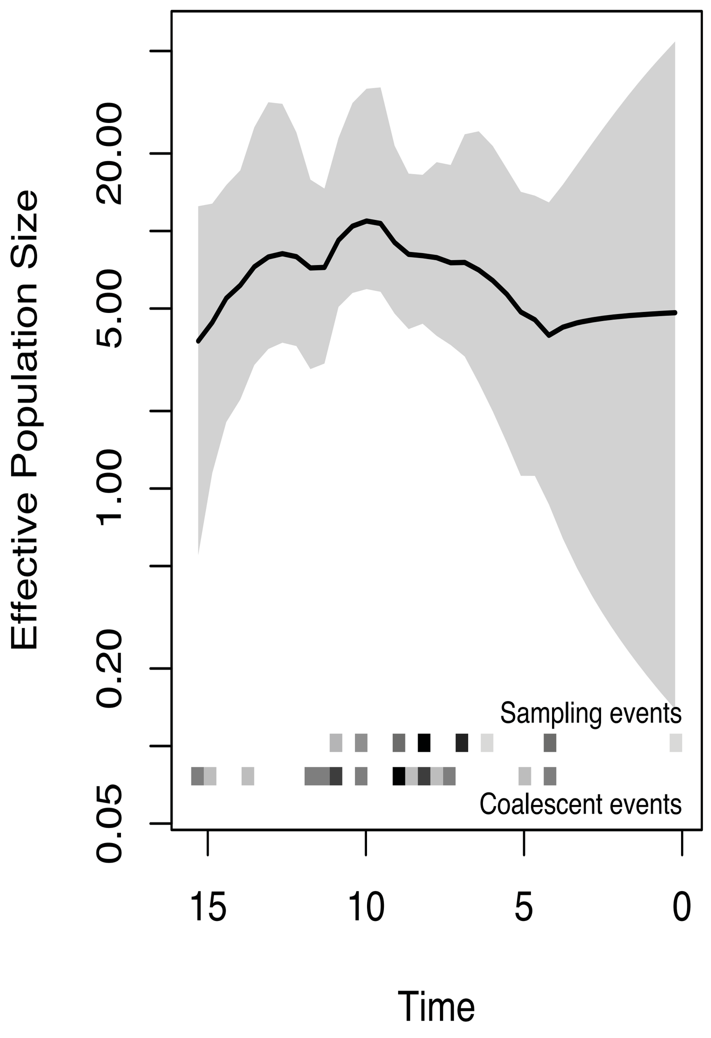


#### Clade 20

**Major MLST is 1580**
**Majority of the isolates are from North America**
**Most recent year of isolate sampling - 2008**

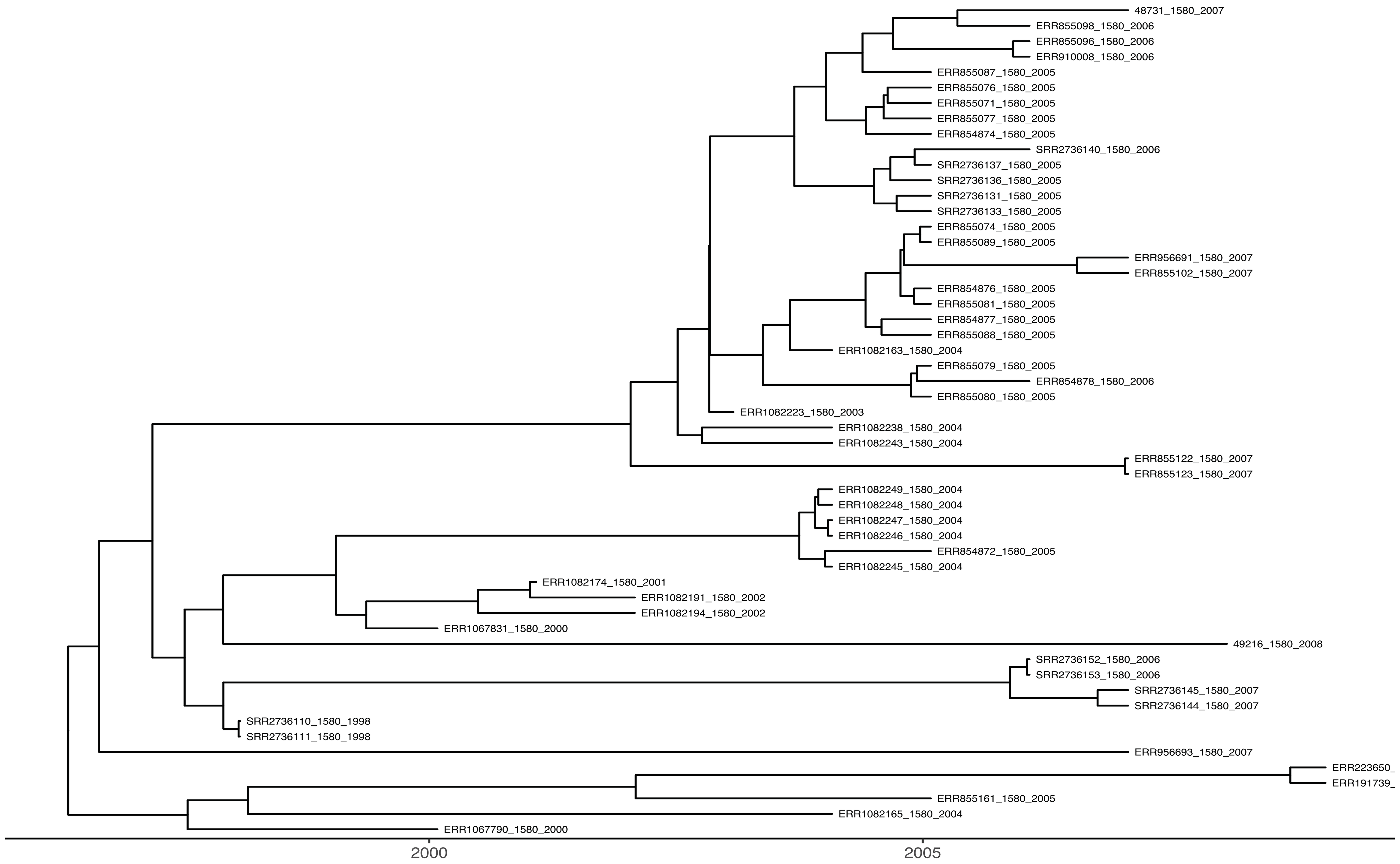


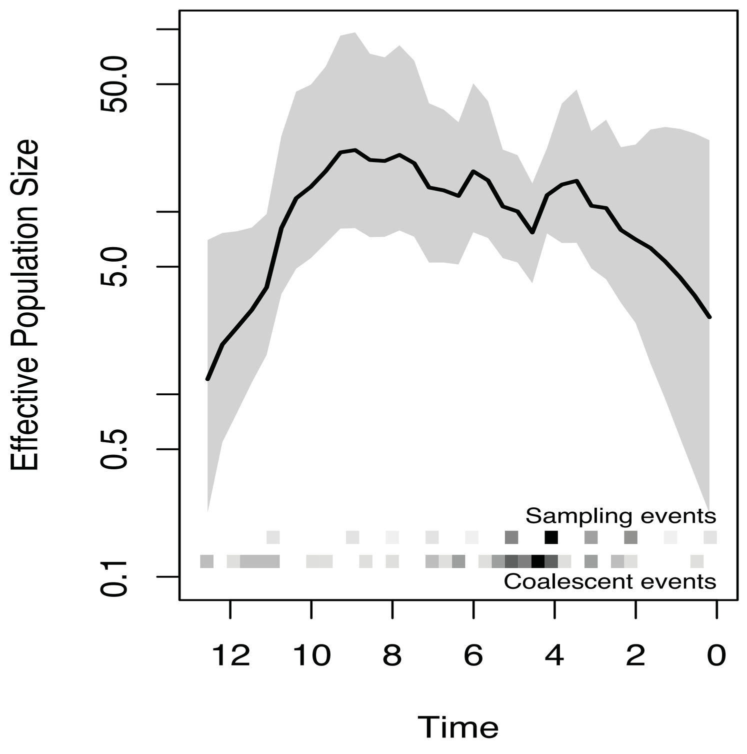


#### Clade 21

**Major MLST is 8126**
**Majority of the isolates are from Europe**
**Most recent year of isolate sampling - 2013**

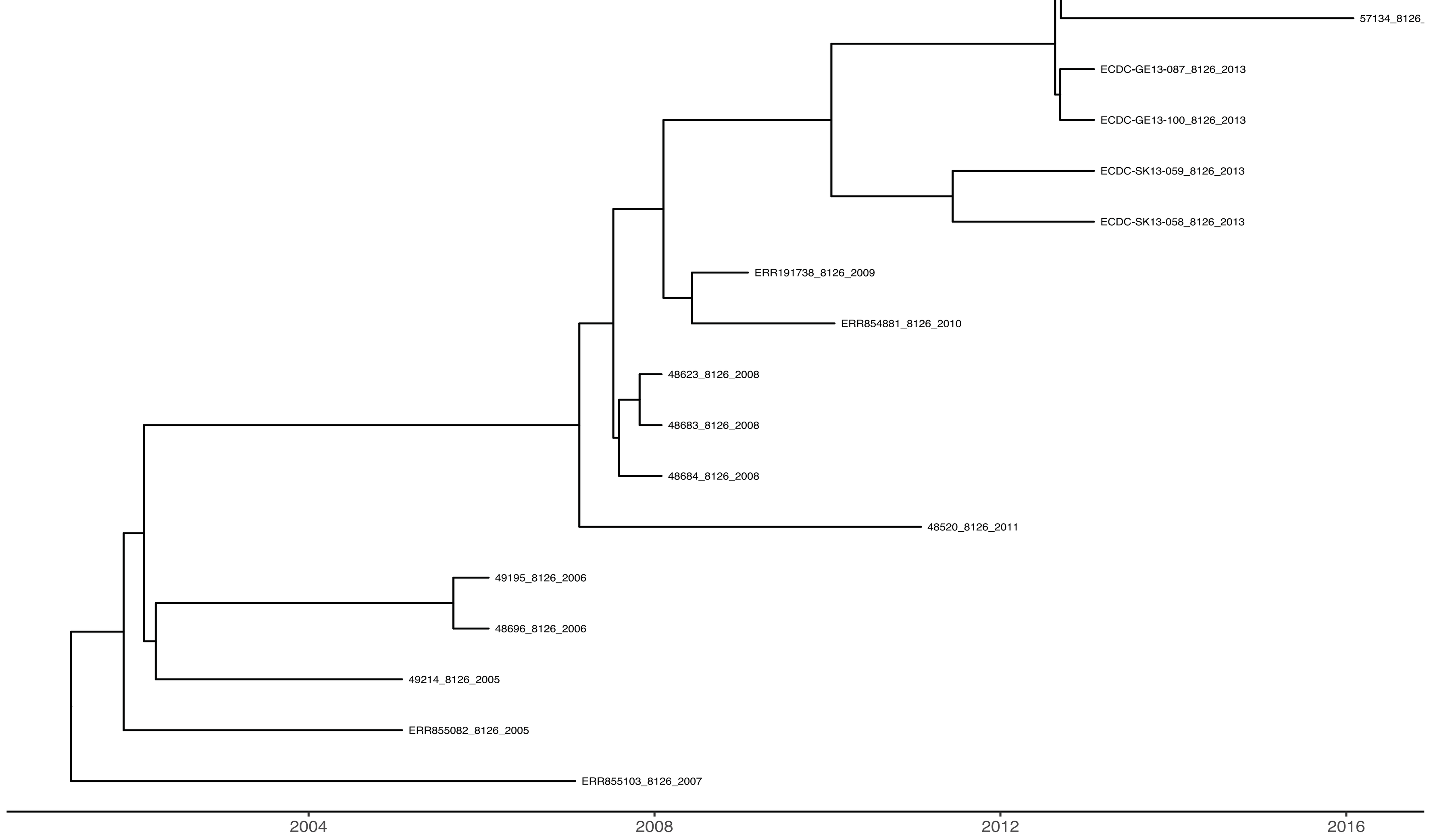


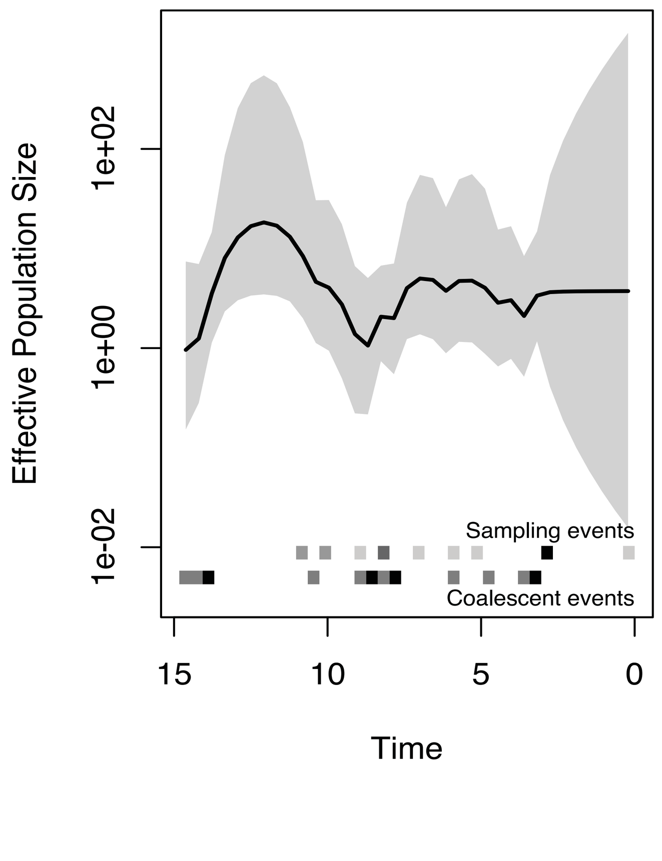

#### Clade 22

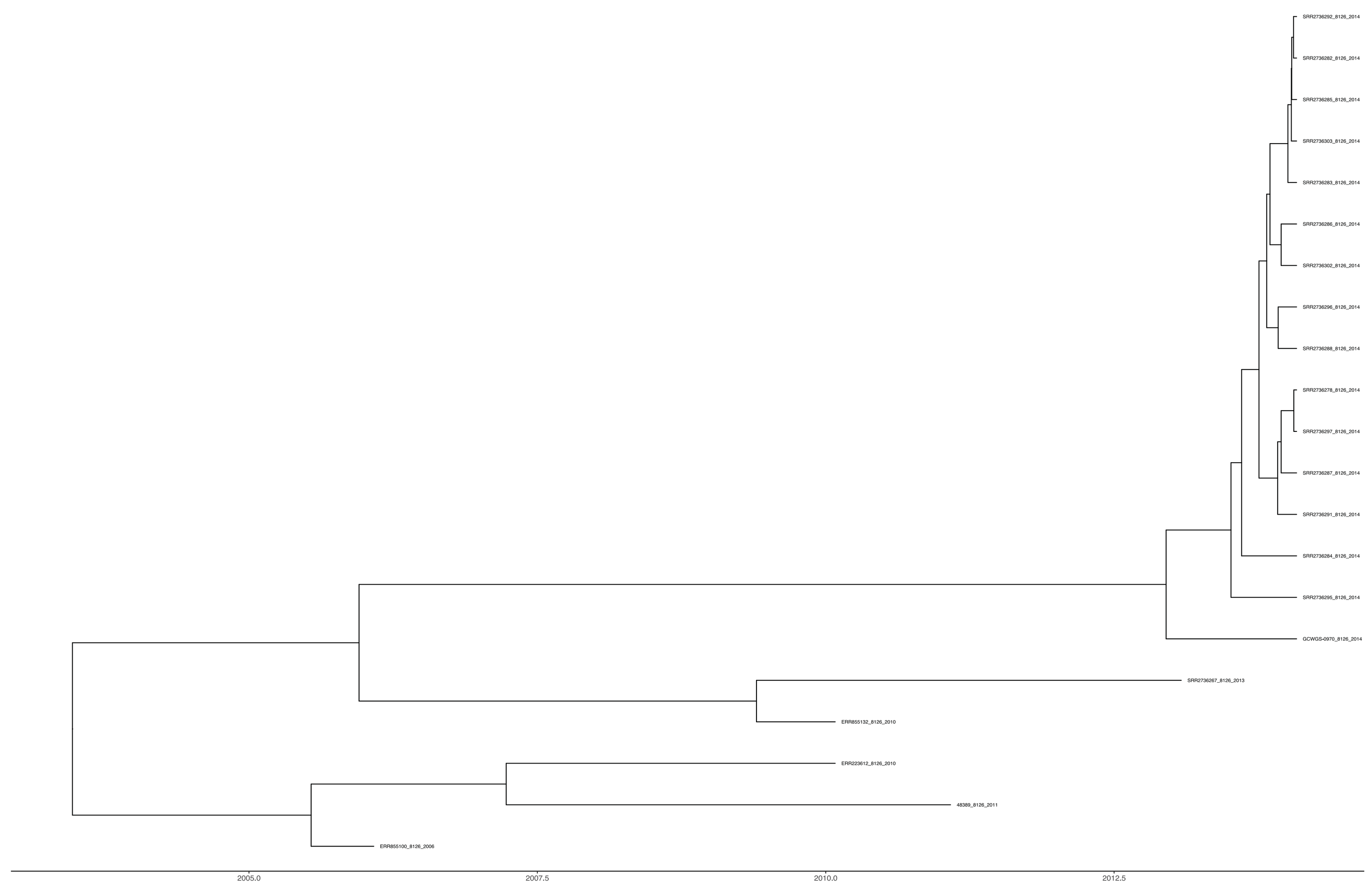

**Major MLST is 8126**
**Majority of the isolates are from North America**
**Most recent year of isolate sampling - 2014**

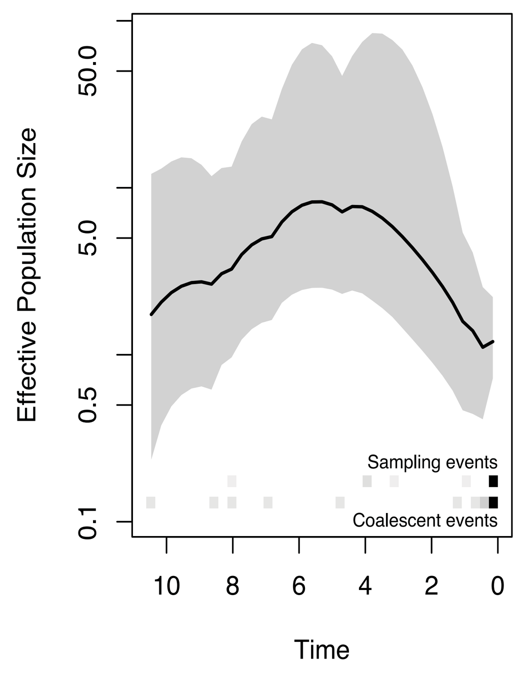


#### Clade 23

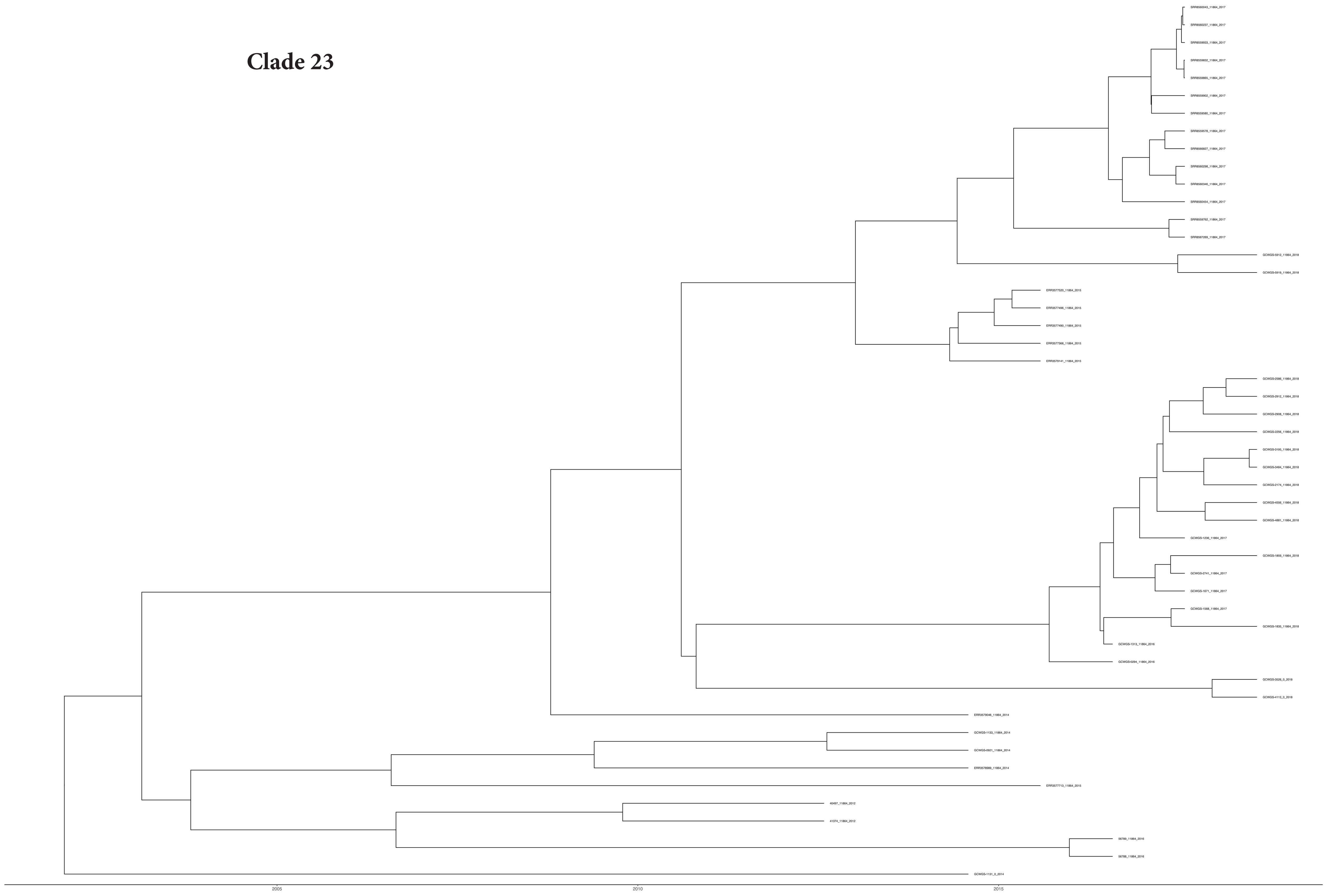

**Major MLST is 11864**
**Majority of the isolates are from North America, Europe and Oceania**
**Most recent year of isolate sampling - 2008**

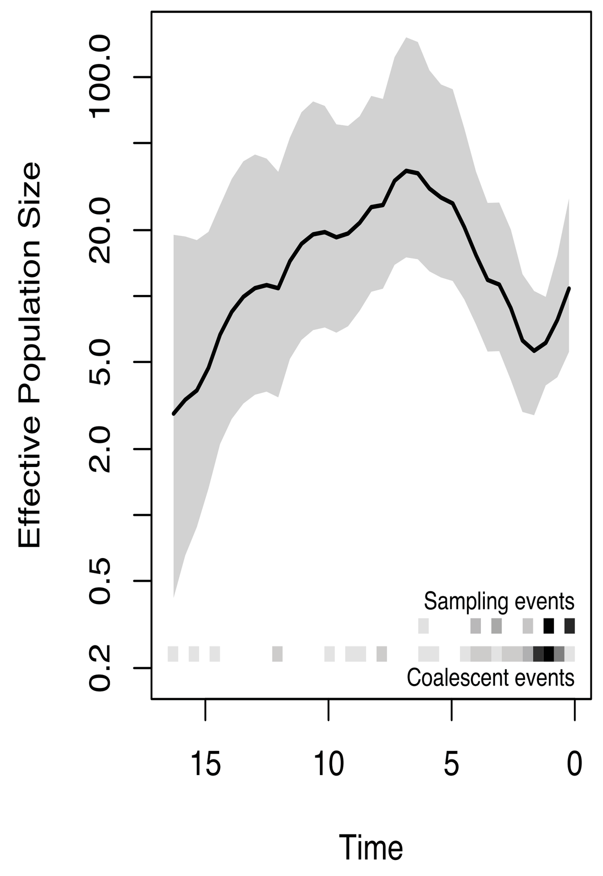


#### Clade 24

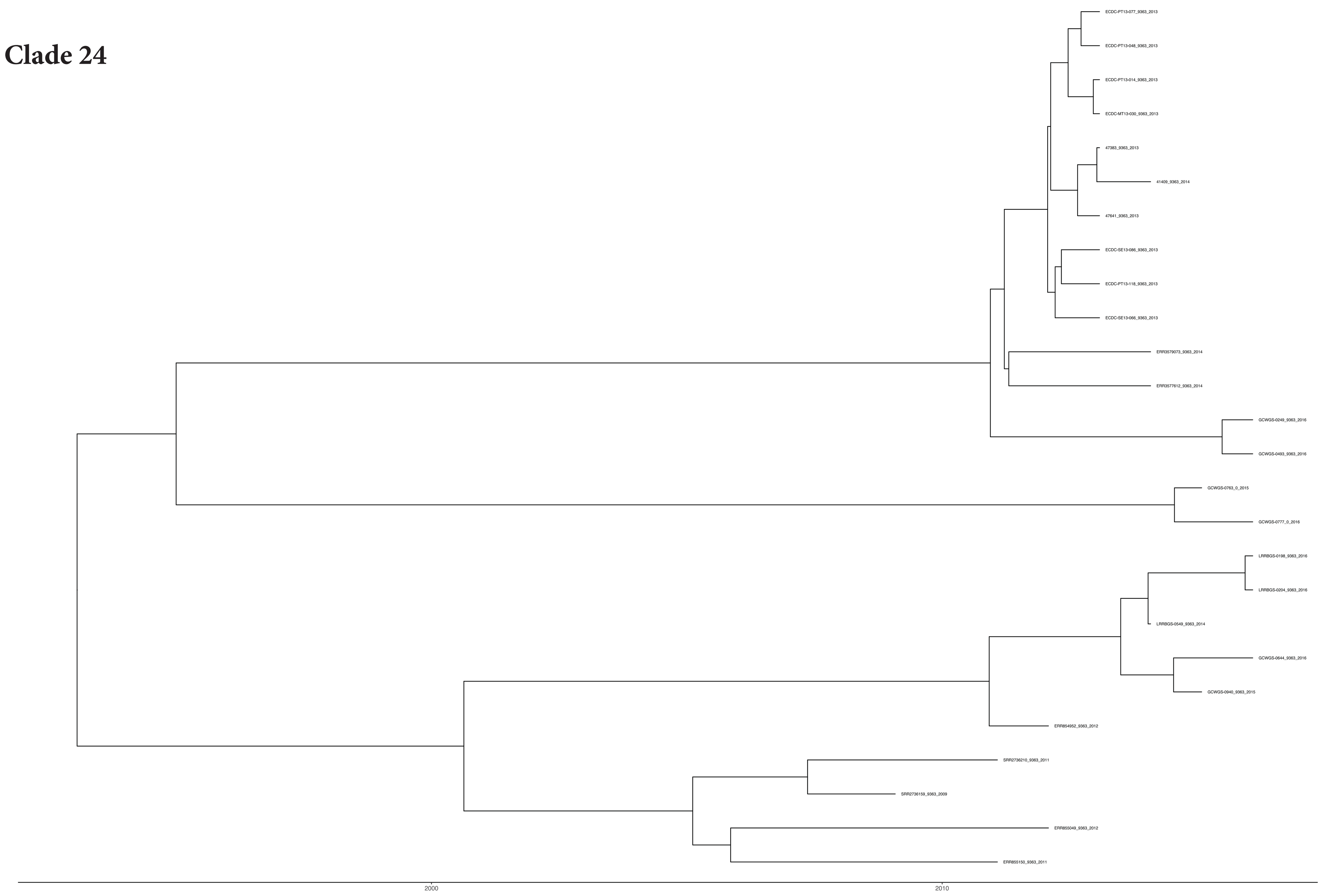

**Major MLST is 9363**
**Majority of the isolates are from North America and Europe**
**Most recent year of isolate sampling - 2016**

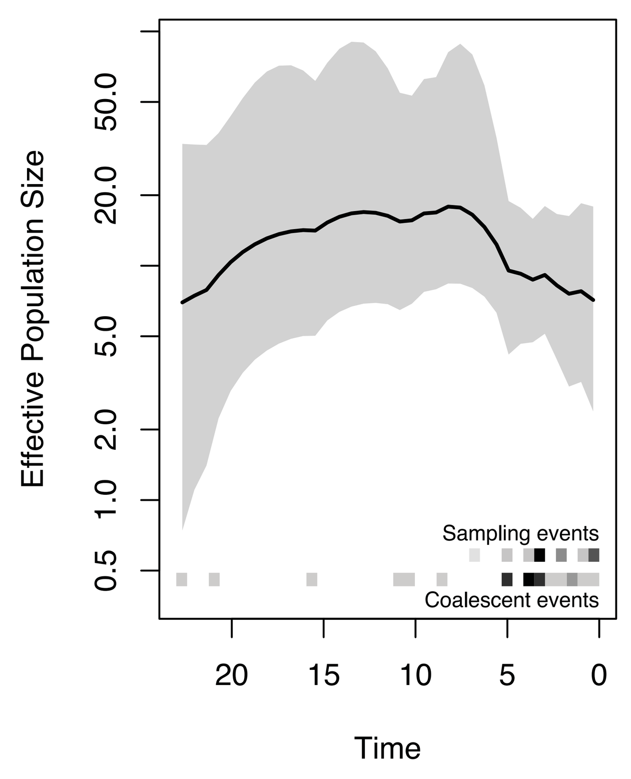

**Clade 25**

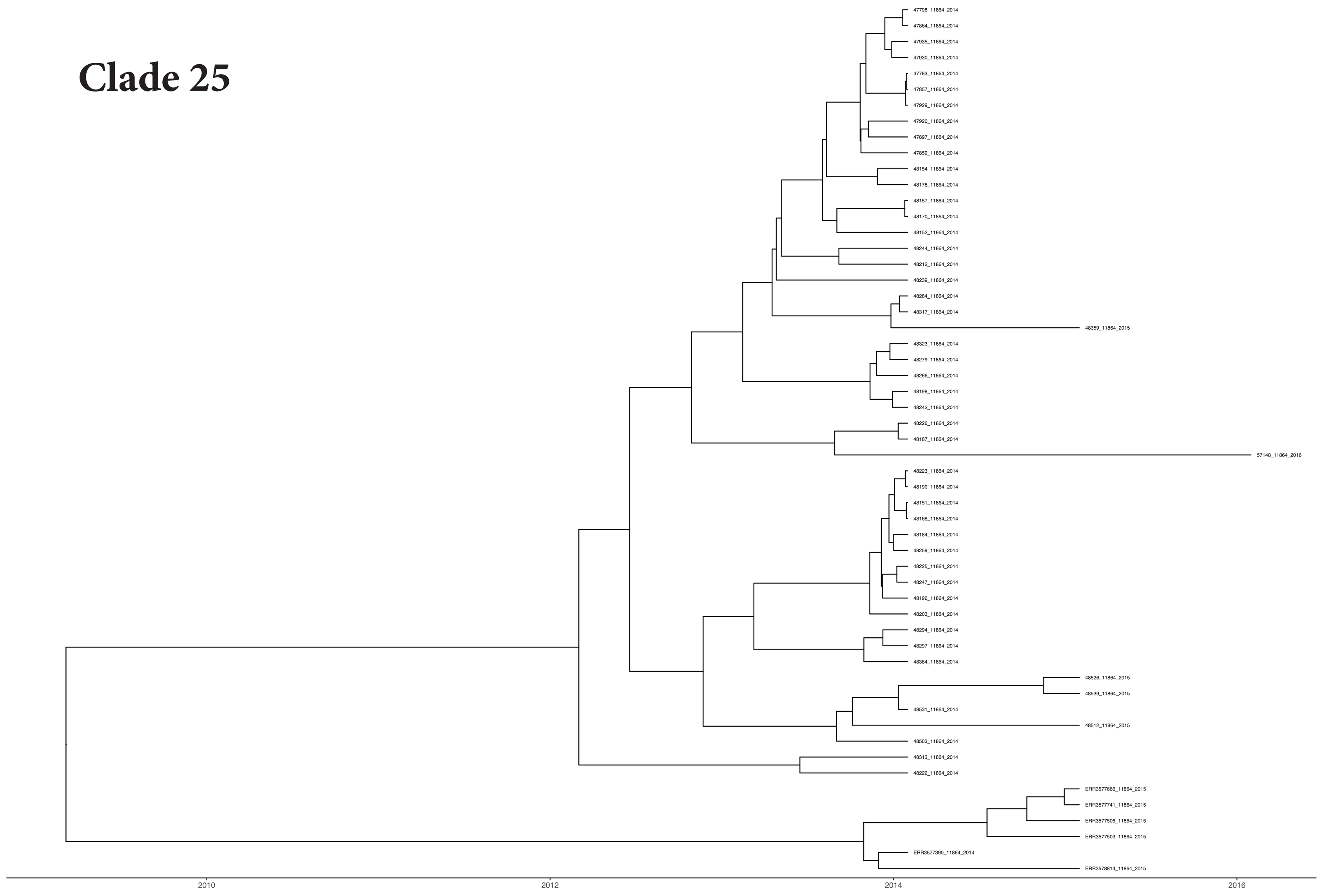


**Major MLST is 11864**
**Majority of the isolates are from Europe**
**Most recent year of isolate sampling - 2016**

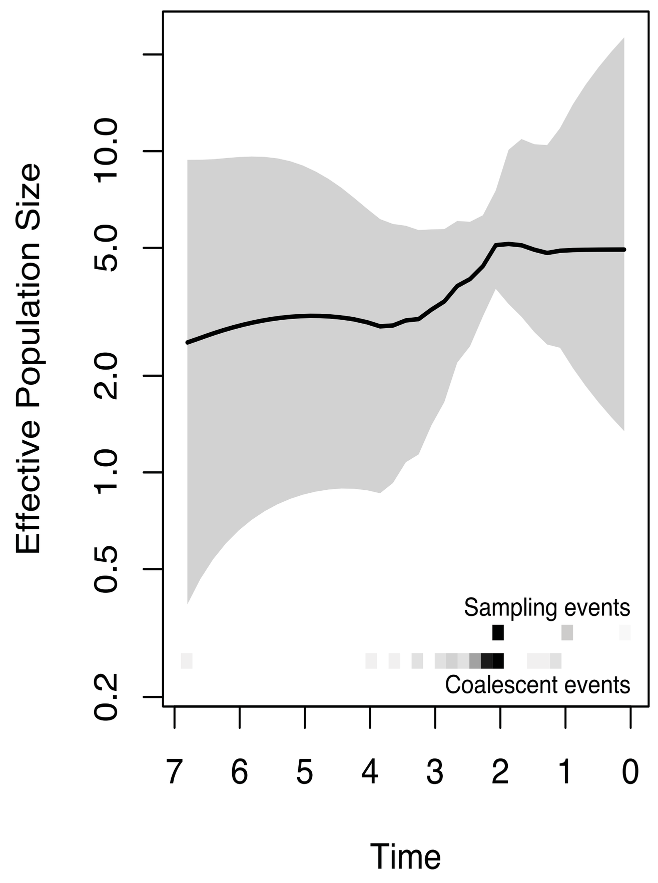


**Clade 26**

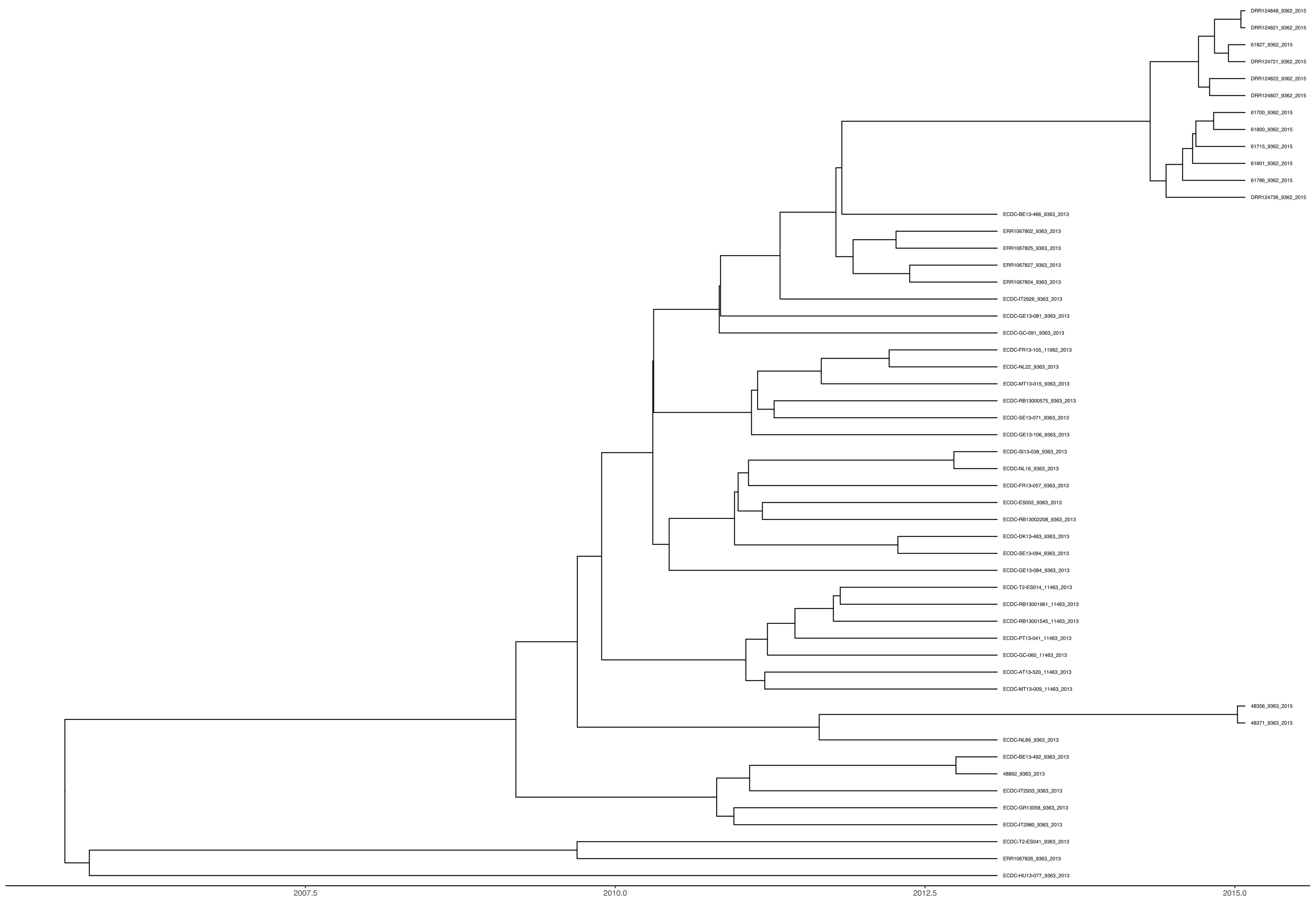


**Major MLST is 9363**
**Majority of the isolates are from Europe and Asia**
**Most recent year of isolate sampling - 2015**

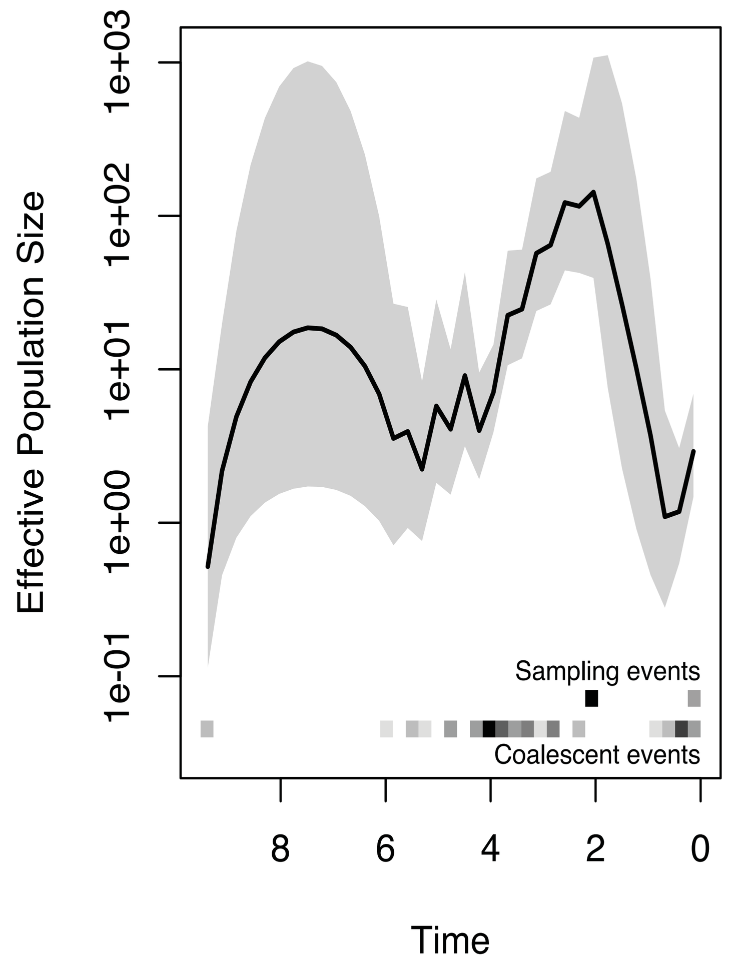


**Clade 27**

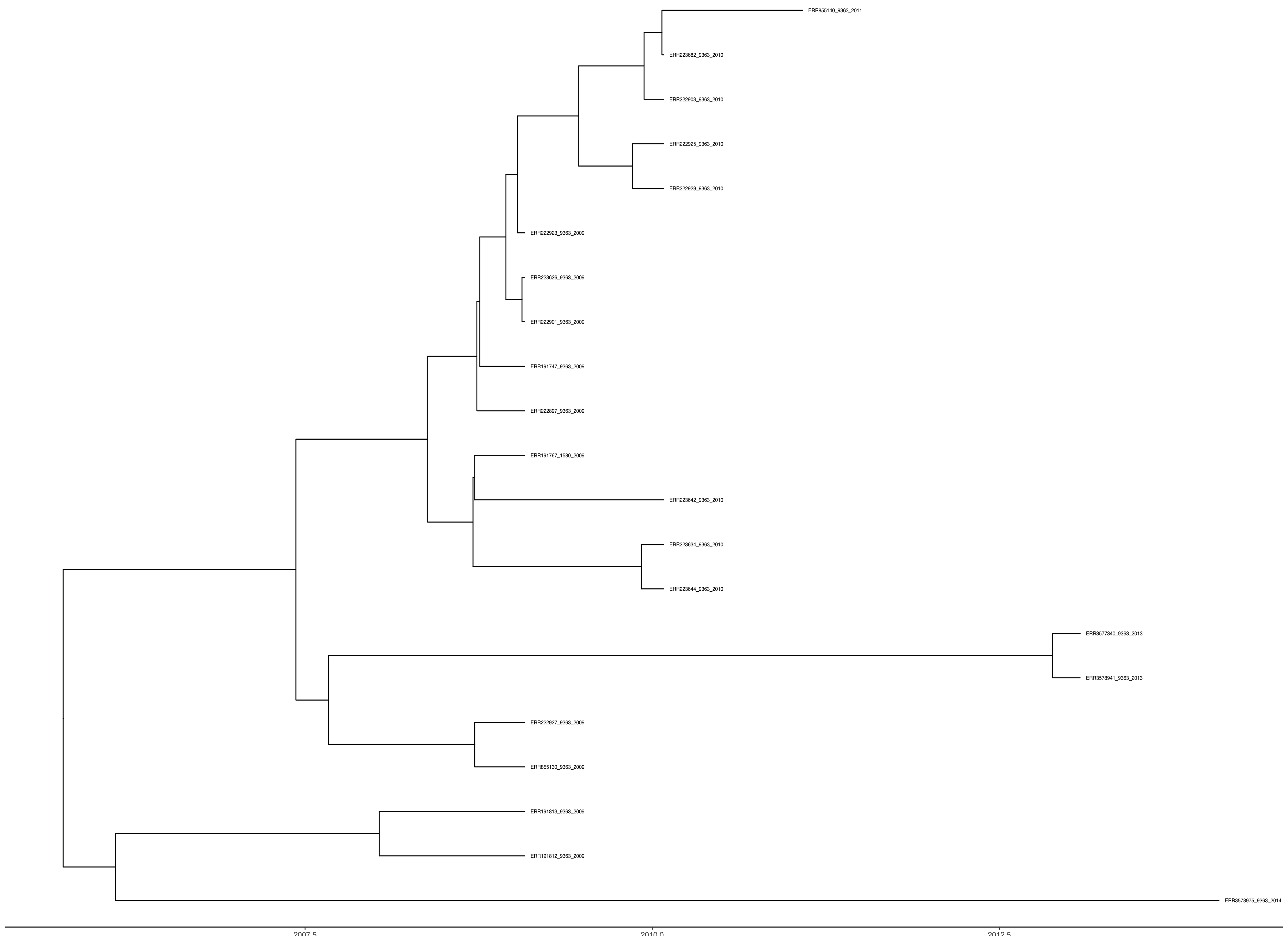


**Major MLST is 9363**
**Majority of the isolates are from North America**
**Most recent year of isolate sampling - 2014**

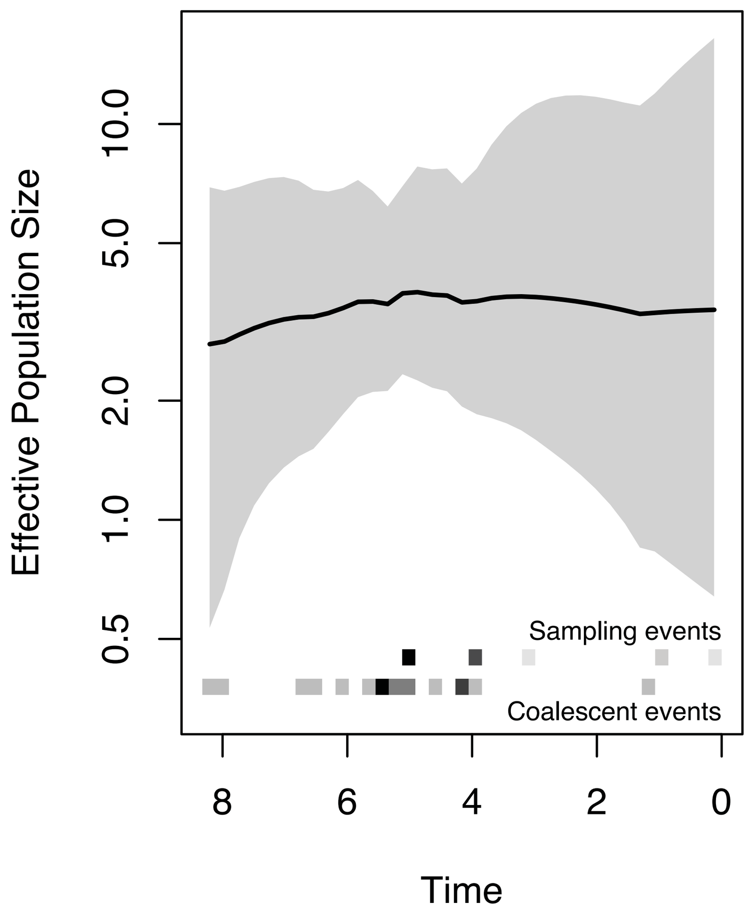

#### Clade 28

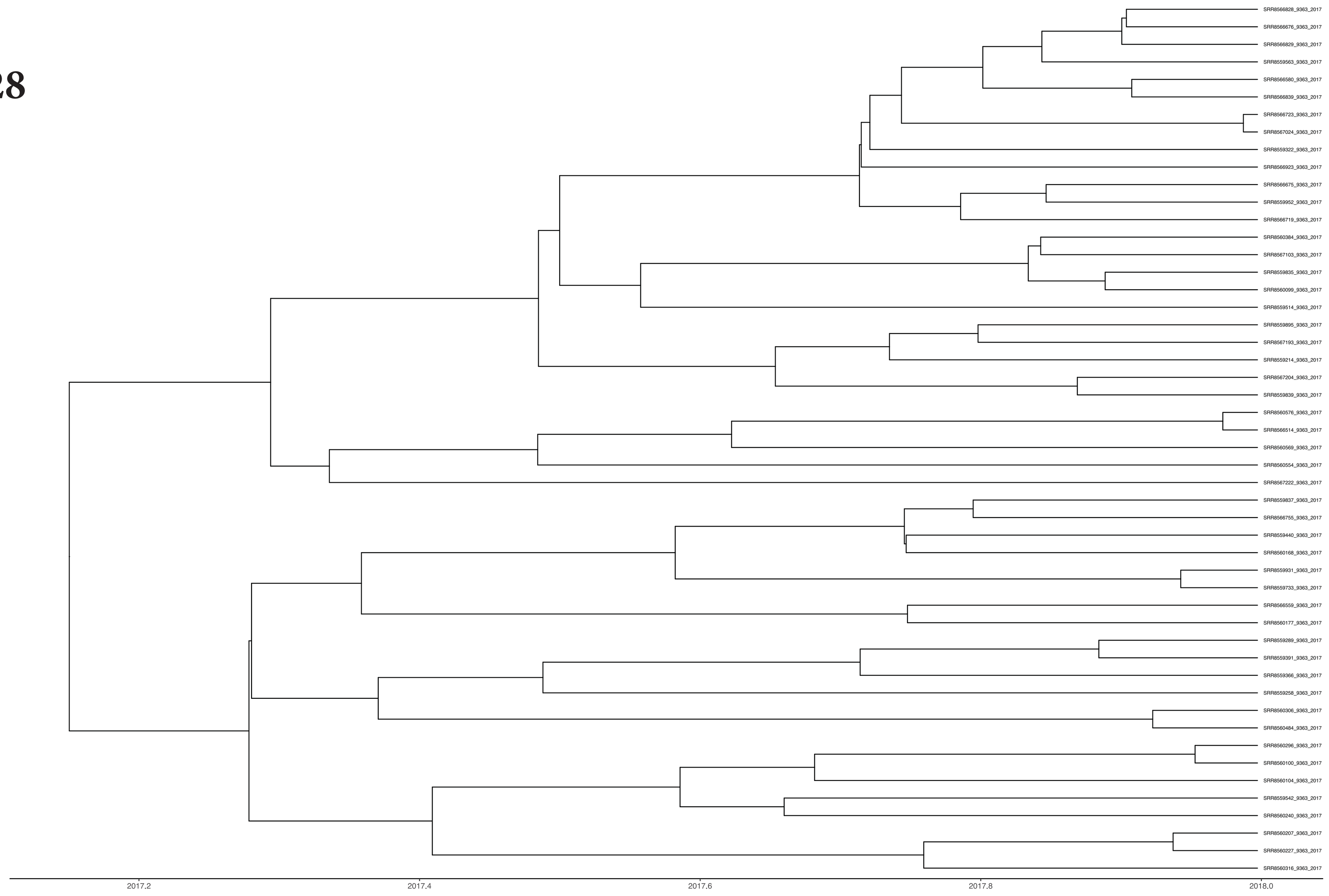

Major MLST is 9363
Majority of the isolates are from Oceania
Most recent year of isolate sampling - 2017

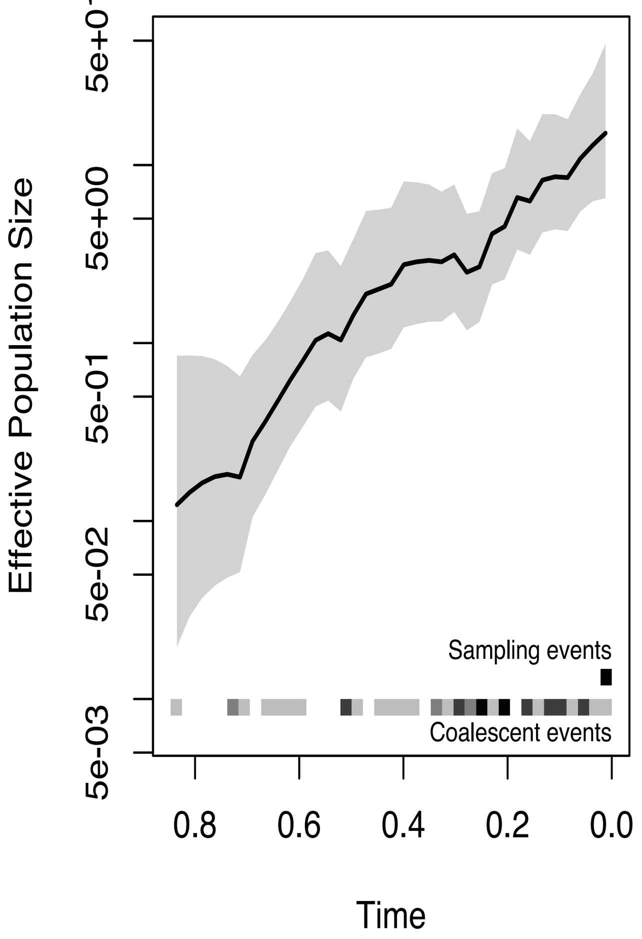


#### Clade 29

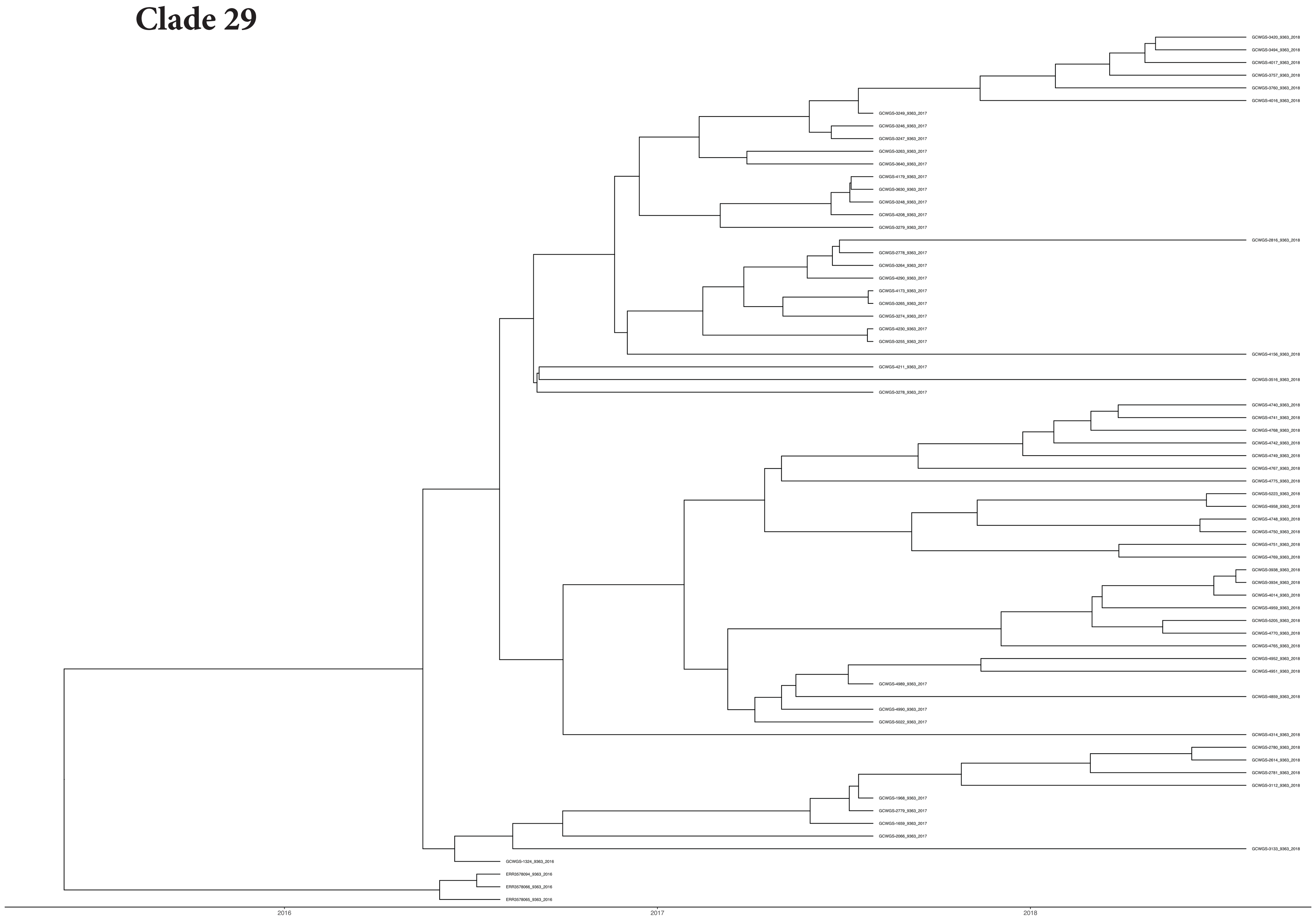

Major MLST is 9363
Majority of the isolates are from North America
Most recent year of isolate sampling - 2018

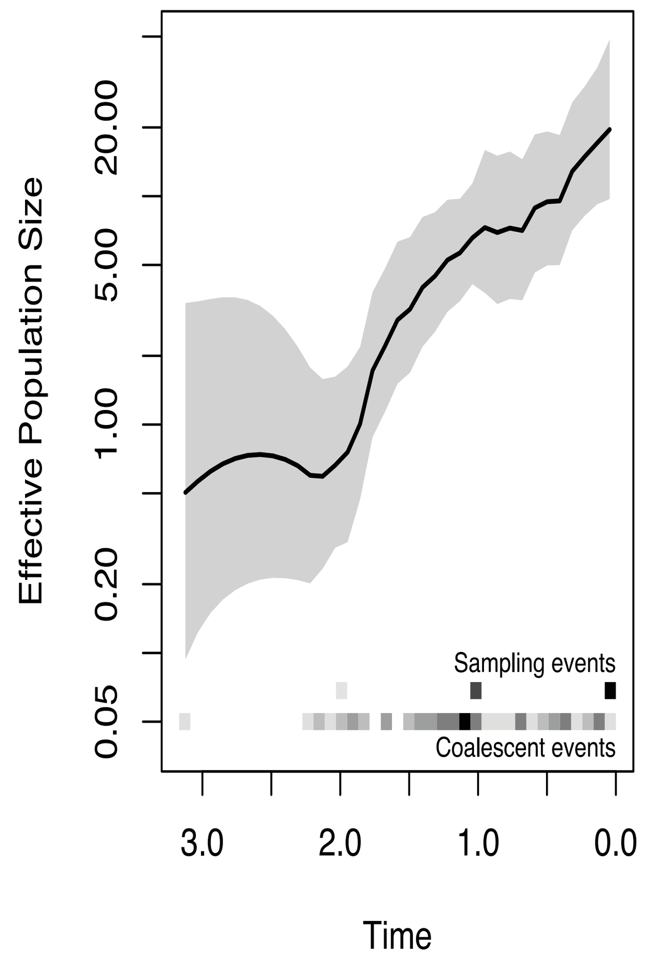


#### Clade 30

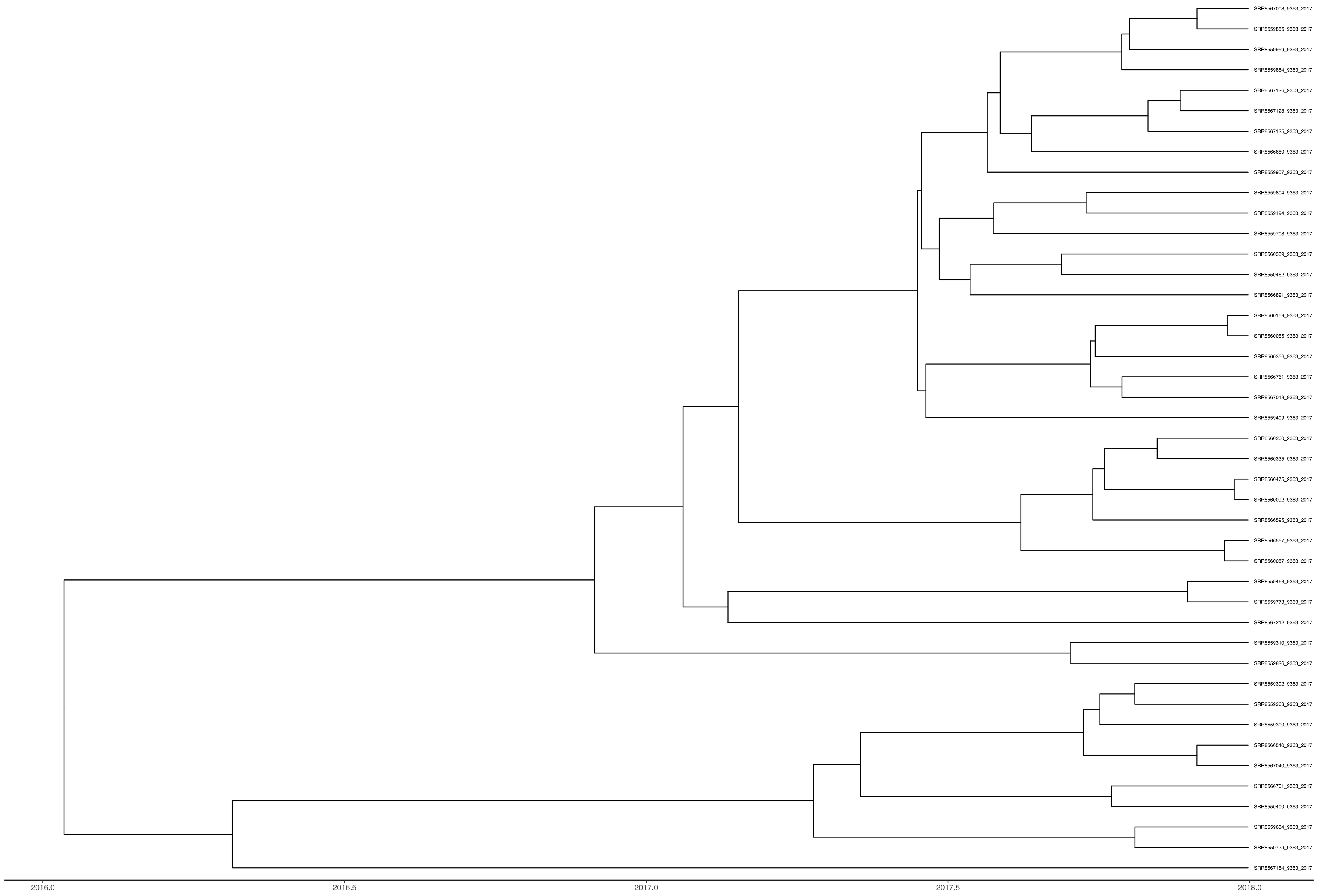

Major MLST is 9363
Majority of the isolates are from Oceania
Most recent year of isolate sampling - 2017

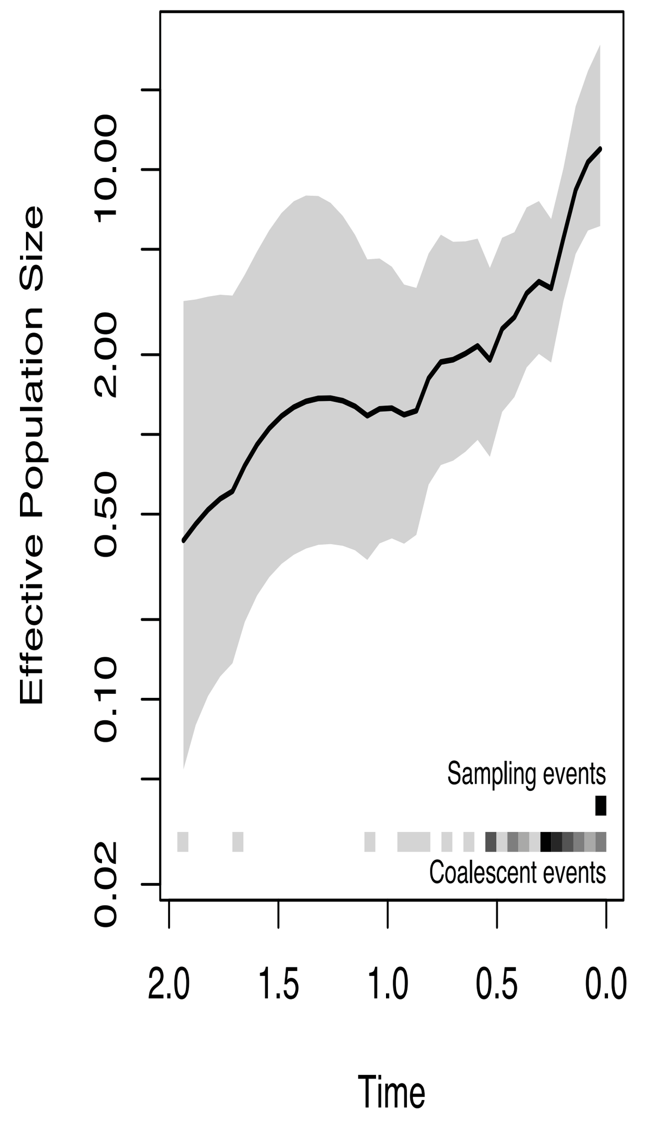

**Clade 31**

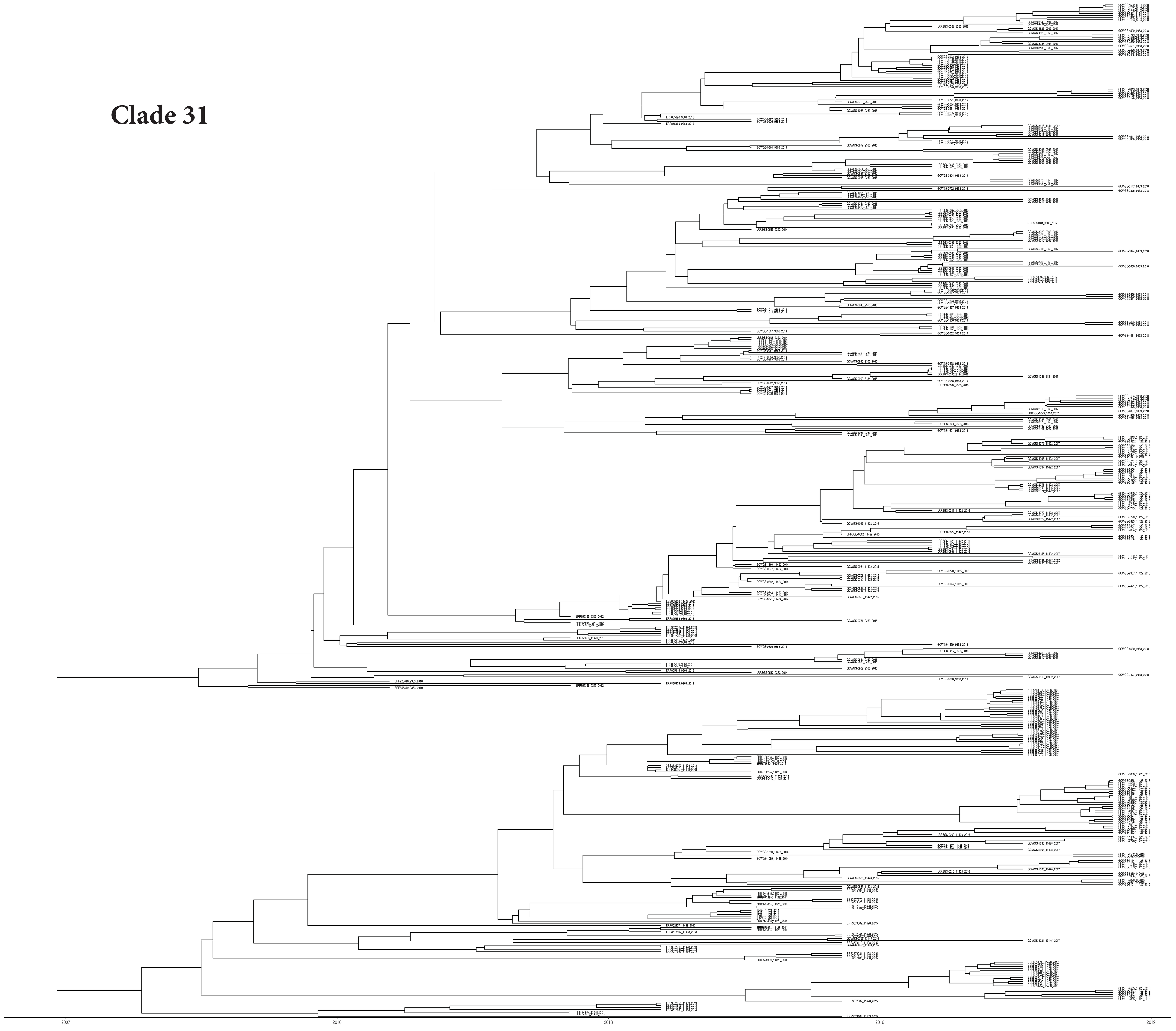

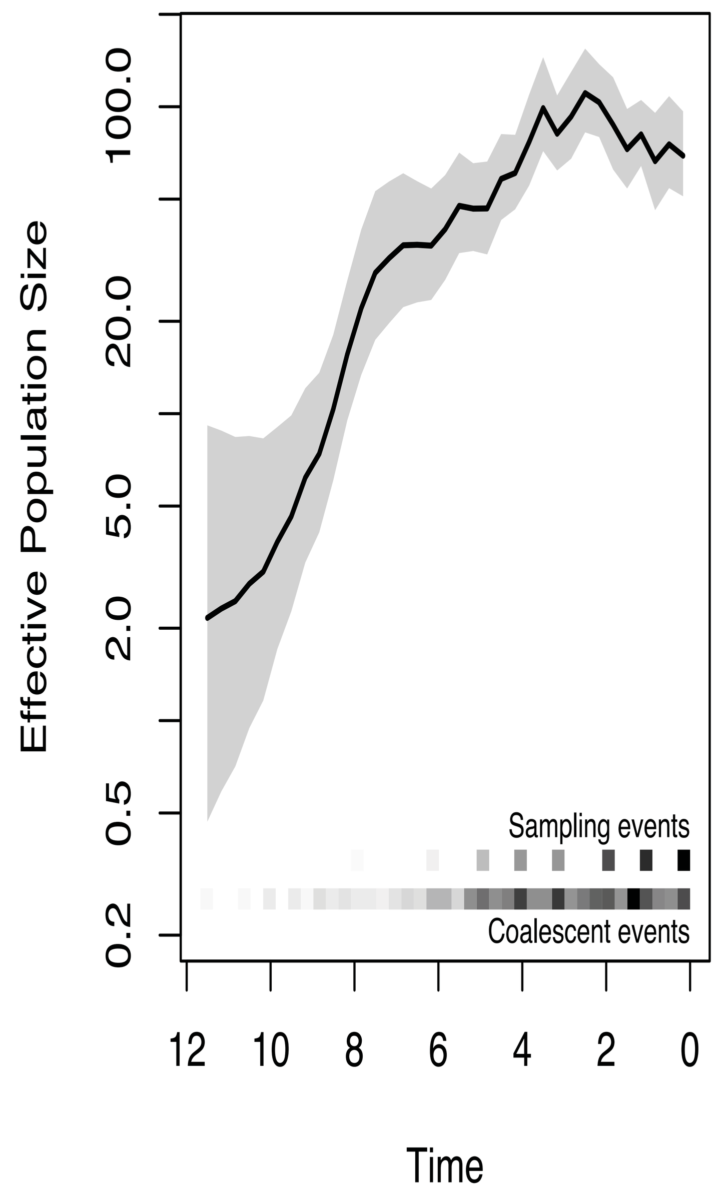


**Major MLST is 9363, 11428 and 11422**
**Majority of the isolates are from North America , Europe and Oceania.**
**Most recent year of isolate sampling - 2018**

**Major MLST is 9363**
**Majority of the isolates are from North America**
**Most recent year of isolate sampling - 2018**

**Clade 32**

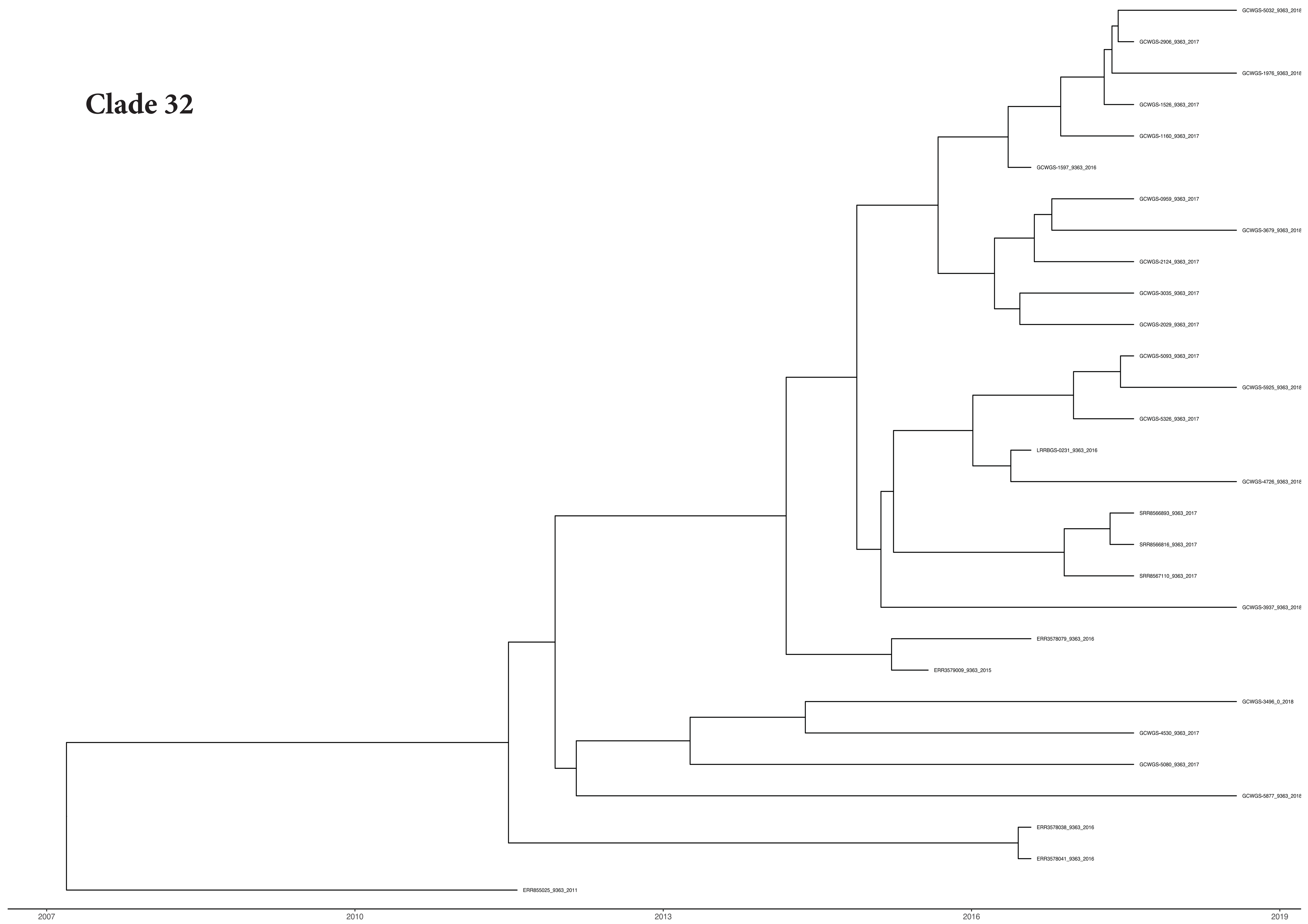


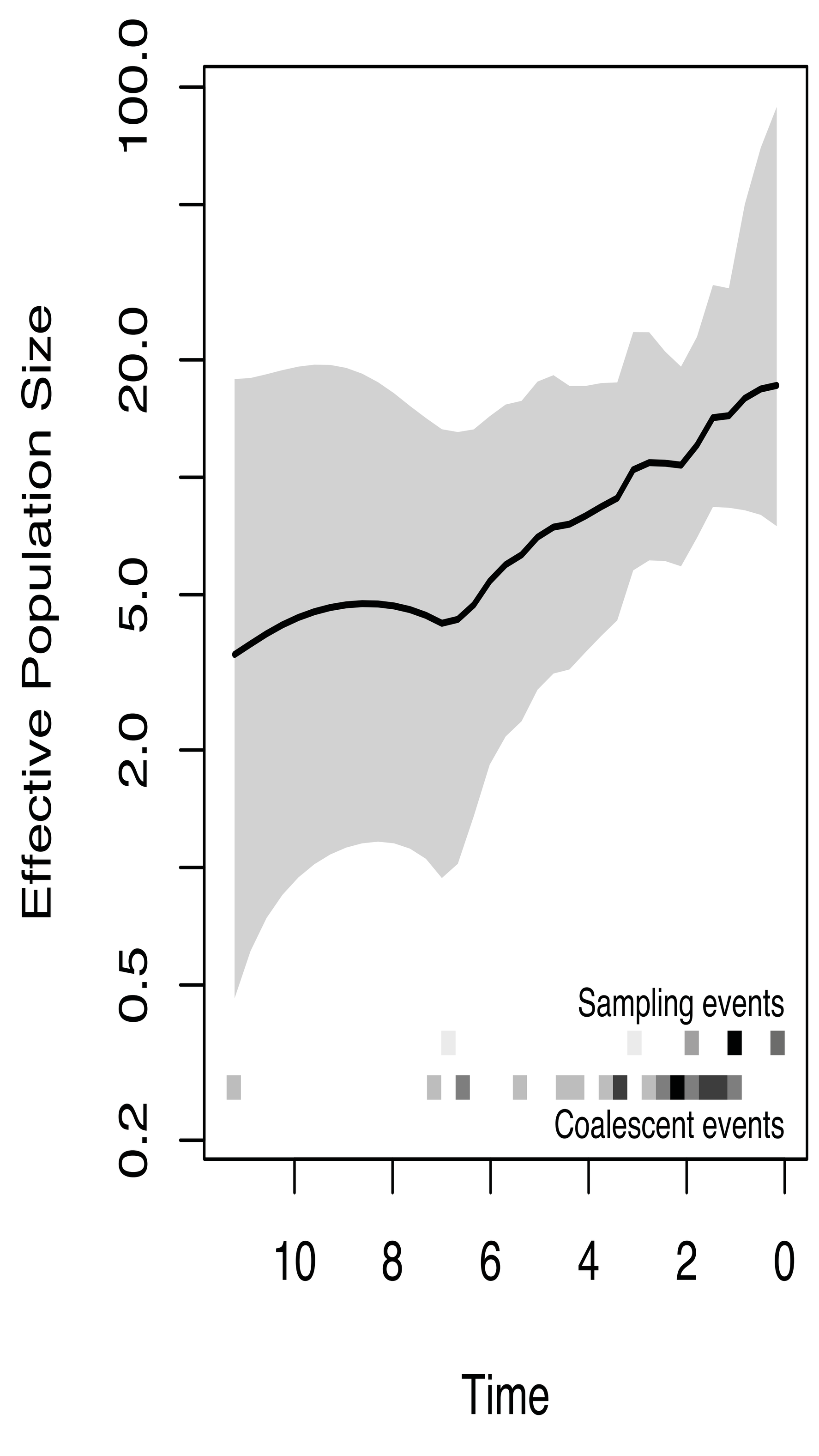

Figure S2

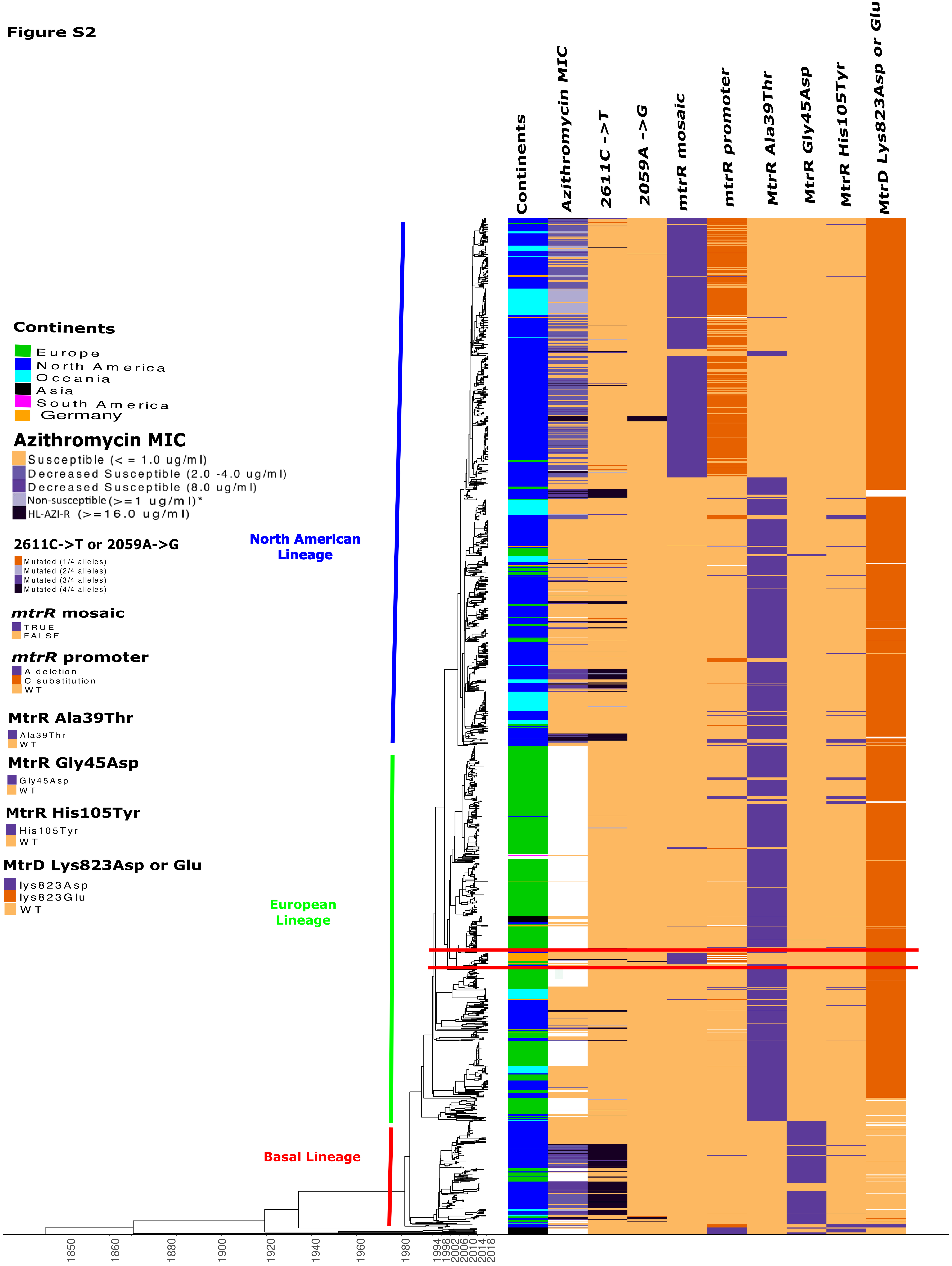
Continents
Azithromycin MIC
2611C ->T
2059A ->G
mtrR mosaic
mtrR promoter
MtrR Ala39Thr
MtrR Gly45Asp
MtrR His105Tyr
MtrD Lys823Asp or Glu
Continents
Europe
North America
Oceania
Asia
South America
Germany
Azithromycin MIC
Susceptible (< = 1.0 ug/ml)
Decreased Susceptible (2.0 -4.0 ug/ml)
Decreased Susceptible (8.0 ug/ml)
Non-susceptible (>=1 ug/ml)*
HL-AZI-R (>=16.0 ug/ml)
2611C->T or 2059A->G
Mutated (1/4 alleles)
Mutated (2/4 alleles)
Mutated (3/4 alleles)
Mutated (4/4 alleles)
mtrR mosaic
TRUE
FALSE
mtrR promoter
A deletion
C substitution
WT
MtrR Ala39Thr
Ala39Thr
WT
MtrR Gly45Asp
Gly45Asp
WT
MtrR His105Tyr
His105Tyr
WT
MtrD Lys823Asp or Glu
lys823Asp
lys823Glu
WT
North American Lineage
European Lineage
Basal Lineage
1850
1860
1880
1900
1920
1940
1960
1980
1994
1998
2002
2006
2010
2014
2018

#### Figure S3

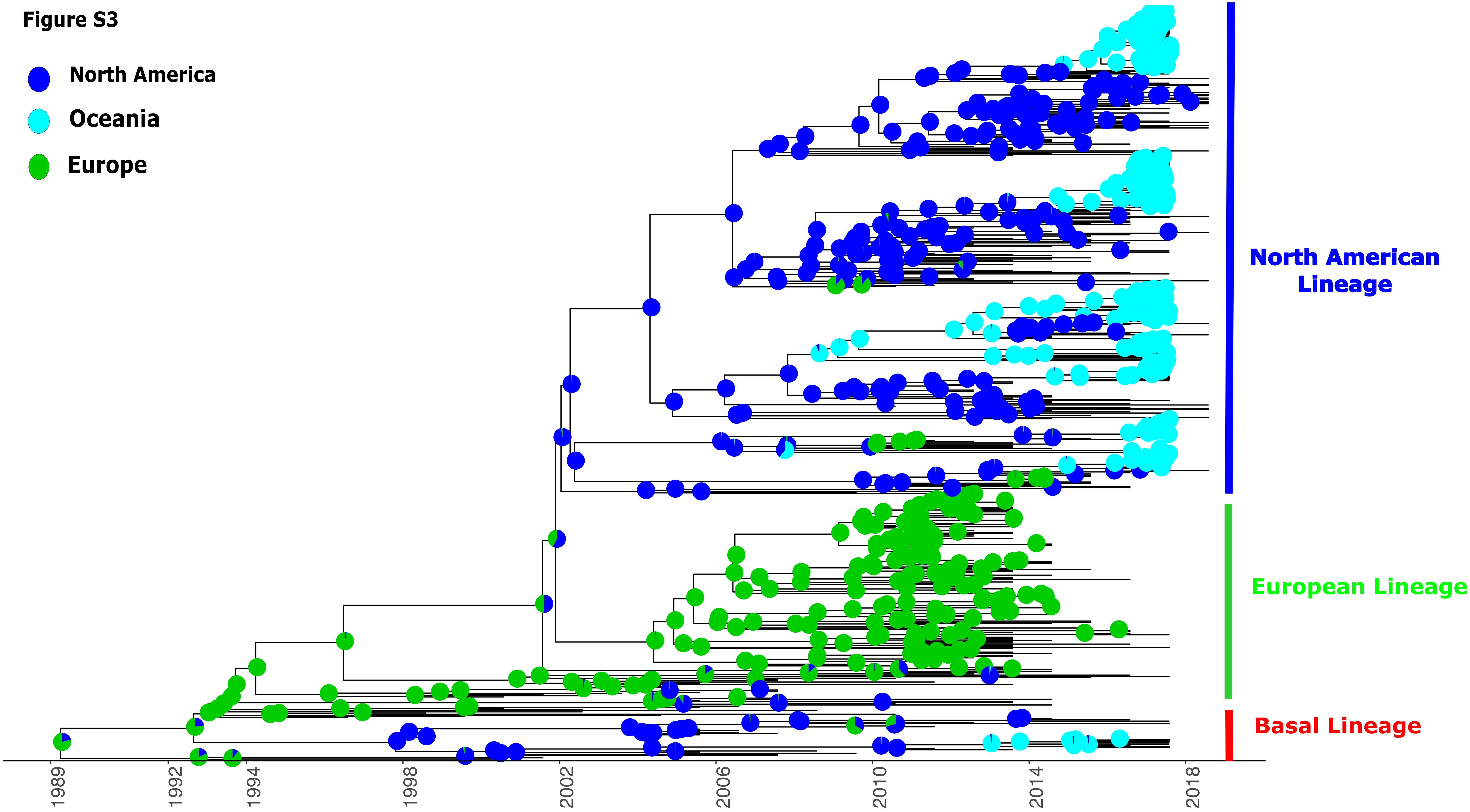
North America
Oceania
Europe
North American Lineage
European Lineage
Basal Lineage
1989
1992
1994
1998
2002
2006
2010
2014
2018

Figure S4

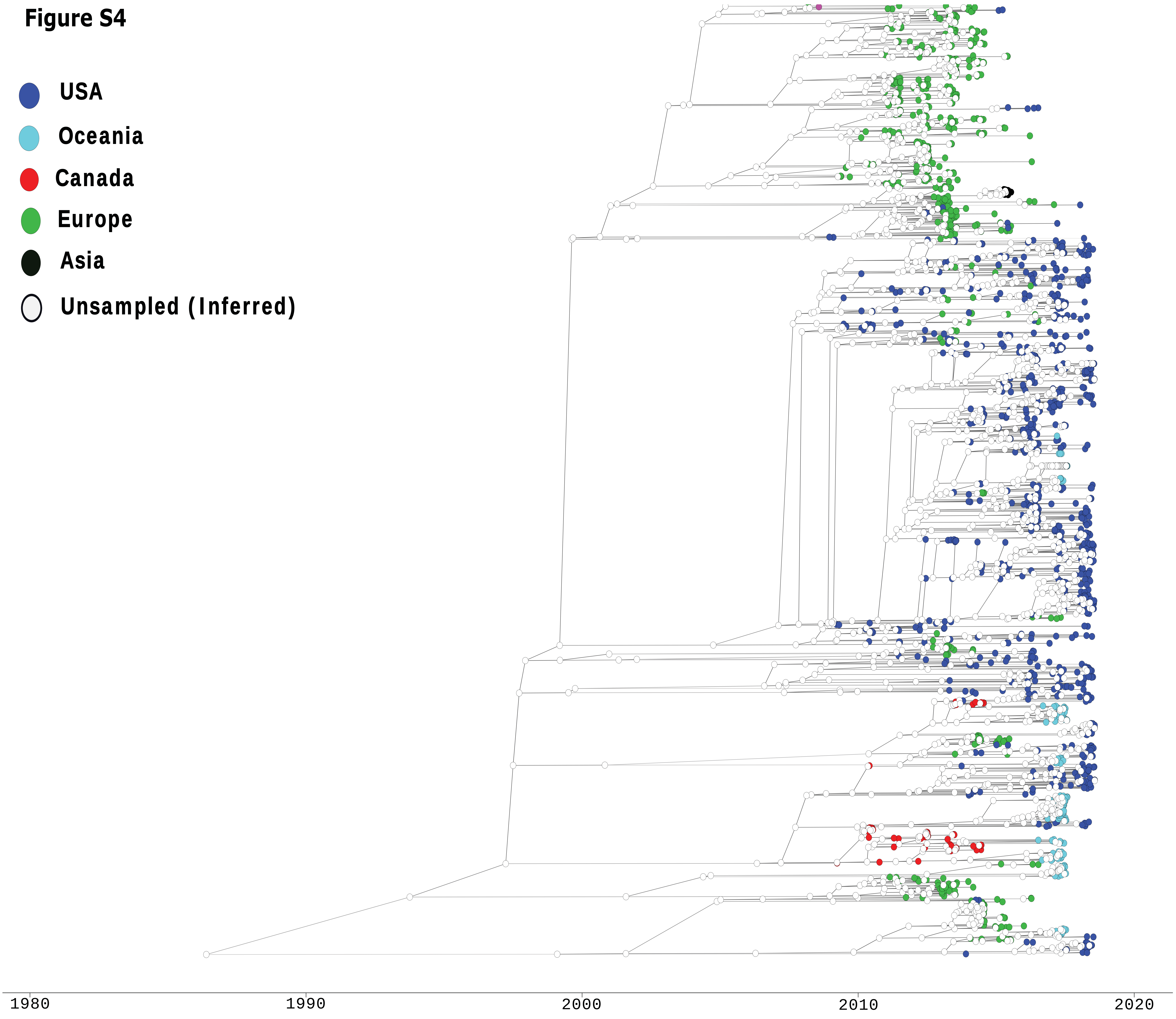
USA
Oceania
Canada
Europe
Asia
Unsampled (Inferred)
1980
1990
2000
2010
2020

**Figure S5**

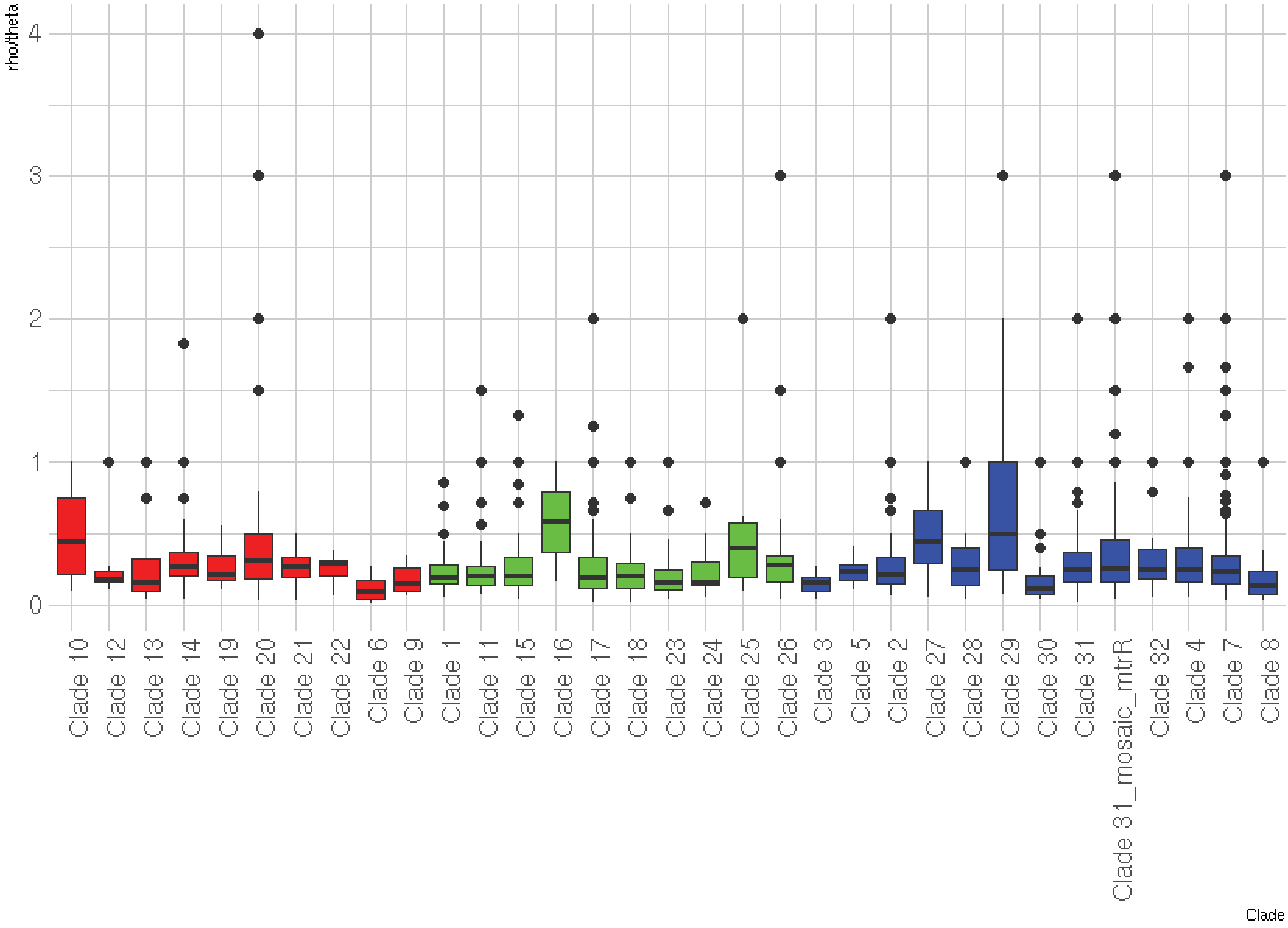
rho/theta
4
3
2
1
0
Clade 10
Clade 12
Clade 13
Clade 14
Clade 19
Clade 20
Clade 21
Clade 22
Clade 6
Clade 9
Clade 1
Clade 11
Clade 15
Clade 16
Clade 17
Clade 18
Clade 23
Clade 24
Clade 25
Clade 26
Clade 3
Clade 5
Clade 2
Clade 27
Clade 28
Clade 29
Clade 30
Clade 31
Clade 31_mosaic_mtrR
Clade 32
Clade 4
Clade 7
Clade 8
Clade

**Figure S6**

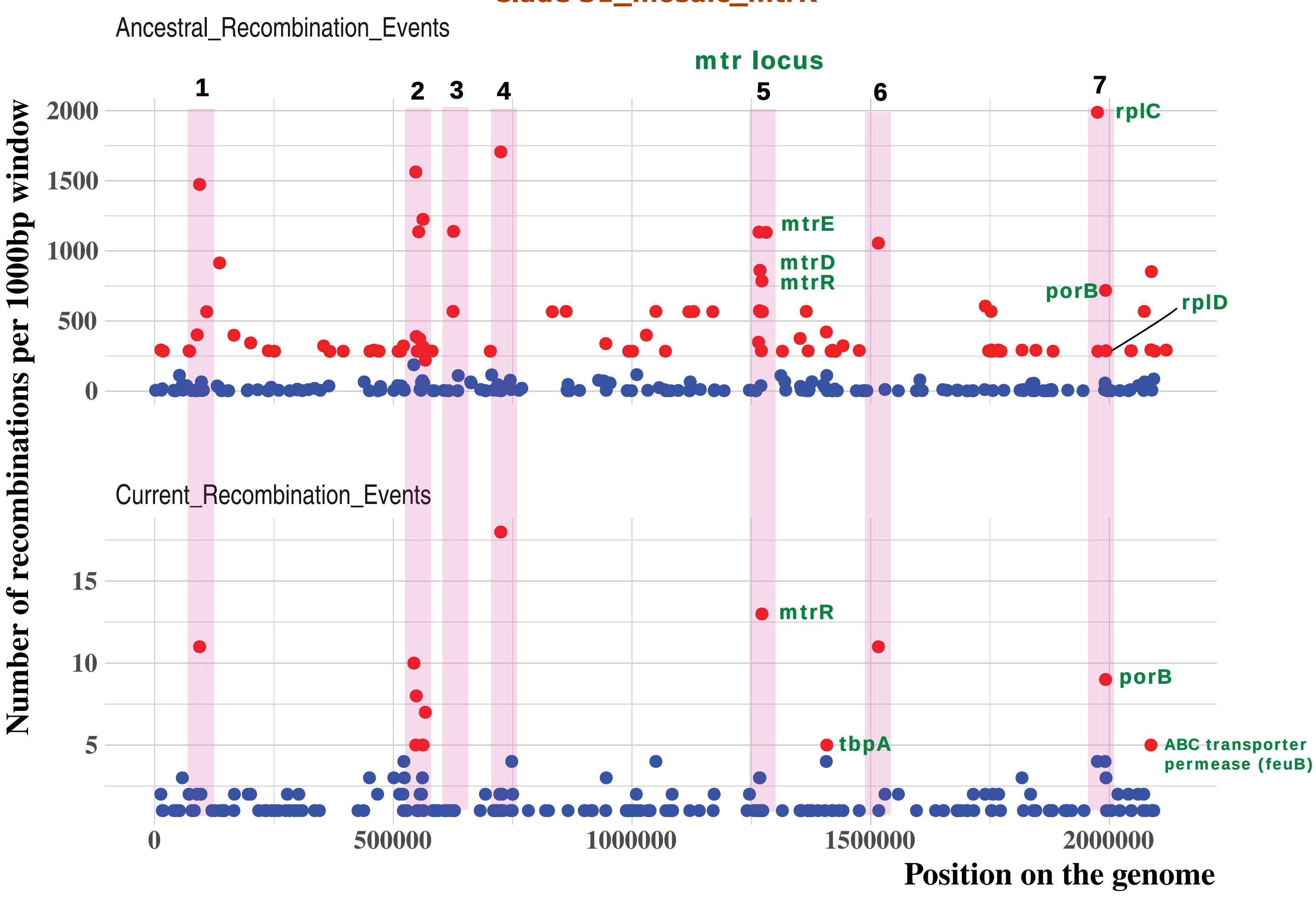


| | Genes in recombination hotspots* | |
|---|---|---|
| Hotspot regions | Gene Id | Annotation |
| 1 | WHOP_00089C | glycoside hydrolase family protein |
| 2 | WHOP_00586 | Uncharacterized conserved protein |
| 2 | WHOP_00590 | phage associated protein |
| 2 | WHOP_00591 | phage associated protein |
| 3 | WHOP_00597 | phage associated protein |
| 4 | WHOP_00762C | aspartyl/glutamyl-tRNA amidotransferase subunit B (gatB) |
| 4 | WHOP_00763C | Hypothetical membrane protein |
| 5 | WHOP_01388C | outer membrane efflux protein (mtrE) |
| 5 | WHOP_01398C | cbb3-type cytochrome c oxidase subunit II |
| 6 | WHOP_01655 | Uncharacterised protein |
| 7 | WHOP_02170 | 50S ribosomal protein L3 (rplC) |

*Genes in recombination hotspots with > 1000 events are only shown in this table

**Figure S7**

**Clade 4**

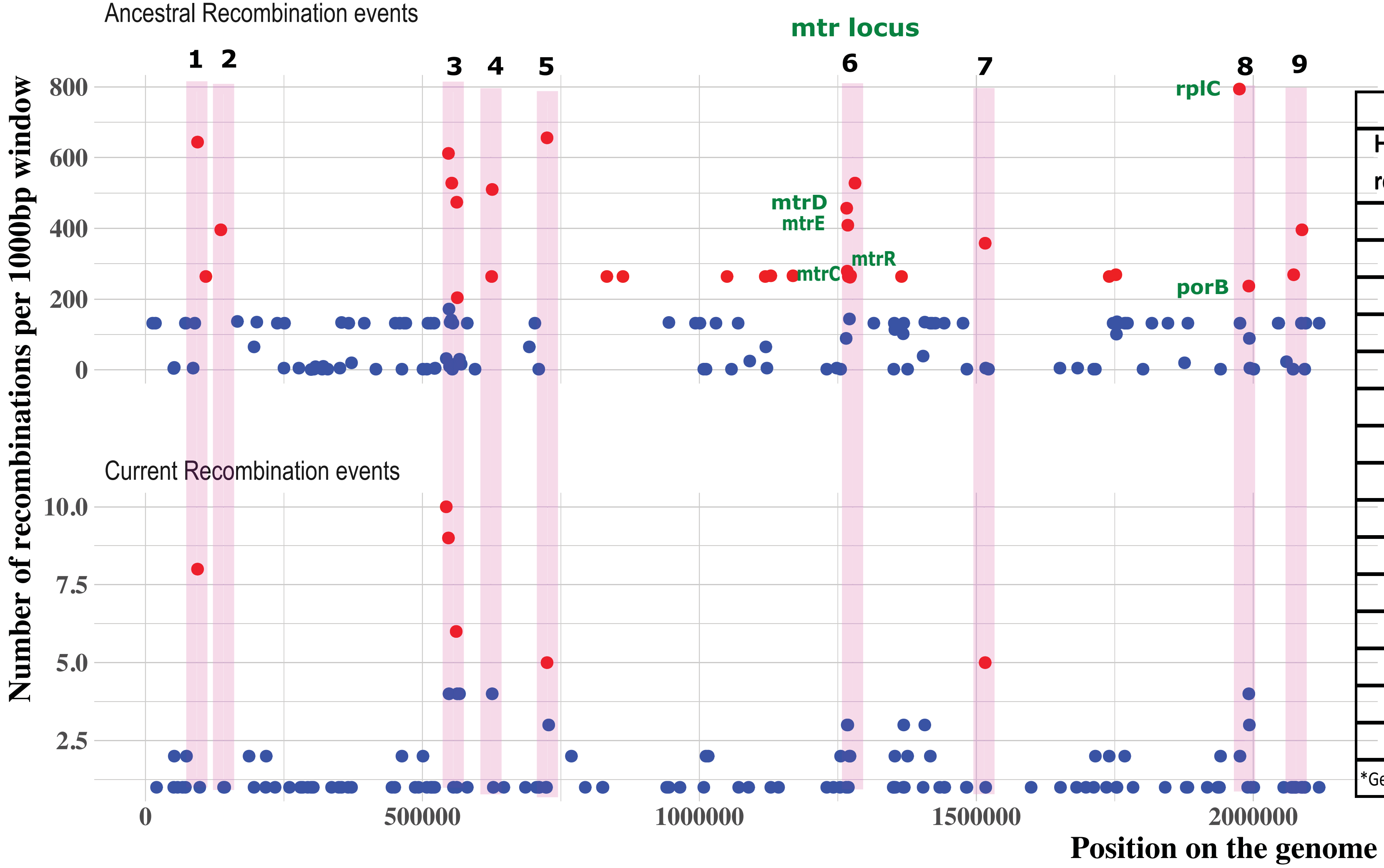


| | Genes in recombination hotspots* | |
|---|---|---|
| Hotspot regions | Gene Id | Annotation |
| 1 | WHOP_00089C | glycoside hydrolase family protein |
| 2 | WHOP_00131 | putative secretion protein (cvaA_1) |
| 3 | WHOP_00590 | phage associated protein |
| 3 | WHOP_00595 | phage associated protein |
| 3 | WHOP_00605 | phage associated protein |
| 4 | WHOP_00666C | Holliday junction resolvase-like protein (yqgF) |
| 4 | WHOP_00667C | Uncharacterized ACR |
| 4 | WHOP_00668C | CreA protein |
| 5 | WHOP_00763C | Hypothetical membrane protein |
| 6 | WHOP_01388C | outer membrane efflux protein (mtrE) |
| 6 | WHOP_01389C | antibiotic resistance efflux pump component (mtrD) |
| 6 | WHOP_01398C | cbb3-type cytochrome c oxidase subunit II |
| 7 | WHOP_01655 | Uncharacterised protein |
| 8 | WHOP_02170 | 50S ribosomal protein L3 (rplC) |
| 9 | WHOP_02299C | ABC transporter permease (feuB) |

*Genes in recombination hotspots estimated to have > 350 events are only shown in this table

**Figure S8**

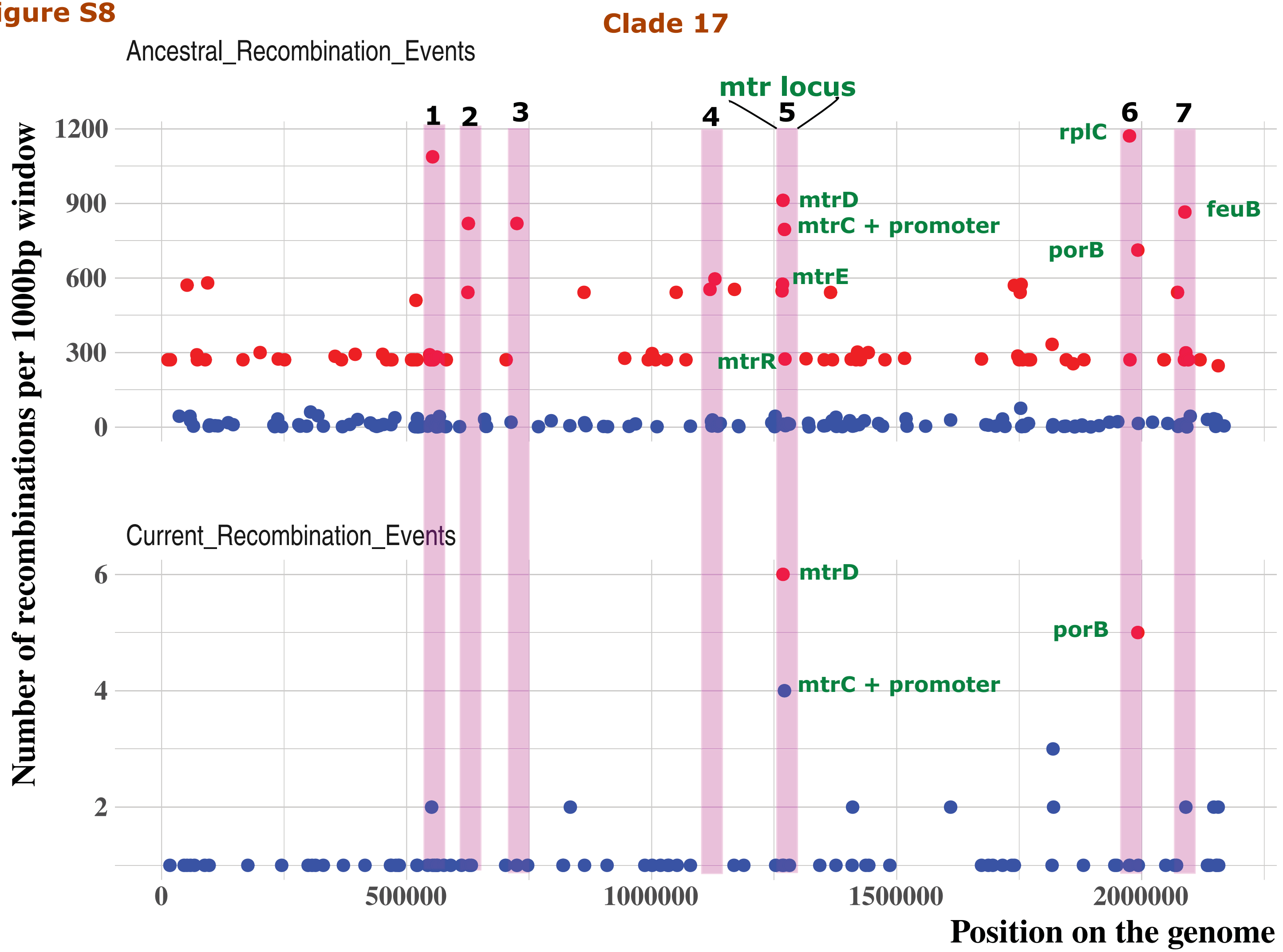


| | Genes in recombination hotspots* | |
|---|---|---|
| Hotspot regions | Gene Id | Annotation |
| 1 | WHOP_00595 | phage associated protein |
| 2 | WHOP_00668C | CreA protein |
| 2 | WHOP_00667C | Uncharacterized ACR |
| 2 | WHOP_00666C | Holliday junction resolvase-like protein (yqgF) |
| 3 | WHOP_00763C | Hypothetical membrane protein |
| 4 | WHOP_01229C | phage associated protein |
| 4 | WHOP_01216 | acyl-CoA hydrolase |
| 5 | WHOP_01389C | antibiotic resistance efflux pump component (mtrD) |
| 5 | WHOP_01390C | antibiotic resistance efflux pump component (mtrC) |
| 6 | WHOP_02170 | 50S ribosomal protein L3 (rplC) |
| 6 | WHOP_02204C | major outer membrane protein porin P.IB (porB) |
| 7 | WHOP_02299C | ABC transporter permease (feuB) |

*Genes in recombination hotspots estimated to have > 600 events are only shown in this table

Figure S9

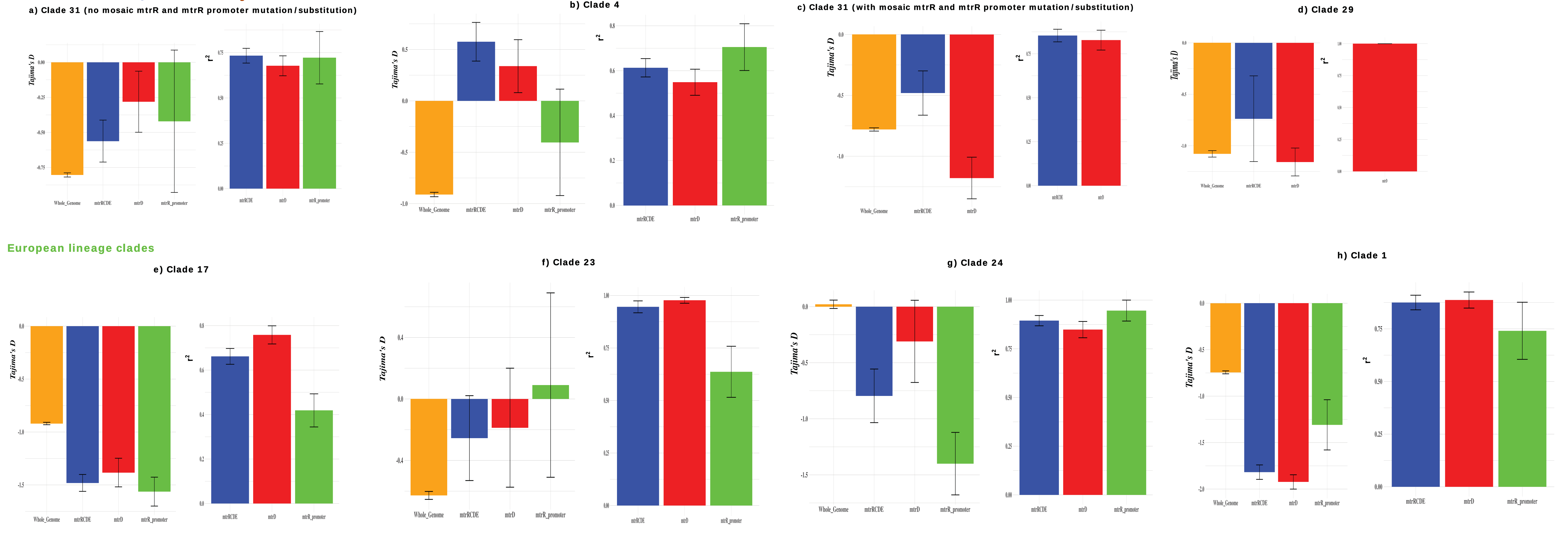

North American Lineage clades
a) Clade 31 (no mosaic mtrR and mtrR promoter mutation/substitution)
b) Clade 4
c) Clade 31 (with mosaic mtrR and mtrR promoter mutation/substitution)
d) Clade 29
European lineage clades
e) Clade 17
f) Clade 23
g) Clade 24
h) Clade 1
Tajima's D
r2
Whole_Genome
mtrRCDE
mtrD
mtrR_promoter

Figure S9(a)

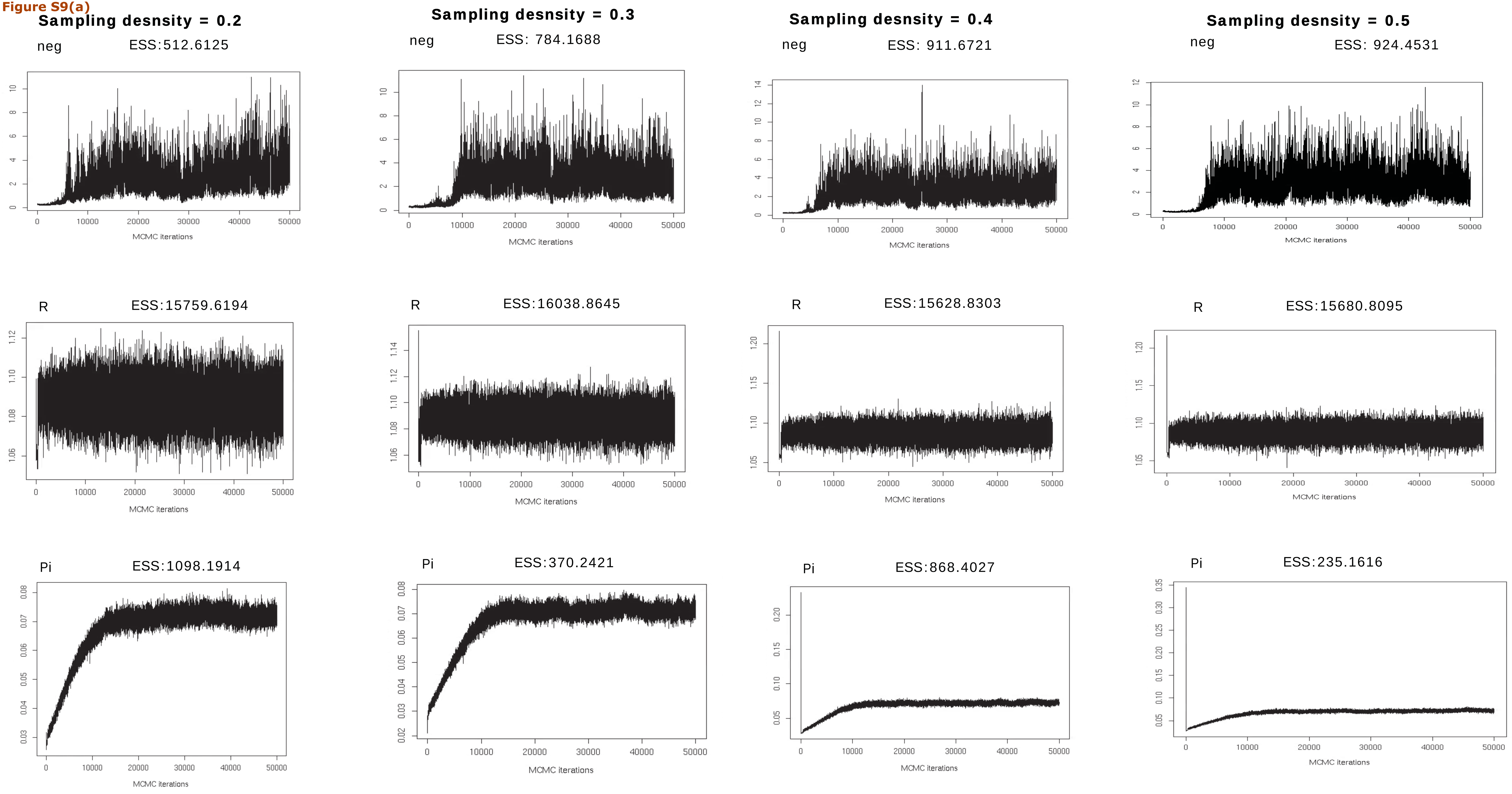
Sampling desnsity = 0.2
neg
ESS:512.6125
R
ESS:15759.6194
Pi
ESS:1098.1914
Sampling desnsity = 0.3
neg
ESS: 784.1688
R
ESS:16038.8645
Pi
ESS:370.2421
Sampling desnsity = 0.4
neg
ESS: 911.6721
R
ESS:15628.8303
Pi
ESS:868.4027
Sampling desnsity = 0.5
neg
ESS: 924.4531
R
ESS:15680.8095
Pi
ESS:235.1616
MCMC iterations

Figure S9(b)

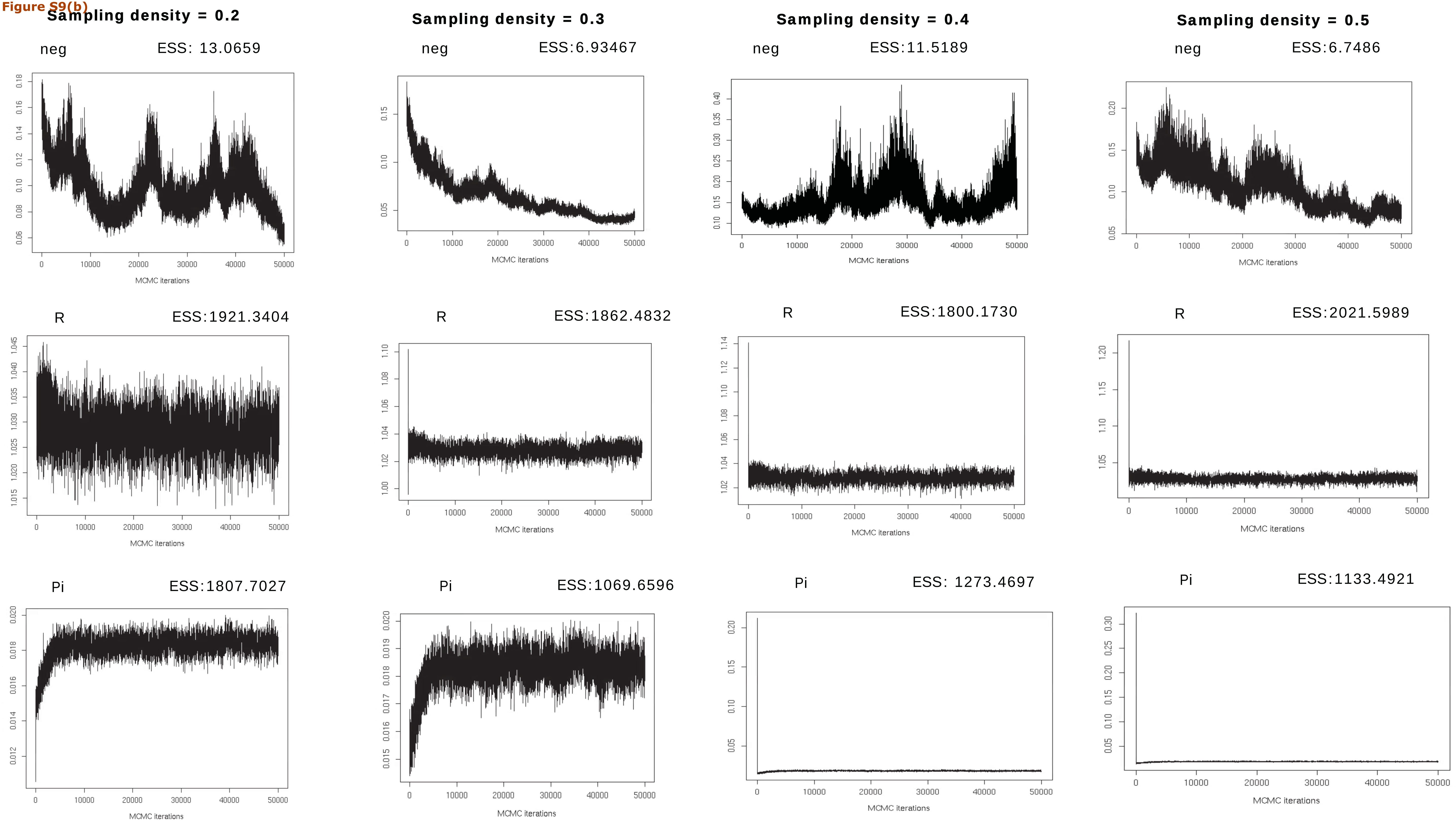
Sampling density = 0.2
neg
ESS: 13.0659
R
ESS:1921.3404
Pi
ESS:1807.7027
Sampling density = 0.3
neg
ESS:6.93467
R
ESS:1862.4832
Pi
ESS:1069.6596
Sampling density = 0.4
neg
ESS:11.5189
R
ESS:1800.1730
Pi
ESS: 1273.4697
Sampling density = 0.5
neg
ESS:6.7486
R
ESS:2021.5989
Pi
ESS:1133.4921
MCMC iterations
